# Cellular dialogues that enable self-organization of dynamic spatial patterns

**DOI:** 10.1101/717595

**Authors:** Yiteng Dang, Douwe Grundel, Hyun Youk

## Abstract

Cells form spatial patterns by coordinating their gene expressions. How a group of mesoscopic numbers (hundreds-to-thousands) of cells, without pre-defined morphogens and spatial organization, self-organizes spatial patterns remains incompletely understood. Of particular importance are dynamic spatial patterns - such as spiral waves that perpetually move and transmit information over macroscopic length-scales. We developed an open-source, expandable software that can simulate a field of cells communicating with any number of cell-secreted molecules in any manner. With it and a theory developed here, we identified all possible “cellular dialogues” - ways of communicating with two diffusing molecules - and core architectures underlying them that enable diverse, self-organized dynamic spatial patterns that we classified. The patterns form despite widely varying cellular response to the molecules, gene-expression noise, and spatial arrangement and motility of cells. Three-stage, “order-fluctuate-settle” process forms dynamic spatial patterns: cells form long-lived whirlpools of wavelets that, through chaos-like interactions, settle into a dynamic spatial pattern. These results provide a blueprint to help identify missing regulatory links for observed dynamic-pattern formations and in building synthetic tissues.

## INTRODUCTION

Highly organized spatial patterns can form when multiple cells, without pre-existing morphogen gradients, communicate with each other to coordinate their gene-expression levels [Gregor et al., 2010; Lubensky et al., 2011; Sgro et al., 2013; Idema et al., 2013, Manyukan et al., 2017; Jörg et al., 2019]. Understanding how a group of cells collectively organize (i.e., the group self-organizes) spatial patterns through cell-cell communications is crucial for understanding and engineering mammalian tissues [Javaherian et al., 2013]. Many synthetic and natural mammalian tissues are monolayers of genetically identical cells (e.g., epithelial sheets) whose gene-expression levels are initially spatially uncorrelated but become more correlated over time during development, resulting in specialized cell-types within tissues. This process often involves cell-cell communication [Menendez et al., 2010]. Recently, there has been a surge of interest in developing and using experimental methods for spatially arranging individual cells in a monolayer and then observing how such a heterogeneous tissue - composed of cells at differing locations having different gene-expression levels - develops over time [Javaherian et al., 2014]. Although quantitative models also now exist to complement such experiments, they are often tailored to specific tissues and signaling molecules so it is challenging to use them as a general framework that one can adapt to different gene-circuits, signaling molecules, and cell-types [Drasdo et al., 2007]. Partly as a result of this deficiency, we currently lack a general, quantitative mechanism for explaining how spatial patterns emerge in heterogeneous tissues made of realistic, mesoscopic numbers (hundreds to thousands) of cells, all without morphogen gradients (Figure 1A - top). Typically, to explain pattern formations, one uses reaction-diffusion equations and invokes the Turing instability - an instability that arises when amplifications of initially small chemical-concentration fluctuations in an uniformly spread field of chemicals lead to spatial patterns (Figure 1A - bottom) [Turing, 1952]. Although this mechanism is insightful, it applies to continuous fields of chemicals or cells (Figure 1A - bottom), and does not treat gene expression levels of individual cells when there are biologically realistic (mesoscopic) numbers of cells (Figure 1B - bottom). Recent work has extended models of biological pattern formation by including more general mechanisms that go beyond the Turing instability [Halatek & Frey, 2018], as well as individual cells and mechanical interactions [Lubensky et al., 2011; Recho et al., 2019]. However, much remains to be explored to obtain a general understanding of the relationship between the properties of cellular communication - the various ways in which the cells secrete and sense signaling molecules - and the gene expression patterns that emerge in a mesoscopic population of cells.

**Figure 1.**
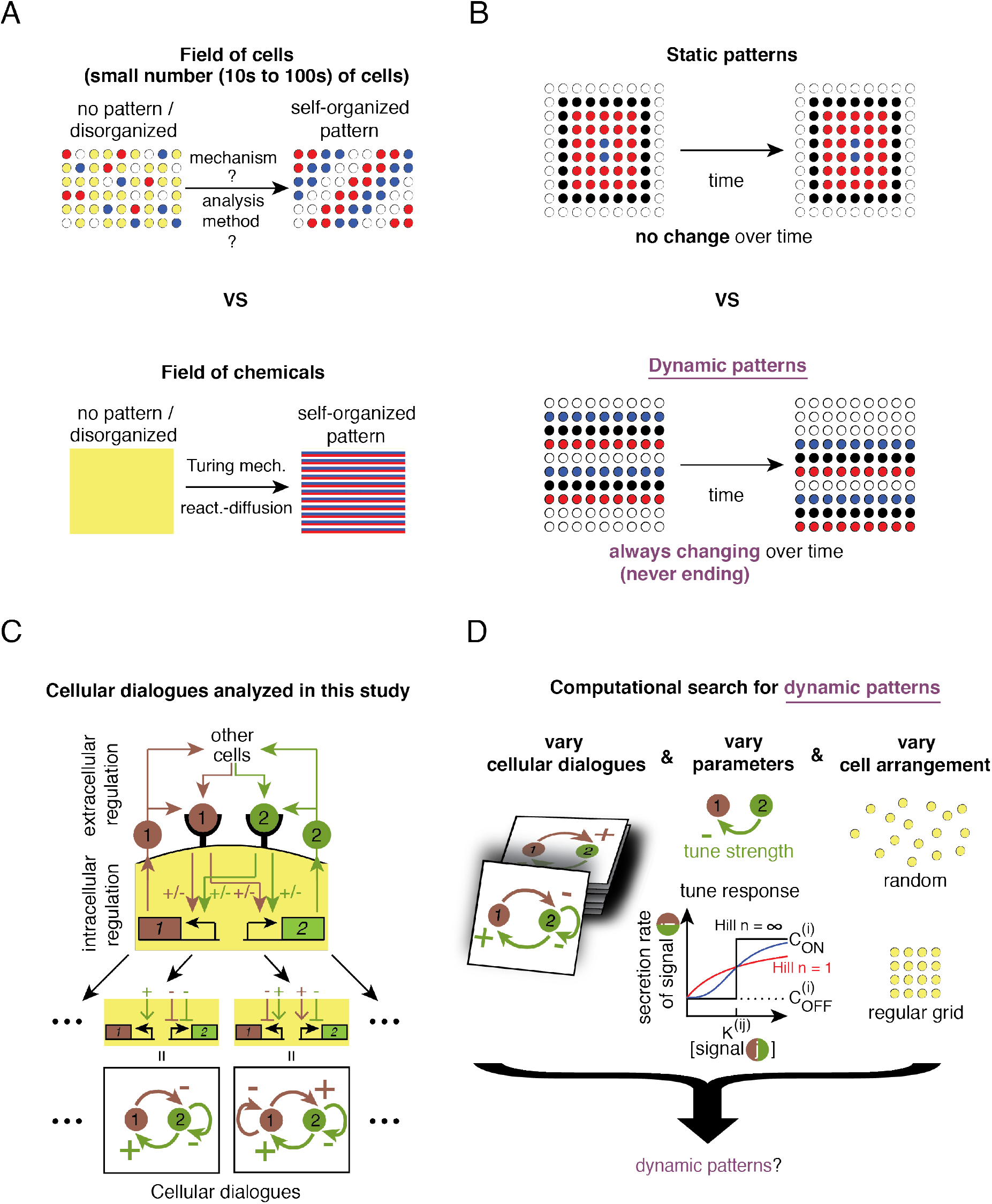
Computationally screening cellular dialogues to find ones that enable dynamic patterns to form. **(A)** Pattern formation by cells versus chemicals. (Top) Mechanisms by which an initially disordered field of a mesoscopic number of cells (~hundreds to thousands) (left panel) become more ordered through cell-cell communication (right panel) remain poorly understood, as is the method to analyze this complex self-organization dynamics. (Bottom) A field of chemicals or a continuum of cells (large number of tightly packed cells) initially having no pattern (left) can form a pattern (right) without pre-existing morphogens. This is usually modelled by reaction-diffusion equations and can be understood through the Turing mechanism. **(B)** Static versus dynamic patterns. (Top) Static patterns do not change over time. (Bottom) In dynamic patterns, a structure changes over time without ever stopping (e.g., shown here is a travelling wave). **C)** Schematic of cellular dialogues. Brown (molecule 1) and green (molecule 2) circles are ligands that bind to their cognate receptors on the cell membrane. Ligand-bound receptors trigger intracellular signal transductions that either positively or negatively regulate the production and secretion of molecules 1 and 2 (molecule 1 can self-promote or self-repress its own secretion while also regulating the secretion of molecule 2, and vice versa). Bottom row shows graphic representation of cellular dialogues. **(D)** Elements that we varied in simulations: cellular dialogues of all possible topologies, the values of the parameters for each cellular dialogue, and spatial arrangement of cells. Our study first begins with an infinite Hill coefficient (i.e., digital response to each of the two signaling molecules) and a regular lattice. After reporting the outcomes of these simulations, we report the result of relaxing these two constraints and well as other elements not depicted.

We sought to resolve this shortcoming by developing an open-source software that visually simulates spatial-patterning dynamics - a software that can be easily modified, is expandable with more ingredients, and will likely be useful for both research and educational purposes (https://github.com/YitengDang/MultiCellSim). We also developed algorithms for analyzing these simulations. With the software and analysis algorithms, we sought to reveal quantitative mechanisms by which mesoscopic numbers of cells can use their spatially heterogeneous gene-expression levels as a seed to form spatial patterns. In particular, we focused on dynamic patterns - patterns that constantly change over time without ever stopping such as oscillations and spiral waves [Sgro et al., 2013] - instead of static patterns that remain still after forming (Figure 1B). Through an exhaustive computational search, we discovered all the ways in which cells can communicate with just two diffusing molecules to form dynamic patterns, including those that have been experimentally observed. We found that just a few ways of communicating, which we refer to as “cellular dialogues”, can generate a large palette of complex, dynamic spatial patterns such as chaotic whirlpools of wavelets and travelling waves of various shapes and orientations. Viewing these simulations as exact numerical experiments, we devised an analytical (pen-and-paper) approach that recapitulates the simulations and used it to understand why only certain cellular dialogues can sustain dynamic spatial patterns. As we will show, we discovered that cells can form dynamic spatial patterns through a three-stage, “order-fluctuate-settle” process. Starting from a configuration in which there is no spatial correlation among cells’ gene-expression levels, we observed that cells rapidly become more spatially correlated over time, resulting in whirlpools of wavelets formed out of their correlated gene-expressions. This is then followed by a prolonged transient phase in which the whirlpools of wavelets annihilate each other while new ones are being formed. This results in the spatial correlation fluctuating over time in a seemingly chaotic manner that is reminiscent of deterministic chaos seen in logistic difference equation for population growth. Finally, as the fluctuations settle - due to the wavelets settling down - a dynamic spatial pattern such as a travelling wave emerges. This represents a symmetry-breaking transition in which the dynamic pattern (e.g., traveling wave) chooses a direction to travel in, even though there are no earlier indications that any one direction would be preferred over any other direction. We show that self-organized dynamic patterns survive wide variations in gene-expression noise, cells’ responses to the sensed molecules, spatial arrangements of cells (even when cells are randomly scattered), and diffusive motions of cells (i.e., each cell randomly moving). For each of these elements, we quantified how much perturbation is sufficient to disrupt the formation of dynamic spatial patterns. As a theoretical study, our study focuses on how cells *can* form dynamic spatial patterns, rather than how certain cells *do* form them. Despite not tailoring our study to a particular multicellular system, our computational search yielded cellular dialogues that are known to generate dynamic spatial patterns in certain multicellular systems. Our paper ends by discussing these examples as well as suggesting how one can expand our work, including the open-source software, to computationally identify cellular dialogues that may produce dynamic spatial patterns that are observed in multicellular systems but whose underlying cellular dialogue remains unknown.

## RESULTS

### Computational search for cellular dialogues that enable self-organized patterns

We built an open-source visualization software that simulates all possible ways in which cells can communicate - which we call “cellular dialogues” - by secreting, sensing, and responding to two diffusing molecules (Figure 1C). Our simulations combine reaction-diffusion equations these describe the concentrations of the two diffusing molecules – and a cellular automaton this describes the cells’ gene-expression levels that are set by the concentrations of the two molecules. We represent a cellular dialogue as a network diagram that consists of two nodes (one for each molecule) joined by signed arrows, which can be positive (activating) or negative (repressing). A signed arrow denotes how the sensing of one molecule, represented by the node on which the arrow begins, increases (for a positive arrow) or decreases (for a negative arrow) the sensing cell’s secretion rate of a molecule that is represented by the node on which the arrow ends (the same or the other molecule) (Figure 1C). We assume that both molecules diffuse on a faster timescale than the cells can respond to them - the two molecules “rapidly” diffuse - as is the case in many multicellular systems [Heemskerk et al., 2019]. We first considered cells that digitally respond to each molecule: a cell secretes “molecule-*i*” at either a low rate (“OFF” state for molecule-*i*) or a high rate (“ON” state for molecule-*i*). If molecule-*j* activates (represses) molecule-*i*, then a cell becomes ON (OFF) for molecule-*i* if and only if it senses a concentration of molecule-*j* that is *above* a set threshold concentration. We will later relax this assumption and consider cells with continuous response to the sensed molecules. We first considered these digital cells for two reasons. First, experimental studies have shown that signal transduction pathways such as MAPK or other phospho-relay cascades that are triggered by ligand-bound receptors and control gene-expressions downstream, as in our digital cells (Figure 1C), can have an effective Hill coefficient with a value of 4 or more (e.g., as high as 32 [Trunnell et al., 2011]). An effective Hill coefficient characterizes the “sharpness” of cell’s response to a ligand [Ferrell & Ha, 2014; Plotnikov et al., 2011; Trunnell et al. 2011]. Such high numbers are due to multiple molecular parts amplifying each other’s effects in combination. A digital (ON/OFF) response models such high-valued Hill coefficients. Second reason for considering digital cells is that a digital response simplifies the mathematics used for describing the response, while retaining its main qualitative features, even when the actual Hill coefficient of the system being modeled is relatively low [Alon, 2007]. Finally, the digital cells also have a reporter gene for each molecule, called genes “1” and “2”, that are also either ON or OFF to reflect the secretion state of its corresponding molecule (Figure 1C - brown and green boxes). In the simulations, we assigned a distinct color to each of the four states, which are (ON for gene-1, ON for gene-2), (ON, OFF), (OFF, ON), and (OFF, OFF).

We began each simulation by randomly assigning the four gene-expression states (i.e., four colors) to each cell so that the gene expression levels (four colors) were spatially uncorrelated. Thus, the field of cells initially did not exhibit any spatial organization. We quantitatively verified this with a “spatial index” metric which is a weighed spatial autocorrelation function that is zero when cells are completely, spatially disorganized and increases towards one as cells become more spatially organized (see Section S1.3). We then observed how each cell’s state (i.e., four colors) changed over time to determine whether a spatial pattern formed and, if so, what type of a pattern that was. For each cellular dialogue, we fixed the values of all parameters (e.g., threshold concentrations, secretion rates for each molecule), and then ran thousands of simulations that each started with a different, randomly-chosen disordered configuration (see Section S2). We ran the simulation for a wide range of parameter values for every possible cellular dialogue (Figure 1D and see Section S2). We first performed such a computational search with immobile digital cells that were placed on a regularly spaced lattice. We will first describe these results in the next sections. Afterwards, we will describe how these results change when we relax the constraints – by randomly displacing cells so that they no long form a regular lattice, having each cell randomly and constantly move, allowing the Hill coefficient to be any finite value (i.e., analogue instead of digital response), and enabling stochastic gene expressions (Figure 1D).

### Cellular dialogues enable self-organization of wide array of dynamic patterns

The computational search revealed a surprisingly wide variety of dynamic patterns, from never-ending and temporally repeating travelling waves (Figure 2A and Video S1) to temporally non-repeating, complex patterns consisting of whirlpools of wavelets that perpetually evolved over time in a chaos-like manner (Figure 2B). All patterns self-organized from completely disorganized fields of cells by their ON/OFF-states becoming more spatially correlated over time (Figure 2A-B). The time taken to self-organize widely varied and depended on the type of pattern formed. For example, if we assume that a gene-expression change such as an ON-cell becoming an OFF-cell takes one minute – this is one time-step of the simulation and every cell synchronously changes their ON/OFF states – then horizontal waves could take nearly six hours to form (Figure 2A) whereas the constantly changing, complex whirlpool of wavelets would not show any signs of settling into any pattern that cyclically repeats itself even after a week or longer (i.e., until we terminate the simulation) (Figure 2B). Since the simulations are deterministic for now – we will later add gene-expression noise – we can conclude that once a simulation reproduces a spatial configuration that it had before, the cell population has formed a dynamic pattern that periodically repeats itself forever.

**Figure 2.**
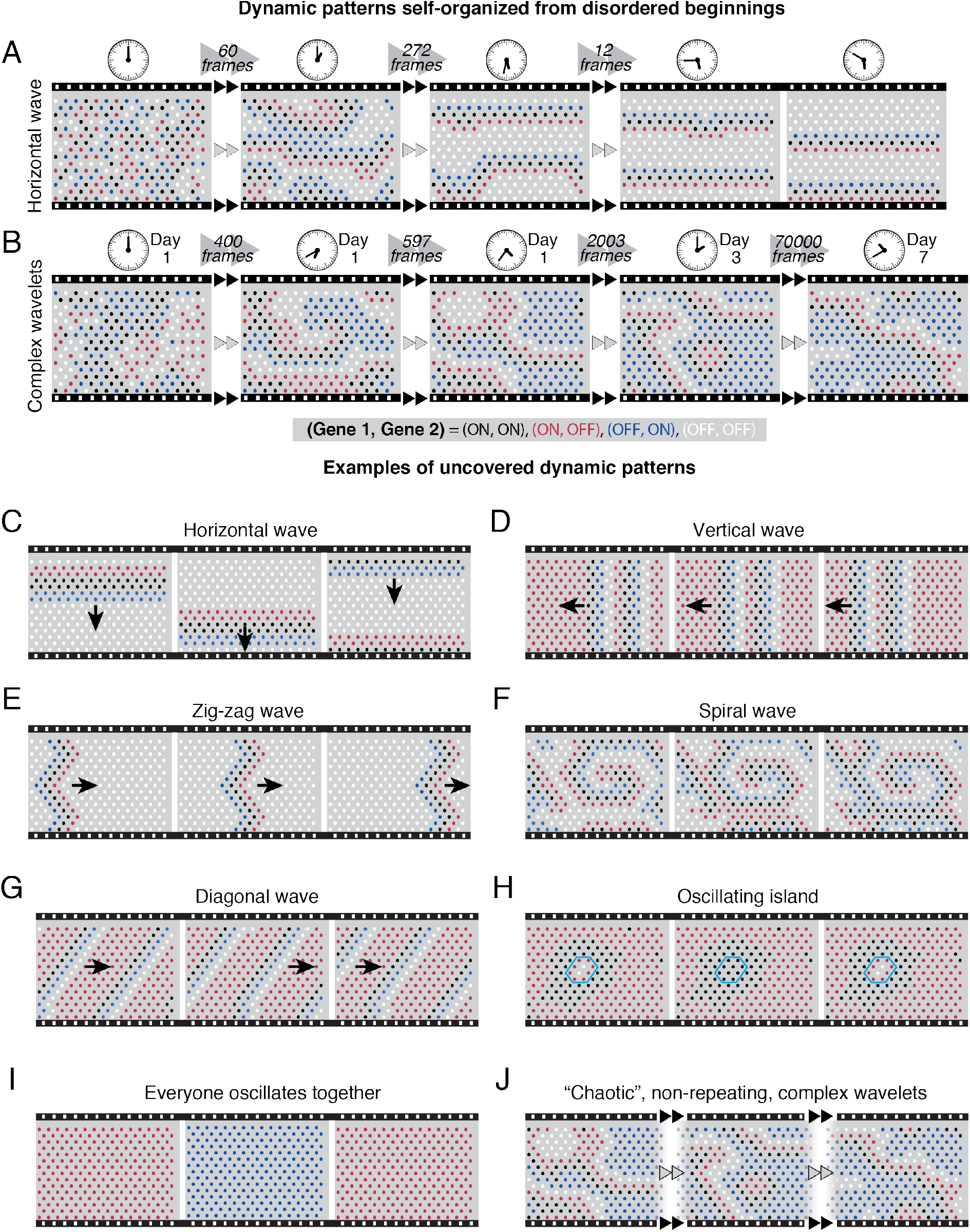
Examples of self-organized dynamic patterns found through computational screening. In all the figures shown here, a cell (drawn as a circle) can have four colors. Each color represents a distinct gene-expression state, (gene 1 = ON/OFF, gene 2 = ON/OFF): Black means (ON, ON), red means (ON, OFF), blue means (OFF, ON), and white means (OFF, OFF). In all the simulations, a field of cells starts with a completely spatial disordered configuration - there is no correlation between neighboring cells’ gene-expression states - as exemplified by the leftmost picture shown in (A). **(A)** Travelling wave of horizontal bands. Snapshots of the formation process shown at different stages of a simulation. Assuming that one timestep in the simulation takes one minute, the clocks show time passed from noon (beginning of the simulation). **(B)** Complex pool of multiple wavelets formed, starting with a spatially disorganized field of cells. Snapshots at different stages of the simulation are shown. Assuming that one timestep represents one minute, the clock and the days elapsed indicate at which timesteps in the simulation the snapshots are taken. **(C-J)** Each filmstrip shows three non-contiguous snapshots of a moving, dynamic pattern that formed, starting from a spatially disorganized configuration (not shown, see examples in the first snapshots in A). Where shown, the arrows represent the direction of travel. The dynamic patterns are: (C) a single travelling horizontal band, (D) travelling vertical bands, (E) a travelling zig-zag band, (F) a spiral wave, (G) travelling diagonal bands, (H) a small island of cells (enclosed in the blue hexagon) oscillating over time while all cells outside the island remain static, (I) every cell oscillates between red and blue with period 2, and (J) seemingly chaotic, never-ending dynamics in which multiple wavelets form and meet and annihilate each other, with the pool of wavelets constantly evolving and never repeating the same configuration throughout the simulation.

The dynamic patterns that we uncovered differed in their shape, complexity, and dynamics (Figures. 2C-J, Videos S1-S4, Section S3 and Section 4.1). Among these, the most prominent and striking were rectilinear travelling waves and spiral waves, both of which have high degrees of spatial order (Figures 2C-F). In the case of travelling waves – which can be oriented horizontally, vertically, or diagonally (Figures 2C-D, and 2G) and have a straight or bent shape (Figures 2D-E) – a rigid shape moves across space over time. Since the simulations were deterministic and the system had periodic boundary conditions, we concluded that a spatial configuration recurring over time during a simulation meant that the simulation would periodically and forever repeat the same dynamics from then on, indicating that a dynamic pattern had formed. In the case of travelling waves, this meant that the waves indefinitely repeated themselves, disappearing at one edge of the field and then appearing at the opposite end. This behavior also applies to patterns that do not propagate over space, but rather, oscillate in time. In some cases, such oscillations were limited to a few cells that formed an island (Figure 2H) whereas in others, every cell in the field oscillated together (Figure 2I). In particular, some islands of cells could collectively oscillate with periods that were higher than four time-steps (Figure 2H), indicating that individual cells were not just cycling through a fixed set of four distinct gene-expression states (note that a cell has a total of four distinct (ON/OFF, ON/OFF) states). If collective oscillations were to arise from synchronizations of individually oscillating cells, then we would expect a period of four timesteps or less. This indicates that the collective oscillations with a period greater than four timesteps form through a more intricate mechanism. Finally, some cellular dialogues yielded temporally non-repeating, complex patterns consisting of whirlpools of wavelets that evolved over time in a seemingly chaotic manner (Figure 2J) which, in many cases, transiently existed for tens of thousands of timesteps (i.e., week or longer if one minute represents one time-step) while the cells were on their way to forming a repeating, well-defined dynamic patterns such as horizontal waves.

### Common structural elements in cellular dialogues that generate dynamic patterns

The wide array of dynamic patterns that we observed fall into two categories (Figure 3A): (1) dynamic *temporal* patterns, in which cells periodically oscillate over time but do not propagate information over space (e.g., Figures 2H-I), and (2) dynamic *spatial* patterns, in which cells propagate information over space in the form of a well-defined shape (e.g., a wave front) that moves from one part of the field to another, often from one edge to the other edge of the field (e.g., Figures 2C-F). There are 44 distinct cellular dialogues in total (Section S1.2), that we could group into three categories: (1) those that cannot form any dynamic patterns, (2) those that can form only dynamic temporal patterns, and (3) those that can form both dynamic spatial patterns and dynamic temporal patterns. To categorize them, we developed a method to deduce, for each cellular dialogue, all possible ways that a cell’s state (ON/OFF, ON/OFF) can change over time. Concretely, we constructed a directed graph for each cellular dialogue (see Section S4) which has four nodes (one for each state) that are connected by edges with directions that represent the allowed transitions between the nodes. We deduced how some of the directed edges become inaccessible while others become accessible as we change the cellular dialogue’s parameter values. Then, following the directed edges from node to node yields all possible ways that a cell’s gene-expression can change over time. By looking for graphs that contained cyclic paths, we identified cellular dialogues and ranges of their parameter values (e.g., threshold concentrations and maximal secretion rates for each molecule, cell-cell spacing) that can potentially sustain dynamic patterns if they were to form. Since self-organization of dynamic patterns can only occur for parameter values that can sustain dynamic patterns in the first place, we only had to check these values in simulations to see if they led to dynamic patterns. This method thus vastly reduced the range of parameter values that we had to screen. For each cellular dialogue, we generated a large set of random parameters and ran many simulations (10,000 parameter sets with 10 simulations each), each starting with a different, randomly selected and disordered gene-expression pattern in the field. We checked whether each of these simulations yielded a dynamic pattern using automated methods (Section S2).

**Figure 3.**
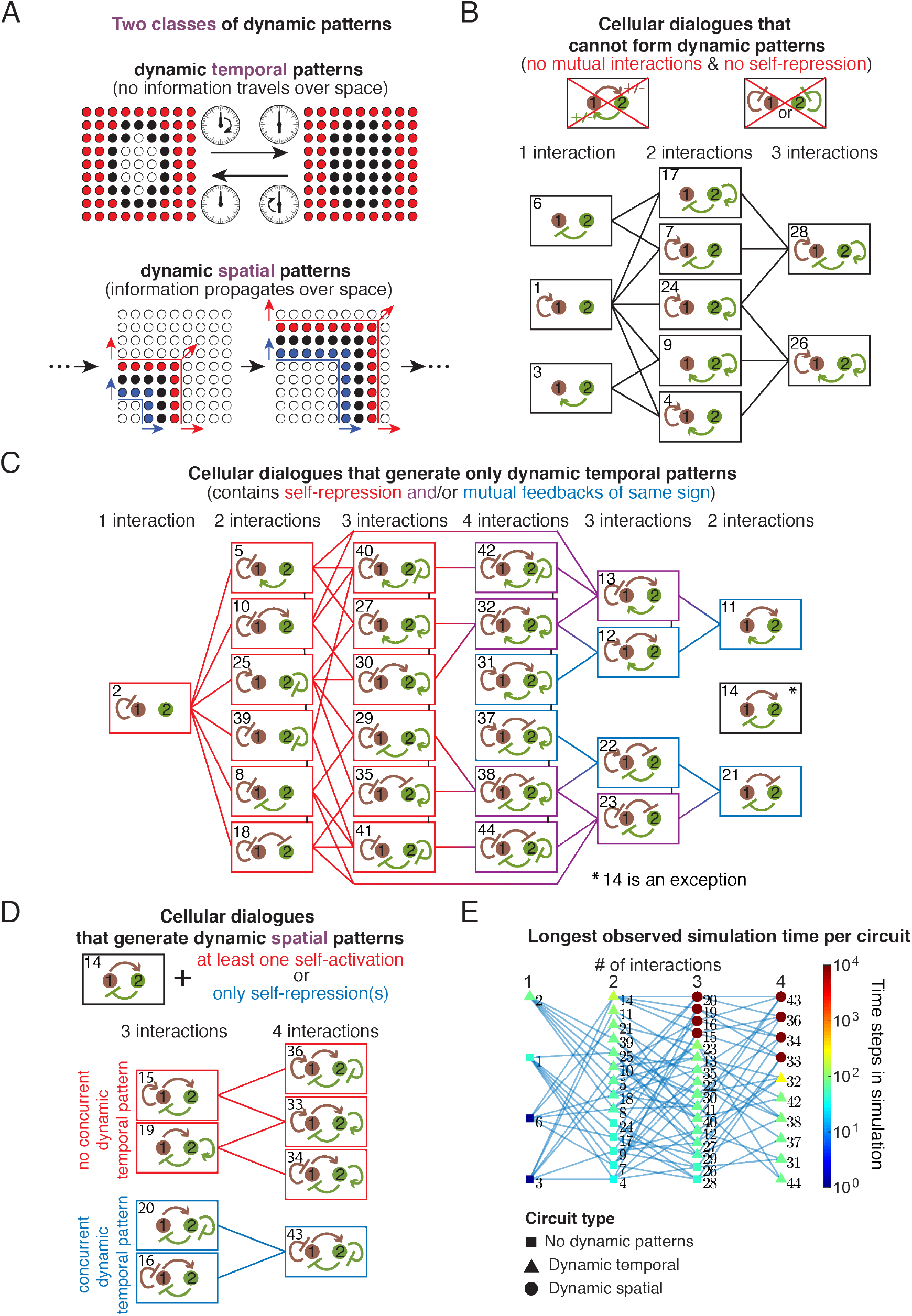
Computational search revealed tree structures that group cellular dialogues based on their ability to generate either static patterns, dynamic temporal patterns, or dynamic spatial patterns. **(A)** Two classes of dynamic patterns. (Top): Dynamic temporal patterns repeat themselves over time without transmitting information across space. (Bottom): Dynamic spatial patterns involve cells that transmit information over space through a coherent structure that moves across the field. **(B-D)** Tree diagrams show a full classification of all 44 unique, non-trivial cellular dialogues into three distinct classes (Section S1.2). In each tree diagram, a cellular dialogue is a leaf (box) that is joined by branches to other cellular dialogues. As one moves from one leaf to the next, an edge is either removed or added to the cellular dialogue. (B) Tree diagram showing all cellular dialogues that cannot generate any dynamic patterns. All cellular dialogues here lack mutual interactions and self-repressions. (C) Tree diagram showing all cellular dialogues that can generate dynamic temporal patterns but not dynamic spatial patterns. These all have either a self-repression (red boxes), a mutual interaction of the same sign (blue boxes), or both (purple boxes). Cellular dialogue 14 is an exception - it has mutual interactions of different signs and no self-interactions. (D) Tree diagram showing all cellular dialogues that can generate dynamic spatial patterns - these can all also generate dynamic temporal patterns. These are all generated by adding at least one additional self-interaction to cellular dialogue 14. Cellular dialogues in the five red boxes have at least one positive feedback loop, and can generate non-oscillatory dynamic spatial patterns (e.g., traveling waves). Cellular dialogues in the blue boxes have only negative self-interactions and produce dynamic spatial patterns but always with a concurrent dynamic temporal pattern (e.g. a traveling wave where the cells oscillate simultaneously) (see Fig. S1 for examples). **(E)** The maximum observed simulation time is a metric that naturally separates the three classes of cellular dialogues (B-D) (see Fig. S2 for other metrics). A node represents a cellular dialogue and the node’s shape represents the type of cellular dialogue (one of the three (B-D)). A node’s color indicates the longest observed simulation time among a large set of simulations that were performed with different parameters.

We discovered that cellular dialogues, when grouped into the three categories mentioned above, form distinct tree structures (Figures 3B-D) in which a node denotes a particular cellular dialogue and an edge connects two nodes if one node (cellular dialogue) comes from the other node (another cellular dialogue) by adding or removing one interaction. The fact that tree structures emerged, which link the different cellular dialogues together if they form the same type of patterns, suggests that there may be common elements in the cellular dialogues that belong to the same tree. Indeed, we found that all ten cellular dialogues (Figure 3B) that can only generate static configurations, and no dynamic patterns at all, consist of two molecules that do not mutually regulate each other and also do not have any self-repressions. We also found that twenty-six cellular dialogues can produce dynamic temporal patterns but not dynamic spatial patterns (Figure 3C). Their common feature is that they all contain a self-repression and/or a mutual feedback of the same sign (i.e., both molecules either activate or repress each other’s production). The sole exception to this rule, within this family of cellular dialogues, is cellular dialogue 14 (Figure 3C). As a result, it is disconnected from the tree of cellular dialogues that form dynamic temporal patterns (Figure 3C). Cellular dialogue 14 consists of an activator-inhibitor pair, whereby one molecule promotes the production of the second molecule, which in turn represses the production of the first molecule. Here, neither molecule regulates its own production. Cellular dialogue 14 is also special because we discovered that all cellular dialogues that one can obtain from it by adding one or more self-interactions - there are eight such cellular dialogues in total - can yield dynamic spatial patterns, in addition to dynamic temporal patterns (Figure 3D). We could further divide these eight cellular dialogues into two classes: ones that contain only self-repressions (Figure 3D - blue boxes) and ones that contain at least one self-activation (Figure 3D - red boxes). The three cellular dialogues that contain only self-repressions produce dynamic spatial patterns in which the moving shape periodically changes its gene-expression-composition (Figure S1 and Video S3) whereas the five cellular dialogues that contain at least one self-activation yield dynamic spatial patterns such as travelling waves (Figures 2C-G) in which the pattern moves across the field of cells without changing in shape or composition.

### Grouping cellular dialogues based on how fast they form patterns is equivalent to grouping them based on their shared structural elements

So far, we analyzed the patterns formed to determine the category (tree) that a cellular dialogue belongs to (Figures 3B-D). Next, we asked whether we could also obtain this classification from any statistical measures based on our large set of simulations (Figure 3E and Figure S2). Surprisingly, we discovered that if we analyze the typical times or the longest time that a cellular dialogue takes to form a pattern (static configuration or a dynamic pattern), and then group the cellular dialogues based on those times, then we would identify the same three categories of cellular dialogues (Figure 3E). Specifically, we found that all eight cellular dialogues that can form dynamic spatial patterns stood out as taking the longest times to form patterns compared to the other cellular dialogues, by at least about 100-fold longer durations (Figure 3E - circles). As we will later discuss, we found that these long self-organization times (~1 week if one time-step represents one minute) are due to a chaos-like process that is intrinsic to the pattern formation process. We found that all cellular dialogues that cannot form dynamic spatial patterns but do form dynamic temporal patterns take less times to form patterns, by at least a 100-fold less, than the ones that form dynamic spatial patterns (Figure 3E - triangles). Finally, we discovered that the cellular dialogues that cannot form any dynamic patterns and thus only form static configurations – some of which are highly organized patterns – require the least amounts of time to form these configurations (Figure 3E - squares).

### Analytical framework explains how cells collectively sustain dynamic spatial patterns

Although we have identified the five cellular dialogues that enable cells to form and sustain dynamic spatial patterns, we have not yet explained why these cellular dialogues enable cells to sustain the dynamic spatial patterns after having formed them. To find the answer, we developed a theory that does not use simulations and still correctly predicts *when* dynamic spatial patterns occur and explains *how* the cells sustain them (Figures 4A-C). The key idea behind this analytical approach is that a common structure exists at the core of a wide variety of dynamic spatial patterns, from the complex whirlpools of wavelets to spiral waves: we realized that one can build diverse dynamic spatial patterns by gluing together multiple rectilinear waves (i.e., horizontal, vertical, and bent waves - Figure 2) just as gluing together straight, tangent lines can build smooth curves. Thus, if we can understand how cells can sustain rectilinear waves, we can piece them together to understand the more complex shapes that are built out of them. Each rectilinear wave has six distinct layers of gene-expression states (Figure 4A). Three of the layers – “front”, “middle”, and “back” (Figure 4A - red, black, blue cells) – constitute the wave itself and continuously move forward while the other three layers – “exterior front”, “exterior” and “exterior back” – consist of all the other cells. After one timestep, each layer adopts the identity of the layer just behind it (e.g., the exterior-front layer, which is just in front of the front layer, becomes the front layer) (Figure 4C). This must occur at every timestep in order for the wave to continuously propagate, meaning that the concentrations of the two molecules within each layer must coordinately change so that the layers can synchronously move forward. We developed a method to estimate the concentrations of the molecules in each layer (Figure 4B and Section S5). Using this method, we derived six mathematical inequalities, one for each layer, that must all be satisfied in order for the concentrations of the two molecules to coordinately change, which would ensure that a rectilinear wave can propagate (Figure 4C). The inequalities impose relationships among the different parameters of the cellular dialogues, such as the maximal secretion rates and sensing thresholds (Figure 1D; Sections S5.3 and 5.4). By solving these inequalities, we found that only five cellular dialogues – the exact same ones that we computationally identified – can satisfy all six inequalities and thus generate non-oscillatory dynamic spatial patterns (i.e., the ones that do not involve concurrent dynamic temporal patterns) (Figure 3D - red boxes). In accordance with the computational screening, the analytical approach revealed that only two types of rectilinear waves are possible, each differing by which gene-expression state is assigned to each layer: all cellular dialogues with cellular dialogue 15 as the common motif (i.e., molecule-1 promotes its own secretion) generate one type of rectilinear wave (Figure 4D - top row) while the others, having cellular dialogue 19 as the common motif (i.e., molecule-2 promotes its own secretion), generate the other type of rectilinear wave (Figure 4D - bottom row). As an exception, cellular dialogue 33 can generate both types of travelling waves because nested inside it as subgraphs are both cellular dialogues 15 and 19.

**Figure 4.**
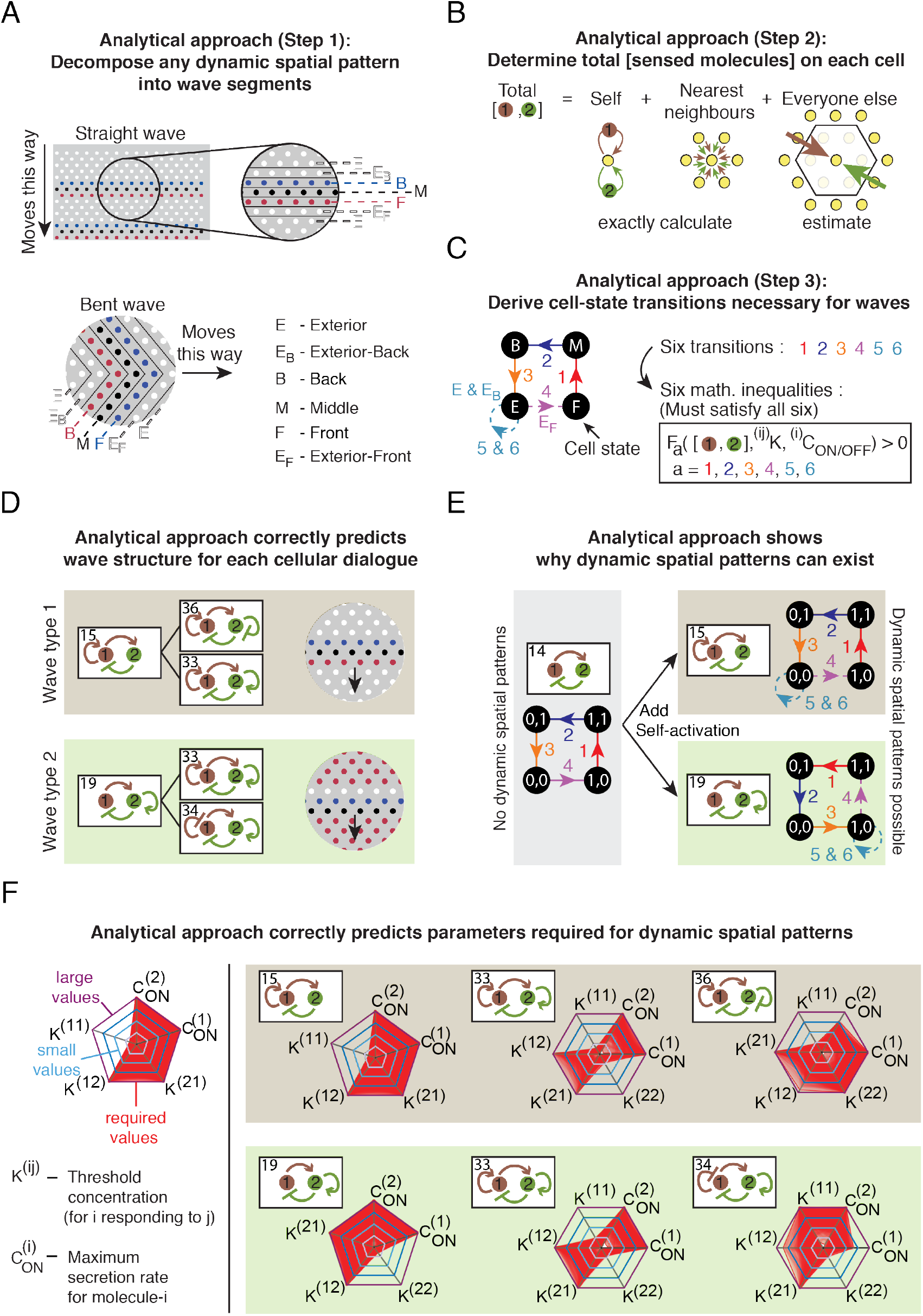
Analytical framework predicts and explains how cells can sustain dynamic spatial patterns. **(A-C)** Three-step overview of an analytical (pen-and-paper) approach to understanding the simulations (see Section S5). **(A)** Step 1: Decompose straight (top) and bent (bottom) waves into distinct layers of cells - cells of the same layer have the same gene-expression state. **(B)** Step 2: Estimate the total concentrations of molecules that a cell senses by exactly calculating the portions of those concentrations that are due to the cell itself and its nearest neighbors, and by approximating the portions of the total concentrations that are due to further-away cells (approximation scheme in Section S5). (C) Step 3: (right) Directed graph-representation showing how a cell must transition to distinct layers shown in (A) at each timestep, which is explained by six mathematical inequalities that are derived through step 2 (see Section S5). **(D)** Numerically solving the six inequalities in shows that only two types of waves, shown here, are possible and which cellular dialogues can produce them (cellular dialogues 15, 36, and 33 for wave type 1; cellular dialogues 19, 33, and 34 for wave type 2). **(E)** Adding self-activation to cellular dialogue 14 yields, in the left column, cellular dialogues 15 and 19. Directed graph-representation showing the gene-expression transition of a cell for each cellular dialogue is shown (details in Section S4). **(F)** Parameter values that allow for sustaining of rectilinear waves, when represented as red points, form a dense region (red region) as shown in these spider charts. These parameter values satisfy the six inequalities derived by the analytical theory (C) (see Fig. S5 for a direct comparison with parameter values found purely through computational search). The spider charts show the following parameters: threshold concentrations *K*^*ij*^ for each molecular interaction and the maximum secretion rate 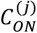 for each of the two molecules.

To understand why only these five cellular dialogues (Figure 4D) generate dynamic spatial patterns, we considered the directed-graph representation of the cellular dynamics that we introduced earlier (Section S4). For a wave, the directed graph must contain a cyclic path that goes through all four nodes – one node for each gene-expression state – since an exterior cell must eventually become a front-layer cell, then a middle-layer cell, then a back-layer cell, and then finally an exterior cell again (Figure 4C). We can intuitively see that cellular dialogue 14, which is the backbone of all five cellular dialogues that generate dynamic spatial patterns (Figure 3D - red boxes), can potentially produce a cyclic graph with these four nodes (Figure 4E - left panel) as long as they permit parameter values that allow for the cyclic traversal by each cell. This is because starting with a gene-expression state of (1, 0) – where the 1 means ON-state for molecule-1 and the 0 means OFF-state for molecule-2 – may lead to (1, 1) due to molecule-1 promoting molecule-2 secretion, which then may lead to (0, 1) due to molecule-2 repressing molecule-1 secretion, which then may lead to (0, 0) due to there being not enough molecule-1 for promoting molecule-2 secretion, and finally, this may lead back to the starting state, (1, 0), due to there being not enough molecule-2 for inhibiting molecule-1 secretion. However, such a cycle through the four nodes alone is insufficient for sustaining a wave because the exterior cells must remain as exterior cells unless they are adjacent to the front or back layer (Figure 4C). But if the exterior cells have state (0, 0) and the front-layer cells have state (1, 0), then the exterior cells near the front layer (i.e., the exterior-front cells) would sense more molecule-1 than the exterior cells that are further away from the wave. Modifying cellular dialogue 14 by having molecule-1 promoting its own secretion, as in cellular dialogue 15, would create the possibility of the exterior-front cells activating molecule-1 secretion and thus transition to (1,0) at the next timestep, thereby becoming front-layer cells whereas the exterior-layer cells remain in the (0, 0) state (Figure 4E - top right). A similar qualitative reasoning also deduces an analogous result for cellular dialogue 19 (Figure 4E - bottom right).

To realize the qualitative scenario described above, a cellular dialogue must contain parameter values that satisfy all six inequalities that we derived (Figure 4C). We found that the five cellular dialogues indeed admit such parameter values and that these values - obtained through the analytical approach - nearly perfectly match those found in the computational screen (Figure S3 and Figure S7C). We can represent these parameter values as spider charts (Figure 4F and Figure S7C), which show that each of the five cellular dialogues can still realize dynamic spatial patterns even after we vary the parameter values over many orders of magnitude. The spider charts also geometrically reveal a common feature among the five cellular dialogues: the threshold concentration must be low for a molecule that promotes its own secretion (Figure 4F - note the inward indentations in the red spider webs along the axis that represents the threshold concentration). This makes sense because, for all types of rectilinear waves (Figure 4D), the exterior-front cells need to turn on the secretion of a molecule that promotes its own secretion by sensing it from the other layers and having a low activation threshold for that molecule would facilitate this. Taken together, our analytical approach unveiled how cells can sustain dynamic spatial patterns.

### Self-organization occurs through a three-stage, “order-fluctuate-settle” mechanism

We now turn to the self-organization process itself. Given that many of the dynamic spatial patterns are travelling waves and that more complex dynamic spatial patterns can be built from gluing together multiple rectilinear waves, we focused on travelling waves and the core features of their self-organization processes. Our simulations revealed that travelling waves form in three stages (Figure 5A and Video S2). First, a field of cells whose gene-expression levels form a completely disorganized spatial configuration rapidly becomes more spatially ordered, meaning that the gene-expression levels of neighboring cells tend to become more correlated over time. To quantify the degree of spatial organization, we used a “spatial index” - a metric that our previous work established whose value is zero for a completely disorganized spatial configuration and increases towards one as the spatial configuration becomes more organized (Figure 5B - left panel’s inset) [Maire & Youk, 2015; Olimpio et al, 2018]. In the following, we equate one timestep of a simulation with one minute and express time in minute or hours. Then we find that this rapid spatial ordering typically takes less than an hour (Figure 5A - green arrow and Figure 5B - left panel). At the end of this process, the cells have formed multiple whirlpools of wavelets (Figure 5A - frame at 0.33 hours). Thus begins the second stage of self-organization: long-lived complex dynamics - lasting for days or even weeks - in which multiple wavelets travel through the field of cells, meeting and annihilating each other, all the while as the cells form new wavelets to replace the destroyed ones (Figure 5A - filmstrip from 0.33 hours to 55 hours). During this days-long dynamics, the spatial organization neither stably increases nor decreases - the spatial index wildly and unpredictably fluctuates over time (Figure 5B - left panel and Figure S4). This wild fluctuation in the spatial index (Figure 5C - left panel and Figure S4) represents multiple wavelets forming and annihilating at various, seemingly random locations and wavelets unpredictably morphing over time, all despite the fact that the simulations are completely deterministic.

**Figure 5.**
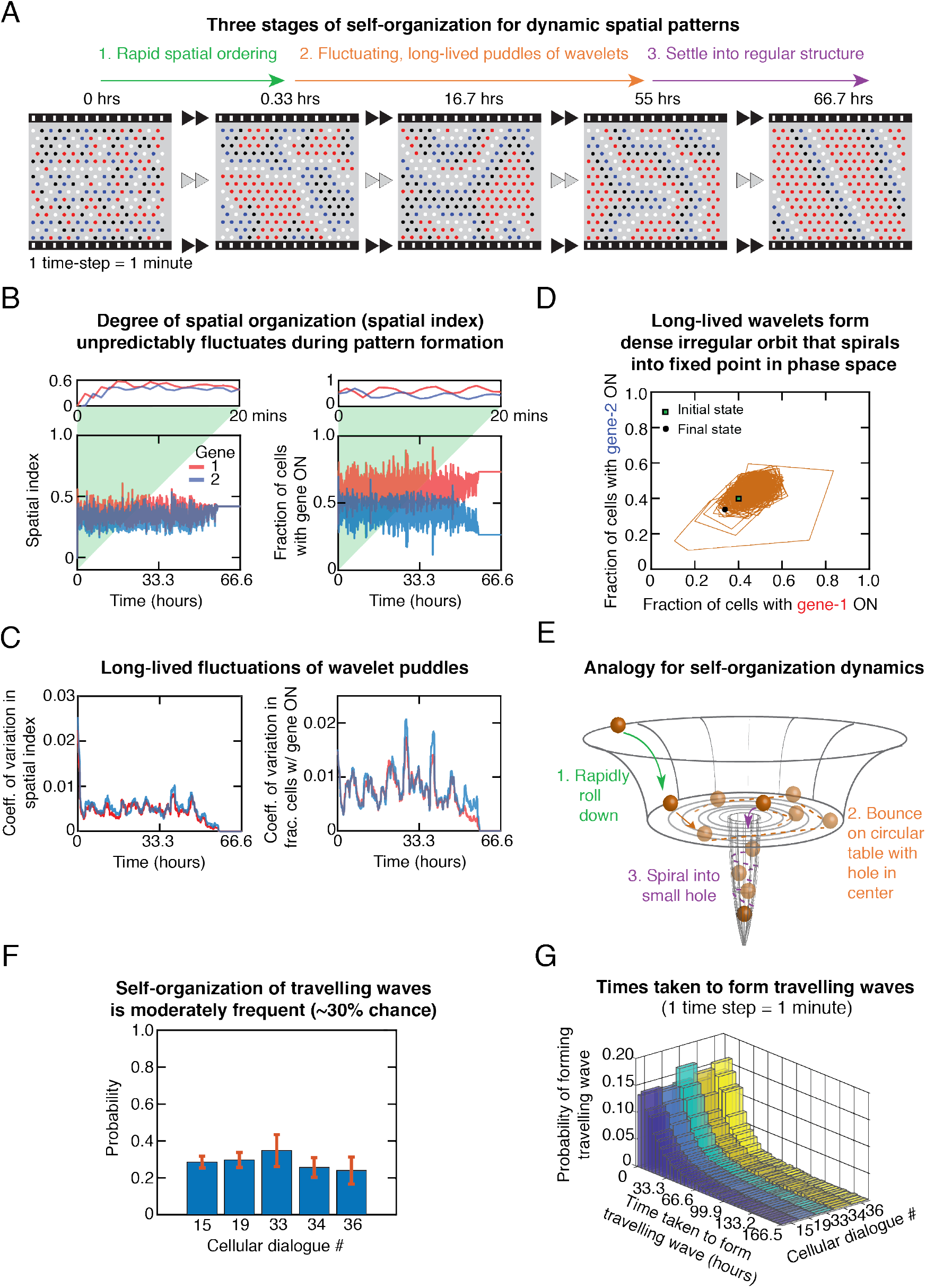
Three-step, “order-fluctuate-settle” process leads to formation of dynamic spatial patterns. Snapshots of a simulation showing the three stages of a traveling-wave formation - the three stages are described above the filmstrip. Assuming that one time-step of a simulation represents one minute, indicated above each snapshot is the elapsed time in hours. Color scheme for cells is the same as in Fig. 2. **(B)** Two macroscopic parameters - the spatial index and the fractions of cells with a particular gene ON - plotted as a function of time for the wave-forming simulation shown in (A). One minute represents one timestep. (Left panel) The spatial index - with magnitude between zero and one - measures the degree of spatial organization (zero means complete disorder, i.e. no spatial correlation in gene-expression among cells, and increasing values correspond to more spatial organization). Inset shows the spatial index rapidly increasing for the first twenty timesteps. Spatial index for gene 1 (red) and gene 2 (blue). (Right panel) Fractions of cells with gene 1 ON (red) and of gene 2 ON (blue) for a typical wave-formation process. Inset shows the first twenty timesteps. **(C)** For data in (B) and genes 1 (red) and 2 (blue), we used a moving window to compute the moving coefficient of variations in the spatial index (left panel) and in the fractions of cells with the specified gene ON (Section S1.3). **(D)** For a typical simulation that self-organizes into a travelling wave, we plot the trajectory in phase space formed by the fractions of cells with gene 1 ON and gene 2 ON. The trajectory begins at the square (first timestep of the simulation) and terminates at the circle (last timestep of the simulation). **(E)** Analogy for the three-stage self-organization process - a billiard ball rolls down a bowl, bounces around on the flat circular bottom, and then fall through a tunnel after finding a small hole drilled into the circular bottom. **(F)** Probability of forming a traveling wave for each of the five cellular dialogues (detailed results in Fig. S8). Averages are over all parameter sets for which at least one travelling wave formed in the computational screening (Section S2). Error bars are standard errors of the mean. **(G)** Distributions of the time taken to form a traveling waves for each of the five cellular dialogues that enable cells to form dynamic spatial patterns (detailed results in Figure S5).

Crucially, we verified that the same spatial configuration is never repeated throughout the days-long dynamics which could, in fact, last for weeks or longer if we do not terminate the simulations (i.e., some fields of cells never reach a steady-state and never attain a dynamic pattern within our observation times). Such unpredictable, chaos-like dynamics is followed by the third and final stage of the self-organization process: the wavelets die down and as this occurs, a more rigid, spatially ordered structure that travels as a wave emerges (Figure 5A - last frame). During this final process, the spatial index’s fluctuation rapidly decays, typically over a few hours. The spatial index then eventually settles at a value that is higher than the value that it had, on average, during the wild fluctuations. This settling process takes a few minutes to several hours (Figure 5A - purple arrow and Figure 5B - left panel). Importantly, leading up to this last stage, there are no clear indications that a well-organized regular shape will emerge. This highlights the unpredictable, seemingly chaotic nature of the self-organization dynamics.

The spatial index, one for each gene, represents a macrostate variable - it is a single number that measures how much of a spatial correlation there is for a single gene among cells. Another macrostate variable is the fraction of cells that have the same gene-expression level (i.e., fractions of cells that have gene-i in the ON-state). There are two such fractions, one for each gene. During the self-organization process, the values of these two fractions wildly fluctuate over time - just like the spatial indices - as the constantly modulating wavelets keep changing their shapes and meeting and annihilating each other for days. Afterwards, the two fractions’ fluctuations quickly decay over time - the decay takes a few hours whereas the whole self-organization process takes days - and eventually settle at steady-state values (Figure 5B-C: right panel and Figure S4). When we view the temporal change of these two fractions as a trajectory in a plane defined by the two fractions (i.e., phase space), we see an irregular orbit that eventually stops at a single point (Figure 5D - black circle). Specifically, a point in the two-dimensional phase space - representing the values of the two fractions at a given time - moves erratically within a restricted region of the plane. If we follow the trajectory with a pencil, we would obtain a jagged curve that densely and nearly entirely fills the whole space within the restricted region - reminiscent of trajectories of a chaotic system - that encloses the single point at which the trajectory eventually terminates (Video S4). This description remains unchanged if we restart the simulation with different starting values of the two fractions.

The phase-space trajectory described above suggests the following analogy for the self-organization dynamics (Figure 5E): a ball quickly rolls down a steep side of a large bowl, speeding up as it does so, until it reaches the bowl’s flat bottom. This is the first stage of self-organization in which the decreasing height represents more spatial ordering (Figure 5E - green arrow). After reaching the frictionless, flat circular bottom, the ball rapidly bounces off the side walls, like a billiard ball, without ever losing its speed (Figure 5E - brown dashed lines). This bouncing ball, which would produce seemingly chaotic yet deterministic motion - as Newton’s laws of motion are deterministic - represents the second stage of self-organization in which multiple whirlpools of wavelets are created and destroyed. Eventually, the ball would find the small hole, fall into it, and then spiral its way downwards along the side walls of the trench through the hole, until it settles at the bottom of the trench (Figure 5E - purple arrow). This would represent the third and the final stage of the self-organization. The shape of the bowl and the location of the trench would be determined by the parameters of the cellular dialogue.

For each of the five cellular dialogues that can yield dynamic spatial patterns, we found that for parameter values that enable dynamic pattern formations, approximately 30% of the initially disorganized configurations successfully self-organized travelling waves (Figure 5F). Furthermore, for all five cellular dialogues, we discovered that the probability of forming a travelling wave at a given time is well described by an exponential distribution (Figure 5G and Figure S5A), with a characteristic decay time of thousands of timesteps (i.e., tens of hours if one timestep is one minute). This strongly suggests that travelling wave formation is a memoryless process whereby at each timestep, the probability that the next timestep yields a travelling wave remains the same regardless of at which timestep the simulation is at. This reflects our observation that the self-organization dynamics seems unpredictable and chaotic, meaning that watching simulations that yield a traveling wave or another dynamic spatial pattern does not give the observer a sense that the cells are getting anywhere closer to forming a travelling wave as time passes by (Figure S5B-D).

### Dynamic patterns with more complex elements

We next extended our investigation by relaxing the two main constraints in the simulations - the infinite Hill coefficient and having cells on a regular lattice. We modified the simulations by separately adding four, more complex elements (Figure 6A and Section S6): (1) stochastic response to the signaling molecules (Figure 6A - top left), (2) a finite Hill coefficient that could vary over a wide range (i.e., cells no longer digitally respond to the signaling molecules) (Figure 6A - top right), (3) randomized locations of cells instead of each cell residing on a regular lattice (Figure 6A - bottom left), and (4) random (diffusive) motion of each cell (Figure 6A - bottom right). We quantitatively tuned each element (i.e., we varied and quantified by how much each cell moved). For each element, we asked two questions: Can the cells still form travelling waves if they start with a completely disordered spatial configuration? (Figure 6B - top) - this probes the self-organization capability - and (2) can cells still sustain travelling waves after forming them? (Figure 6B - bottom) - this probes whether dynamic spatial patterns can be sustained after forming. In general, we found that cells could still form a wide range of dynamic spatial patterns with the four additional elements (Figure 6C). For example, we discovered that cells under the influence of a moderate noise could form a band that travels as a wave despite a number of cells stochastically obtaining the “wrong” (incoherent) gene-expression state. In this case, the wave thus propagates while stochastically evolving (Figure 6C - top left and Video S6). As another example, we discovered that even when we randomly arrange cells in space, instead of on a regular lattice, the cells could still form never-ending, complex wavelets (Figure 6C - bottom left and Video S7). More generally, by running many simulations for each of the four complex elements, we discovered that the dynamic spatial patterns that we previously observed, on a regular lattice with an infinite Hill coefficient (Figure 2), still formed as long as the amount of the deviation introduced by the four elements, relative to the regularity of the lattice and the infinite Hill coefficient, was not too large but still not negligible (Figure 6D). For instance, we found that with a moderate noise, dynamic spatial patterns continued to form and persist (Figure 6D-E - top left and Figure S6A). By varying the Hill coefficient over a wide range, we discovered that dynamic patterns could still form for sufficiently low Hill coefficients, up to a value of ~3, but these did not typically include pure (single) travelling waves (Figure 6D - top right and Figure S6B). However, already formed traveling waves were able to persist up to values of Hill coefficient up to ~4 (Figure 6E - top right). This indicates that the finiteness of the Hill coefficient is mainly detrimental to the self-organization of travelling waves whereas it is less detrimental to formation of more complex dynamic spatial patterns (e.g. composed of multiple rectilinear waves) and the cells’ ability to sustain a travelling wave once it is formed. With a Monte Carlo algorithm that randomly displaces cells and quantifies the amount of the resulting “lattice disorder” (Section S6.3), we found that dynamic spatial patterns still formed and persisted even with a high degree of spatial disorder (Figure 6D-E - bottom left; Figure S7C). Notably, even with saturating amounts of spatial disorder, we still observed self-organized wavelets that propagated, albeit with a lesser degree of regularity than in a regular lattice. When we caused the cells to diffusively move about throughout the field - we tuned the cells’ motility by adjusting the diffusivity of their Brownian motion (Section S6.4) - we found that large-scale, uncoordinated motion of cells prevented any kind of dynamic spatial patterns from stably propagating, as large variations between the local environments of individual cells tended to diminish the cells’ ability to reliably propagate information across space (Figure 6D - bottom right). However, we found that motile cells could still propagate waves, once formed, for an extended amount of time before the wave disintegrated even when the cells had a high degree of diffusive motion (Figure 6E - bottom right). Together, these results strongly suggest that diffusively moving cells can sustain travelling waves as long as the waves travel sufficiently rapidly (i.e., compared to the average speed of the cells’ motion). Finally, we considered the influence of external gradients, modelled as spatially varying parameters (Section S6.5), on travelling wave formation. Researchers have suggested that parameter gradients can influence the orientation of Turing patterns such as stripes [Hiscock & Megason, 2015]. Despite our system not being a Turing-patterning system, we observed that a simple step function profile in one of the parameters can have a considerable influence on the direction in which the travelling waves moved. Namely, the waves tended to align perpendicularly to the gradient (Figure S6C).

**Figure 6.**
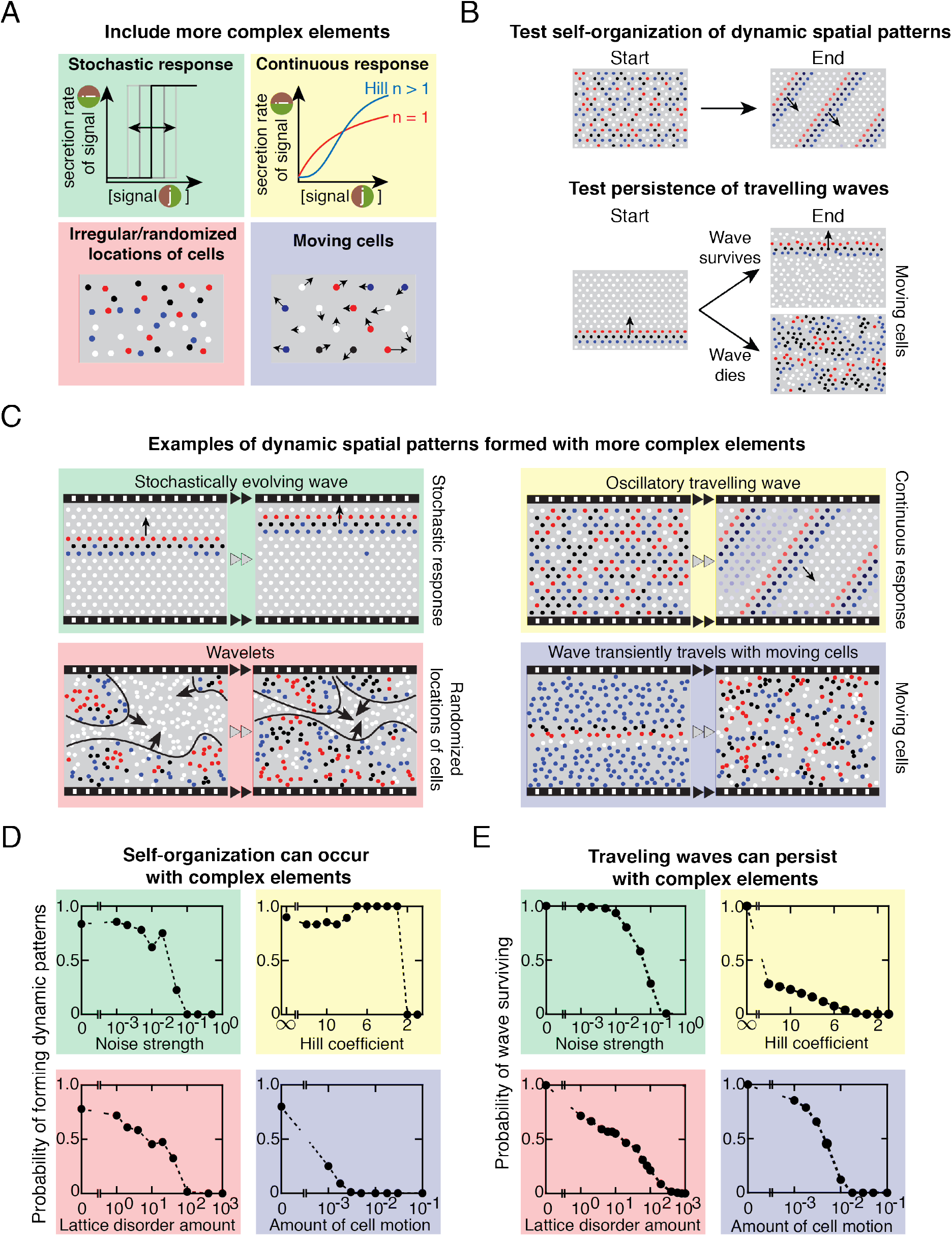
Dynamic spatial patterns still form even with more complex elements. **(A)** Schematic of four additional, more complex elements that we added to our computational screen. **(B)** We examined two features with the elements in (A): (Top) Can a disorganized field of cells still self-organize dynamic spatial patterns? (Bottom) Starting with a travelling wave - since it is the most ubiquitous form of dynamic spatial patterns - can the cells sustain it? **(C)** Examples of dynamic spatial patterns formed for each of the elements shown in (A). Colored boxes that enclose the filmstrips correspond to the colors used for each element shown in (A). **(D)** Fraction of simulations that form a dynamic pattern as a function of the deviation from the more idealized setting - cells placed on a regular lattice and responding digitally with an infinite Hill coefficient - in which the results for Figures 1–5 were reported. Four colored boxes, with each color corresponding to colored box in (A) that shows the modified element in the simulations. For each data point, we ran a large set of simulations with random initial conditions and classified their final states (see Figure S6 for a detailed classification). For finite Hill coefficient, we included all dynamic patterns including never-ending wavelets. For all other cases, we report the fraction of dynamic spatial patterns. All results here are for cellular dialogue 15. **(E)** Fraction of simulations with cellular dialogue 15 that can sustain a travelling wave for at least one full period after starting with a travelling wave. We took parameter values for which the simulations with simpler elements (i.e., infinite Hill coefficient and cells on a regular lattice) can propagate travelling waves.

## DISCUSSION

### Dynamic-pattern producing cellular dialogues that we identified are found in experimental systems

Some of the dynamic-pattern forming cellular dialogues that we identified have already been observed in diverse natural and synthetic biological networks. These cellular dialogues, as a common feature, all have interlocked positive and negative feedbacks (Figure 3D). Experimentalists have observed that, in autonomous cells (i.e. without cell-cell communication), such interlocked feedbacks can cause gene-expression levels to robustly oscillate over time with a tunable frequency [Stricker et al., 2008, Tsai et al., 2008, Li et al., 2017]. Experimentalists have also observed that when a synthetic circuit enables *E. coli* cells to communicate through a cellular dialogue that resembles our cellular dialogue 20 (Figure 3D), the cells’ gene-expression levels (GFP-level) collectively and synchronously oscillate over time and, under certain conditions, spontaneously form travelling waves [Danino et al., 2010]. More generally, the activator-inhibitor structure that is at the core of cellular dialogue 15 is qualitatively similar to the structure of the FitzHugh-Nagumo (FHN) model, which is a prototype model for describing excitable systems such as cells whose gene-expression, metabolite, or ion levels oscillate over time and/or form traveling waves [Gelens et al., 2014; Sgro et al., 2015; Hubaud et al., 2017]. Cellular dialogue 15 has an activating molecule that promotes its own production and an indirect negative feedback through the second molecule. This indirect negative feedback is analogous to the slow repression that exists in the FHN model. As a related matter, the interlocked positive-negative feedback loops of the dynamic-pattern forming cellular dialogues resemble the activator-inhibitor systems that generate Turing patterns [Kondo & Miura, 2010]. However, the mesoscopic numbers of cells in our simulations, with these cellular dialogues, do not generate Turing patterns such as stripes or spots of a fixed size. Conversely, while our simulated cells generate travelling waves under a wide variety of conditions, Turing systems with just two diffusing and reacting chemicals are thought to only generate static patterns, with dynamic patterns only arising when three or more molecules are considered [Turing, 1952]. These are in contrast with our findings based on two-molecule cellular dialogues.

### Our computational and analytical framework may guide in identifying currently unknown mechanisms in experimental systems

Although we focused on cellular dialogues with two molecules and the two genes that they control, our software can easily be modified to include multiple - more than two - extracellular molecules and genes as well as arbitrary regulations of those genes (as showcased by our inclusion of finite Hill coefficients). These extensions would allow one to explore more complex ways that cellular dialogues can mediate dynamic pattern formations. These extensions, the analytical method for analyzing the simulations that we established here, and this study’s results on two-molecule cellular dialogues may together shed light on poorly understood systems in which multiple signaling molecules interact with each other. For many experimental systems, the regulatory links among the various molecular players remain unknown (Figure 7). For example, to form somites, researchers have found that three signaling molecules - Fgf, Notch and Wnt - are regulate one another. But how Wnt and Notch regulate each other so that their levels coordinately oscillate over time remains unknown (Figure 7A) [Oates et al., 2012; Harima & Kageyama, 2013; Sonnen et al., 2017]. Modifying our software to include three-molecule cellular dialogues, and then applying our analysis method to analyze those simulations, may address this question. Doing so may also help identify, in stem cells, the as-yet unknown regulatory links among three signaling molecules - Bmp, Wnt, and Nodal - that lead to self-organized spatiotemporal waves (Figure 7B) for which the Turing mechanism is not involved [Chhabra et al., 2019]. In the *Arabidopsis Thaliana* leaves, the circadian clocks of individual cells are thought to be synchronized through self-organized travelling waves [Wenden et al., 2012; Gould et al., 2018] (Figure 7C). While these waves are known to occur through local cell-to-cell interactions, the exact interaction mechanism remains unknown [Greenwood et al., 2019]. Since the cells within these leaves are on a nearly regular lattice, our results for cells on a regular lattice may be relevant to this system. Finally, in planarian - an organism that regenerates its body after it is cut into pieces - a self-organized Wnt gradient specifies where the tail should be reformed after it is excised. Researchers believe that an as-yet unidentified signaling molecule may interact with Wnt in a mutually antagonistic way to indicate where the head should reform after it is excised (Figure 7D) [Stückemann et al., 2017]. Thus, our results on two-molecule cellular dialogues may guide in testing this hypothesis.

**Figure 7.**
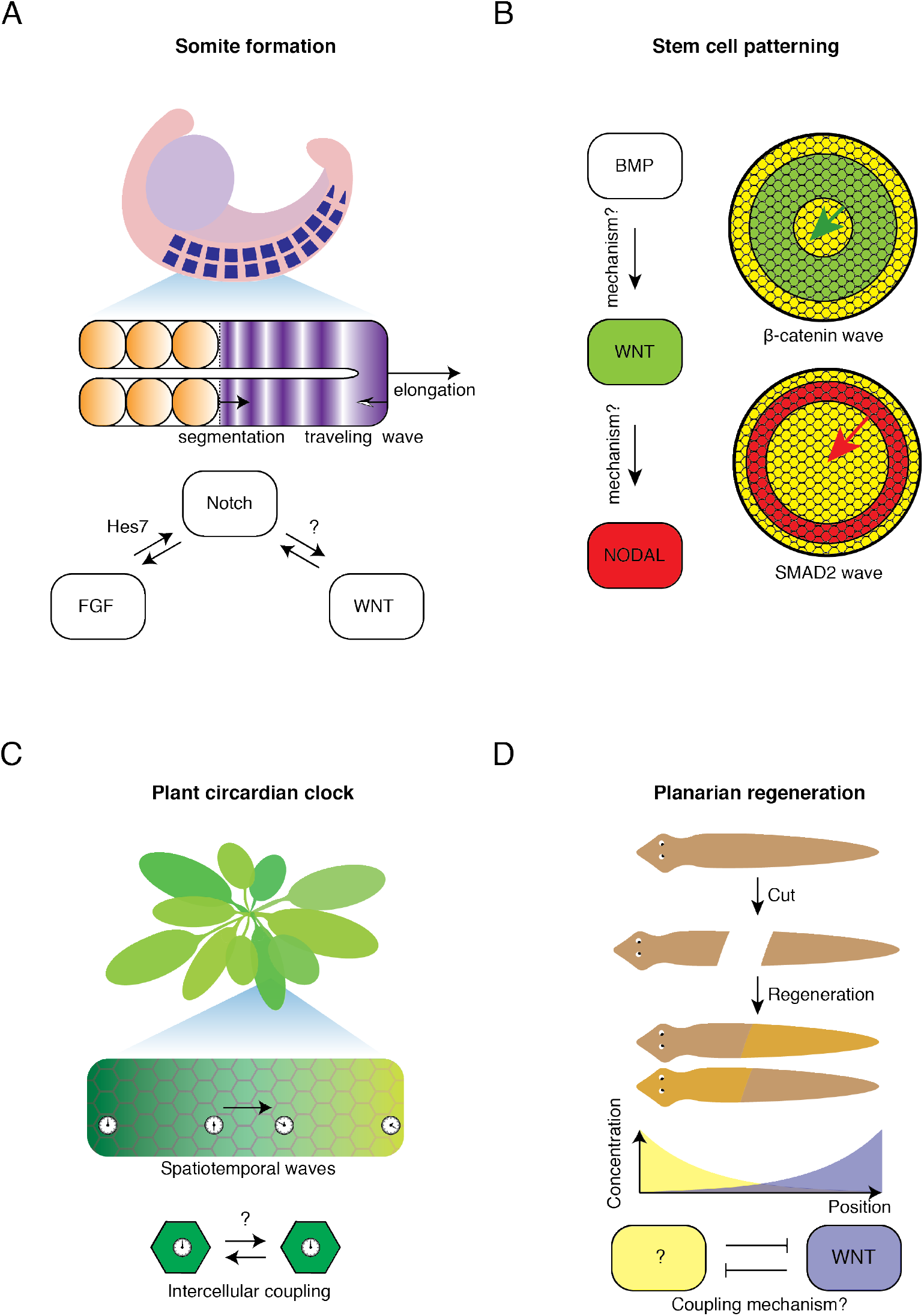
Self-organized dynamic-pattern-forming systems with poorly understood interactions that our software and analytical framework may help in elucidating. **(A-D)** Biological systems with two or more interacting pathways that generate spatiotemporal patterns but whose exact mechanisms and cellular dialogues remain poorly understood. **(A)** During somitogenesis, a wave of gene-expression states propagates along the anterior-posterior axis of an elongating, pre-somite mesoderm. The conventional view is that this wave is mediated by a coupling between individual oscillators - oscillations in expression levels of Wnt, Notch, and Fgf - and/or by large-scale gradients in the gene-expression levels for those molecules. But how Notch regulates Wnt and vice versa remain questionable while Hes7 is known to mediate the Fgf-Notch interaction (Sonnen et al., 2017). Figure partially adapted from (Oates et al., 2012). **(B)** Waves of β-catenin (green ring) and Smad2 (red ring) expression-levels propagate in a field of stem cells. Although we know that these waves form due to BMP inducing β-catenin (part of the Wnt pathway) and SMAD2 (part of the NODAL pathway), how exactly these two inductions occur remains poorly understood (Chhabra et al., 2019). **(C)** The circadian clocks of each cell within the leaf of *Arabidoposis Thaliana* are thought to be coupled to each other through an as-yet-unknown mechanism, which is suspected to involve a variety of hormones, sugars, mRNAs and other molecules (Greenwood et al., 2019). **(D)** Planarian regenerates itself after being cut into two or more pieces. This is thought to rely on mutual antagonism between gradients of Wnt expression (purple) and of an as-yet-unidentified molecule (yellow) (Stückemann et al., 2017). Figure partially adapted from (Stückemann et al., 2017).

### Summary and future outlook

Here we developed an open-source simulation software and a mathematical analysis-framework for studying the self-organization of dynamic spatial patterns in a population of cells that communicate through diffusing molecules. The software contains ingredients that anyone can alter, exclude, or replace with other elements for both research and educational purposes. With this software, we performed simulations that revealed an intricate, three-step process by which mesoscopic numbers (hundreds to thousands) of cells can form dynamic spatial patterns. Through a comprehensive computational screening, we identified all the ways in which cells can communicate with two molecules (cellular dialogues) to generate dynamic spatial patterns – patterns of gene-expression levels that continuously move without ever stopping. We then formulated an analytical framework that recapitulates the results of the simulations by deriving six mathematical equations that can be solved without simulations. By applying this analytical approach to cellular dialogues with more than two molecules, one would obtain similar set of equations (but more than six equations). We have shown that the analytical framework reproduces the correct cellular dialogues, as well as the parameter sets, for which dynamic spatial patterns can be self-organized. We discovered that self-organization of dynamic spatial patterns, by hundreds of cells, typically occurs after a long and seemingly chaotic transient phase in which the overall spatial configuration unpredictably evolves over time with no identifiable regularity, despite the underlying cell-cell interaction rules being deterministic. This highlights the high level of complexity that just a hundred or so cells can exhibit through autocrine and paracrine signaling. By including more complex elements such as stochastic sensing of the molecules and smooth responses to the sensed cytokines (i.e., finite Hill coefficient), we found that dynamic spatial patterns could still form and persist, though possibly in altered forms. By combining reaction-diffusion equations with a cellular automaton, our work takes an underexplored approach of connecting short and long timescales as well as short and long length-scales within one framework. A short time-scale characterizes the diffusing molecules while a long time-scale describes the cells’ response to the molecules. A short length-scale describes the diffusion lengths of the molecules whereas a longer length-scale characterizes the emergent patterns.

Our simulations and analytical approach revealed that the cells modelled here can have arbitrarily high parameter values and still form dynamic spatial patterns such as travelling waves, with a high success rate, as long as their various parameters maintain certain ratios (Section S5.4 and Figures S3B, S7C-D, and S8). Recent results have established that reaction-diffusion systems, without cells, that use three or more molecules can more robustly form patterns than their two-molecule counterparts [Marcon et al., 2016, Zheng et al., 2016]. Likewise, as our software and analytical framework are easily extendable to any number of molecules, it would be interesting to investigate whether increasing the number of signaling molecules would allow cells to more robustly form dynamic spatial patterns and how the “order-fluctuate-settle” mechanism may then change.

Another promising future direction is extending our software and analytical framework to include mechanical interactions such as cells interacting with the extracellular matrix or with other cells. While cellular automata that model such mechanical interactions exist [Graner & Glazier, 1992, Merks & Glazier, 2005], those that combine mechanical and chemical interactions, including intracellular signaling as in our work, remain rare yet promising [Recho et al., 2019]. Outside the context of biology, it would be interesting to interpret and analyze our work in the context of complex systems theory [Bar-Yam, 2003]. In particular, one may investigate the self-organization of dynamic spatial patterns by using information-theoretic approaches [Rosas et al., 2019; Haken, 2004] or other quantitative measures [Kaneko, 1989]. Doing so may link our findings to those of non-living chemical systems that self-organize patterns [Nicolis & Prigogine, 1977].

## SOFTWARE

GitHub: https://github.com/YitengDang/MultiCellSim

## SUPPLEMENTAL INFORMATION

- Supplementary text with mathematical details
- Figures S1-S8
- Movies S1-S7

## ACKNOWLEDGEMENTS

We thank Yaroslav Blanter and members of the Youk laboratory for fruitful discussions. H.Y. was supported by the European Research Council (ERC) Starting Grant (MultiCellSysBio, #677972), Netherlands Organisation for Scientific Research (NWO) Vidi Award (#680-47-544), CIFAR Azrieli Global Scholars Program, and EMBO Young Investigator Award.

## AUTHOR CONTRIBUTIONS

Y.D. and H.Y. designed the research. Y.D. performed the research with guidance from H.Y. D.G. helped with simulations. Y.D. and H.Y. wrote the manuscript.

## SUPPLEMENTARY FIGURES

**Figure S1.**
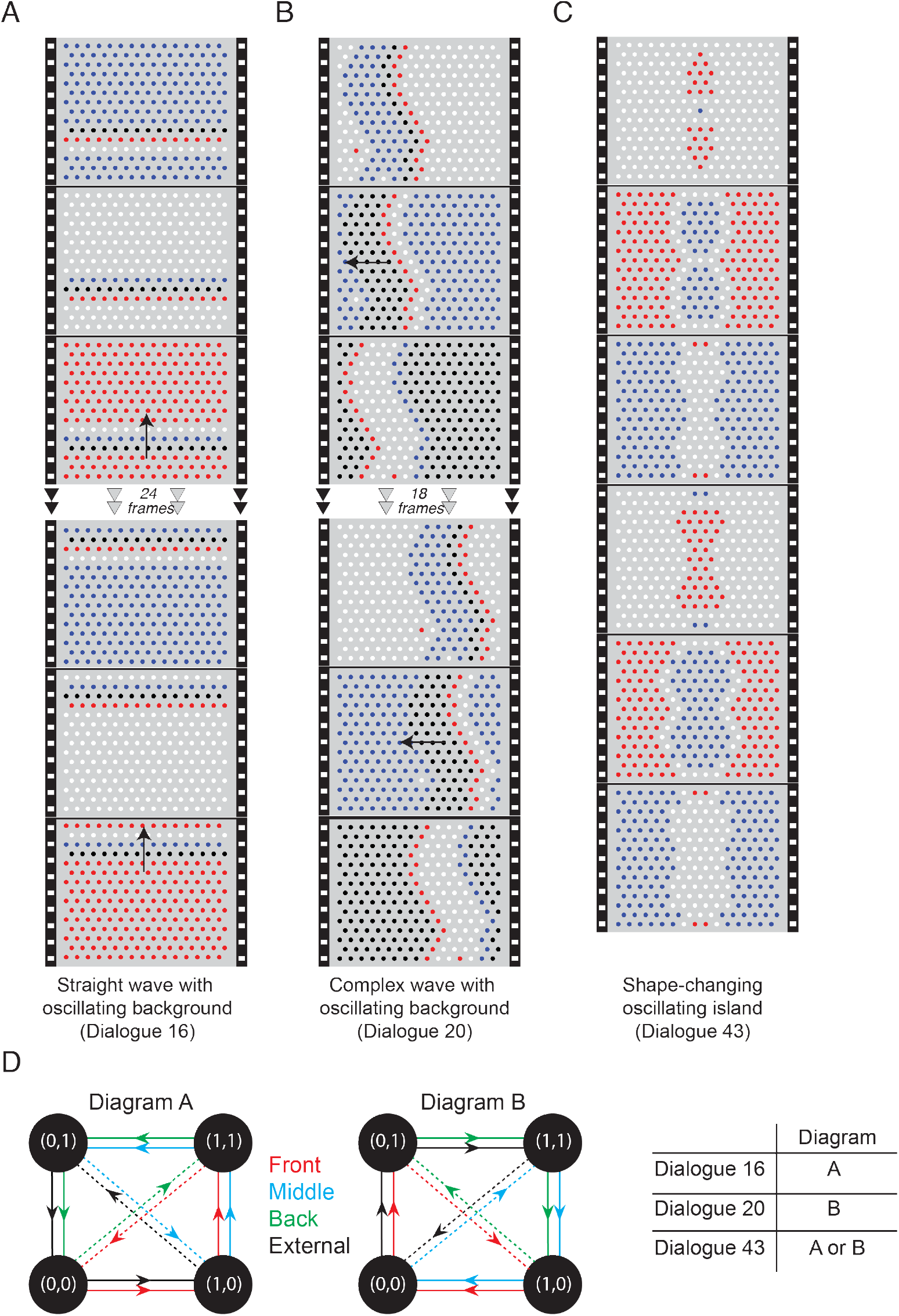
(Related to Fig. 2). Dynamic patterns with oscillatory background. **(A-C)** Examples of information-propagating with oscillatory background, generated by the corresponding networks in Fig. 3D (labelled in blue). **(D)** State diagrams showing the transitions at the single-cell level that each of these patterns undergoes (see Section S4). Each of the cells of the wave and the background cycle through three different states before the pattern moves to the next row of cells. These transitions between these states are depicted in the state diagrams as directed cycles of a graph. Different colors indicate different relative positions of the cells (see Fig. 4A). One of these transitions - indicated by the dashed lines - is concurrent with the displacement of the pattern. There are two possible state diagrams for the three cellular dialogues that generate oscillatory dynamic spatial patterns, as indicated in the table.

**Figure S2.**
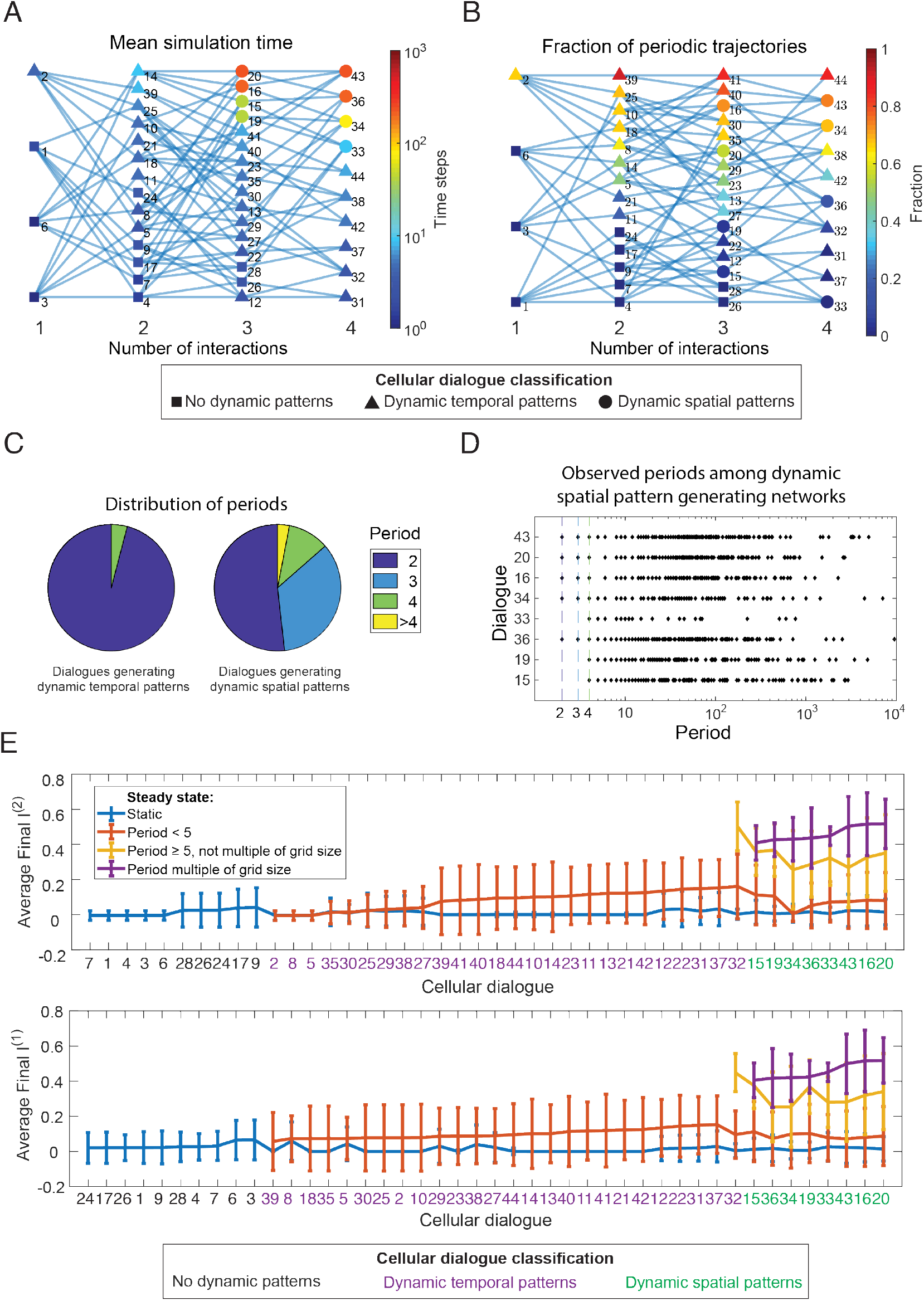
(Related to Fig. 3). Self-organization statistics for the three classes of cellular dialogues displayed in Fig. 3B-D. All results are based on the same simulation data set also used to generate Fig. 3E. **(A)** Mean simulation time across all simulations of a given network. The simulation time is equivalent to the time it takes for the system to reach equilibrium or the maximum simulation time if a trajectory never reaches equilibrium. Same graphical representation as in Fig. 3E. **(B)** Fraction of trajectories with a periodic final state, i.e. a steady state where the final pattern repeats itself after a fixed number of time steps greater than one. Same graphical representation as in Fig. 3E. **(C)** Distribution of the periods of the periodic final states among networks that generate dynamic temporal patterns (Fig. 3C) and networks that also generate dynamic spatial patterns (Fig. 3D). **(D)** Trajectory periods found in simulations of the networks generating dynamic spatial patterns (Fig. 3D). Each diamond represents a period that was observed in at least one simulation. **(E)** Average final spatial index for each of the two genes, sorted by cellular dialogue. The trajectories are divided into different classes depending on the final period of the trajectory (represented by differently colored lines). The average is taken over all trajectories within each class. For example, the purple data points show the average values for the subset of trajectories that have a period which is a multiple of the grid size. Error bars represent SEM.

**Figure S3.**
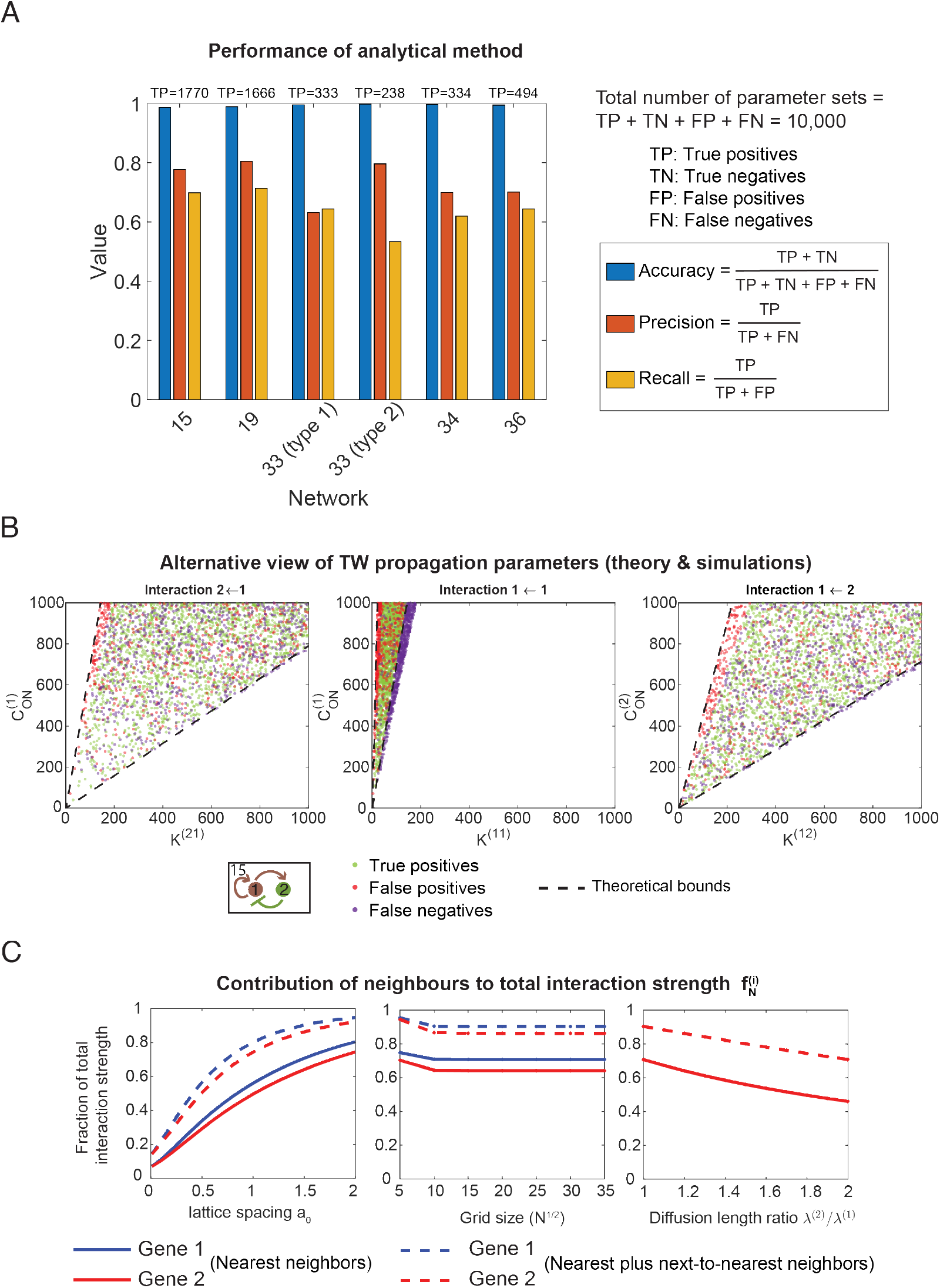
(Related to Fig. 4). Comparison between analytical theory and exact simulations. **(A)** Performance of the predictor quantified using concepts from machine learning. When viewed as a binary classifier, the analytical theory makes binary (yes or no) predictions about whether a parameter set is capable of propagating traveling waves. We compare these predictions with actual simulations to determine whether they are correct or false. The performance of the theory can then be quantified in terms of concepts such as accuracy, precision and recall (see Section S5.6). **(B)** Two-dimensional projections of the parameter sets capable of propagating traveling waves according to the analytical theory and exact simulations (see Figs. 4F and S7C for alternative representations in terms of radar charts). Since there are six varying parameters for each parameter set, we projected the parameter sets onto two-dimensional spaces spanned by the two parameters describing the strength of each interaction – the threshold *K*^(*ij*)^ and the maximum secretion rate 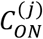. We plot the data points classified as true positives, false positives and false negatives (see Section S5.6), but leave out the true negatives, which are the parameter sets which are correctly predicted to be incapable of sustaining traveling waves. **(C)** Contribution of nearest-neighbors (*f*_*nn*_) and next-to-nearest neighbors (*f*_*nnn*_) to the total interaction strength (Equation S6 in Section S1), as a function of the lattice spacing, the grid size and the ratio between the diffusion lengths (Section S1). We plot the contribution from nearest neighbors and next-to-nearest neighbors as a fraction of the total interaction strength.

**Figure S4.**
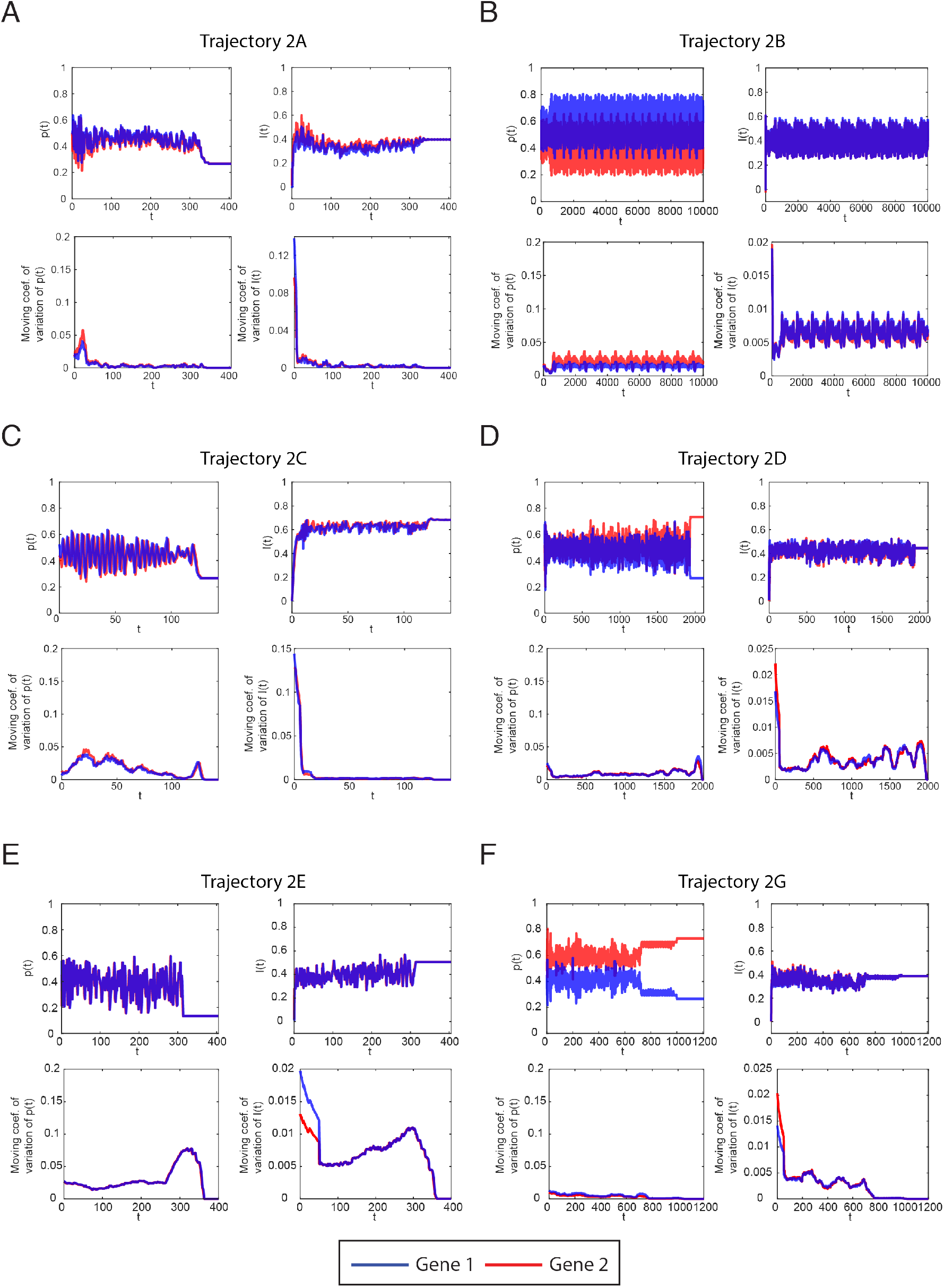
(Related to Figs. 5 and 2). Additional examples of the dynamics of the two macroscopic variables in trajectories displaying the self-organization process. We plot the fraction of cells that have a certain gene ON and the “spatial index” over time (Section S1.3). The plotted trajectories correspond to the trajectories shown in Fig. 2, as indicated by the panel titles. Time is in units of discrete time steps of the cellular automaton. Blue corresponds to gene 1, red to gene 2. For each trajectory, we show four graphs corresponding to the graphs shown in Fig. 5B and 5C. Upper left: mean fraction of cells p(t) that have the indicated gene ON. Lower left: Moving coefficient of variation for p(t) (see Section S1.3). Upper right: Spatial index I(t) for the indicated gene. Lower right: Moving coefficient of variation for I(t) (see Section S1.3).

**Figure S5.**
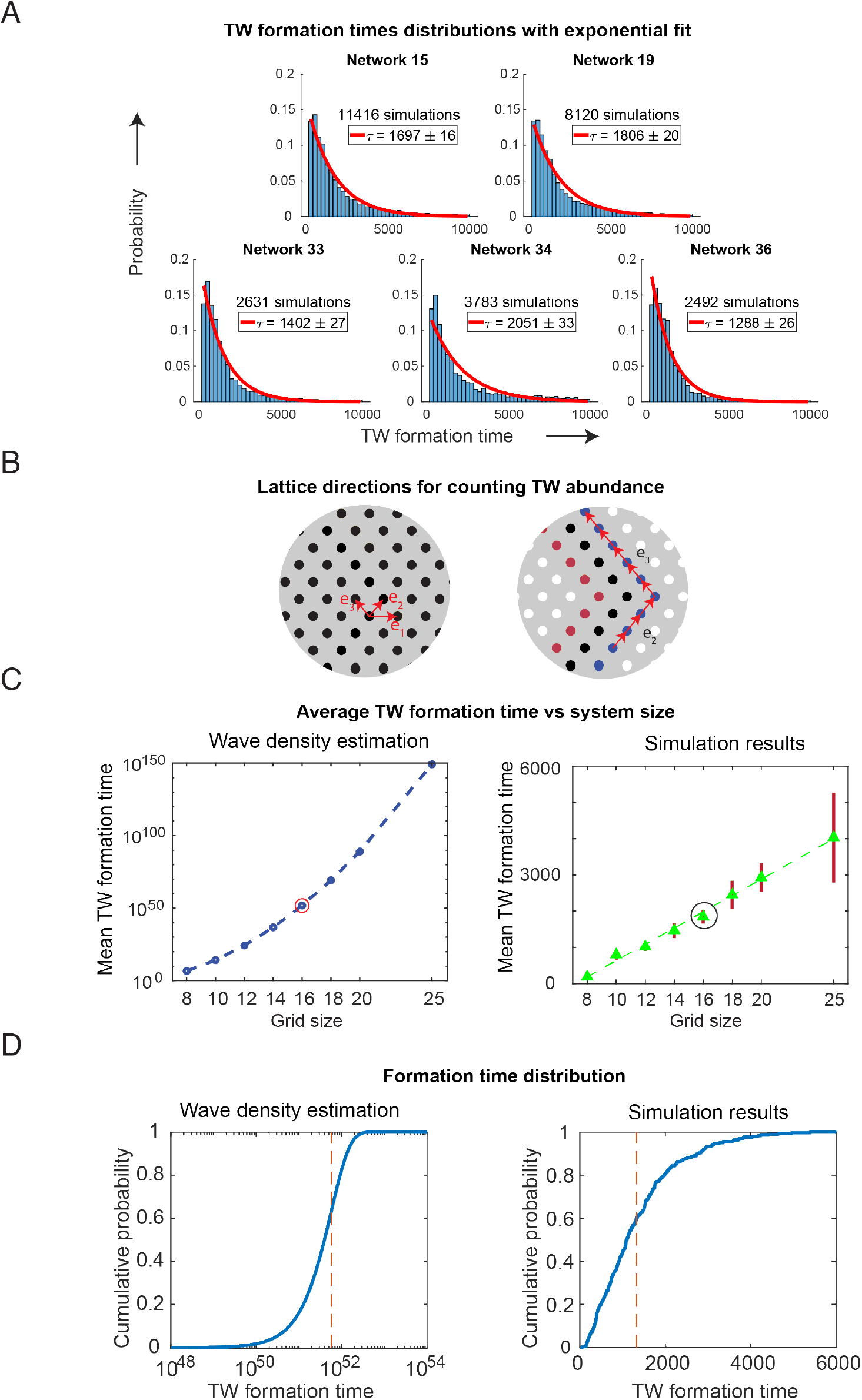
(Related to Fig. 5). Traveling wave (TW) formation time statistics. **(A)** TW formation time distributions from simulations (shown together in Fig. 5G) are fitted by exponential functions, with τ as the expectation value of the fitted exponential distribution. **(B-D)** Analytical calculation (Section S5.1-5.2) reveals that TW formation times do not follow an entirely random process, i.e. one where each next system state is randomly drawn from the set of all states, as one might suspect based on the chaotic appearance of the dynamics and the exponentially distributed formation times. **(B)** Constructions used in the calculation of the abundance of traveling waves in the system (Section S5.2). (Left) Directions on the lattice. (Right) Sketch of the construction used to characterize a single wave. By counting all ways of traversing the lattice, subject to certain constraints, we obtain an estimate of the number of forms of traveling waves of a given type. **(C)** Average TW formation time estimated from the wave density calculation (left) at different grid sizes, compared with the empirical findings from exact simulations (right). Averages are taken over all self-organized TWs among 300 simulations for each grid size. Error bars represent SEM. The highlighted data point is the grid size used in (D). **(D)** The cumulative distribution of TW formation times according to the wave density estimation (top) and from simulations (bottom; from simulation set for grid size 16 also used in C). The red dotted line represents the average formation time.

**Figure S6.**
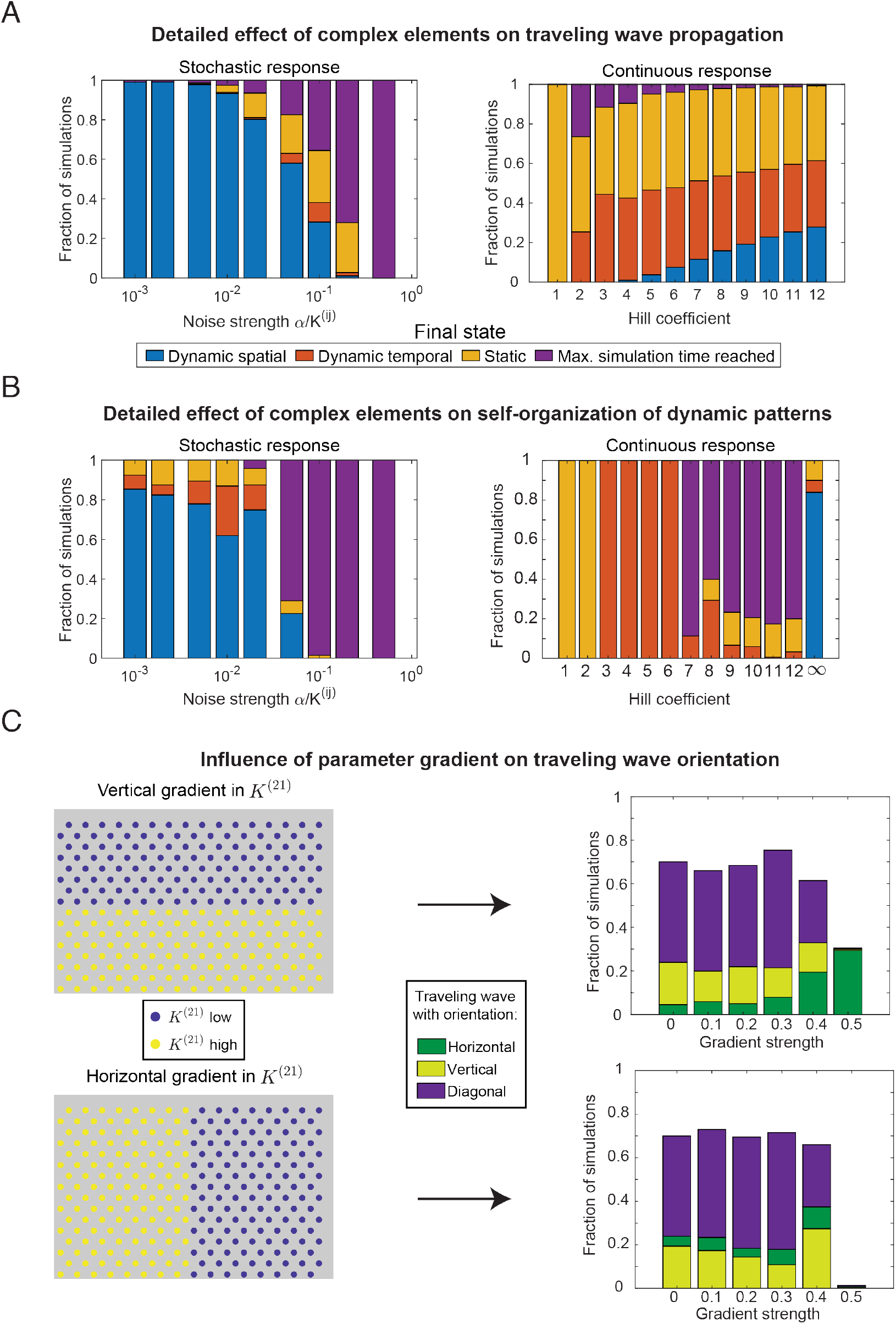
(Related to Fig. 6). Detailed effects of including complex elements into the model on the propagation and formation of dynamic patterns. **(A-B)** Detailed breakdown of simulations with two of the complex elements – stochastic response and continuous response – into four classes of patterns. Dynamic spatial pattern here refers to traveling waves specifically, dynamic temporal patterns to all other simulations that yielded periodic steady states, static patterns to simulations where the final state was non-periodic and max. simulation time reached to simulations that never settled down to a steady state within the total simulation time (10,000 timesteps). **(A)** Effect of complex elements on the formation of dynamic patterns, corresponding to the data also used in Fig. 6E (upper panels). We performed 200 simulations for each value of the noise and 150 simulations for each value of Hill coefficient. **(B)** Effect of complex elements on traveling wave propagation, corresponding to the data also used in Fig. 6D (upper panels). The parameters are chosen such that without noise and with infinite Hill coefficient a straight traveling wave, such as depicted in the lower panel of Fig. 6B, can propagate. We performed 1,000 simulations for each value of the noise and 2,534 simulations for each value of Hill coefficient. **(C)** Effect of a parameter gradient on the orientation of formed traveling waves. Specifically, we considered a step-function gradient for the threshold parameter *K*^(21)^ (Section S1) oriented along either the vertical direction (upper panels) or horizontal direction (lower panels). We classified the orientation of the formed traveling waves as the relative gradient strength (Section S6.5) is increased. The unclassified simulations in the bar graphs did not form traveling waves. We performed 200 simulations for each value of the gradient strength.

**Figure S7.**
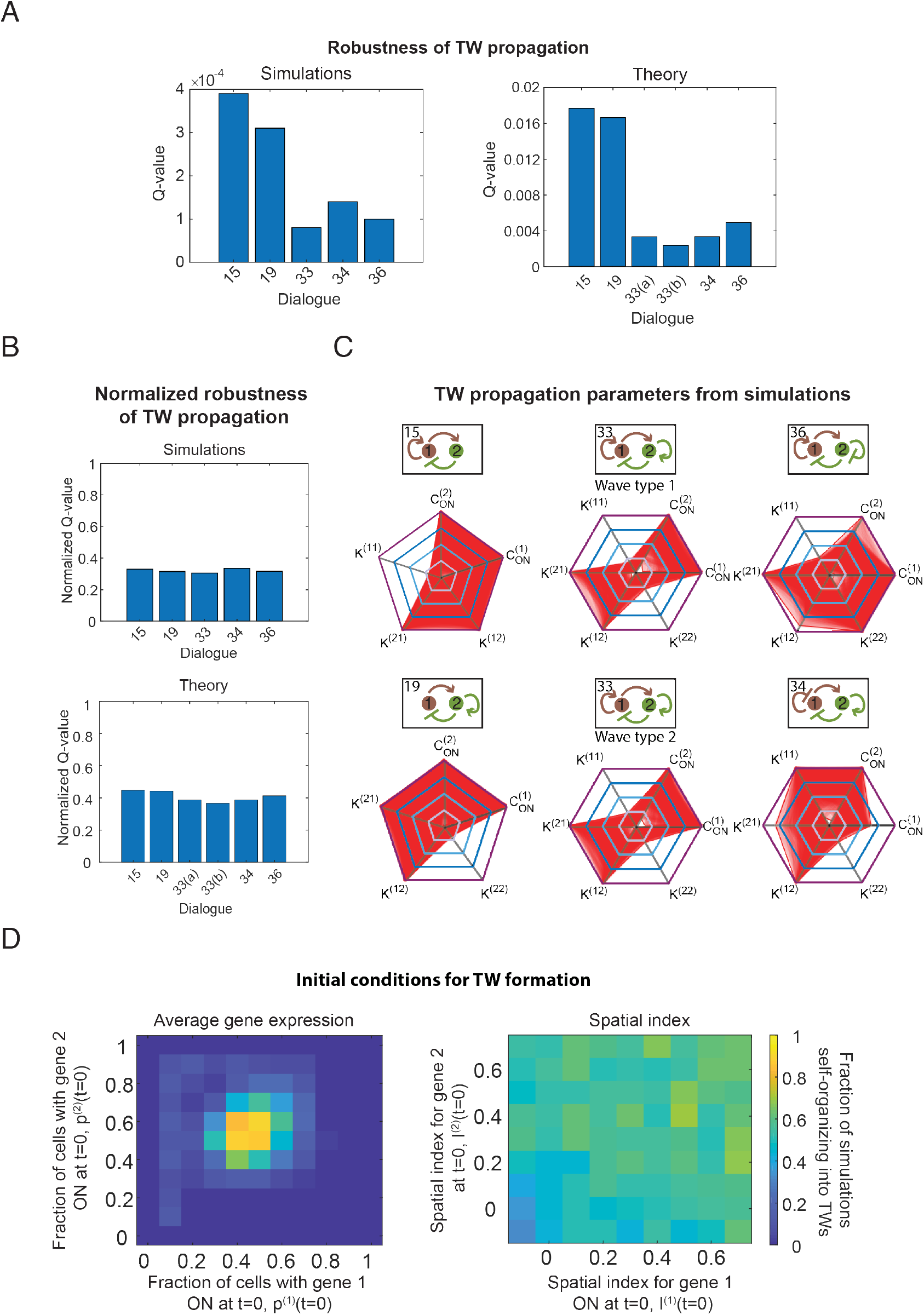
(Related to Fig. 4). Robustness of traveling waves (TWs). **(A)** Robustness of TW propagation, for each cellular dialogue capable of generating TWs in exact simulations and according to our analytical framework. We quantified robustness as the fraction of parameter sets that were capable of propagating a TW (Q-value; see Section S7). Networks 33(a) and 33(b) refer to the two types of TWs this network can generate (see Fig. 4D). Normalized robustness that considers the number of parameters for each parameter set and can be interpreted as the probability that a single random draw of each of the parameter gives a value compatible with TW formation (Section S7). Results are based on testing 10^6^ parameter sets for both theory and simulations. **(B)** Robustness of TW self-organization from random initial states. Results are based on testing 10^6^ parameter sets, with 10 simulations for each parameter set. **(C)** Radar charts or spider charts for the parameter sets for which TWs propagate as found in simulations (compare with theoretical results in Fig. 4F). **(D)** Case studies of the influence of initial state on TW formation. We varied the initial fraction of cells with each of the two genes on (for dialogue 15 at fixed parameter values; left plot) and the initial spatial index for each of the two genes (for dialogue 19 at fixed parameter values; right plot). For each combination of initial values, we performed 100 simulations.

**Figure S8.**
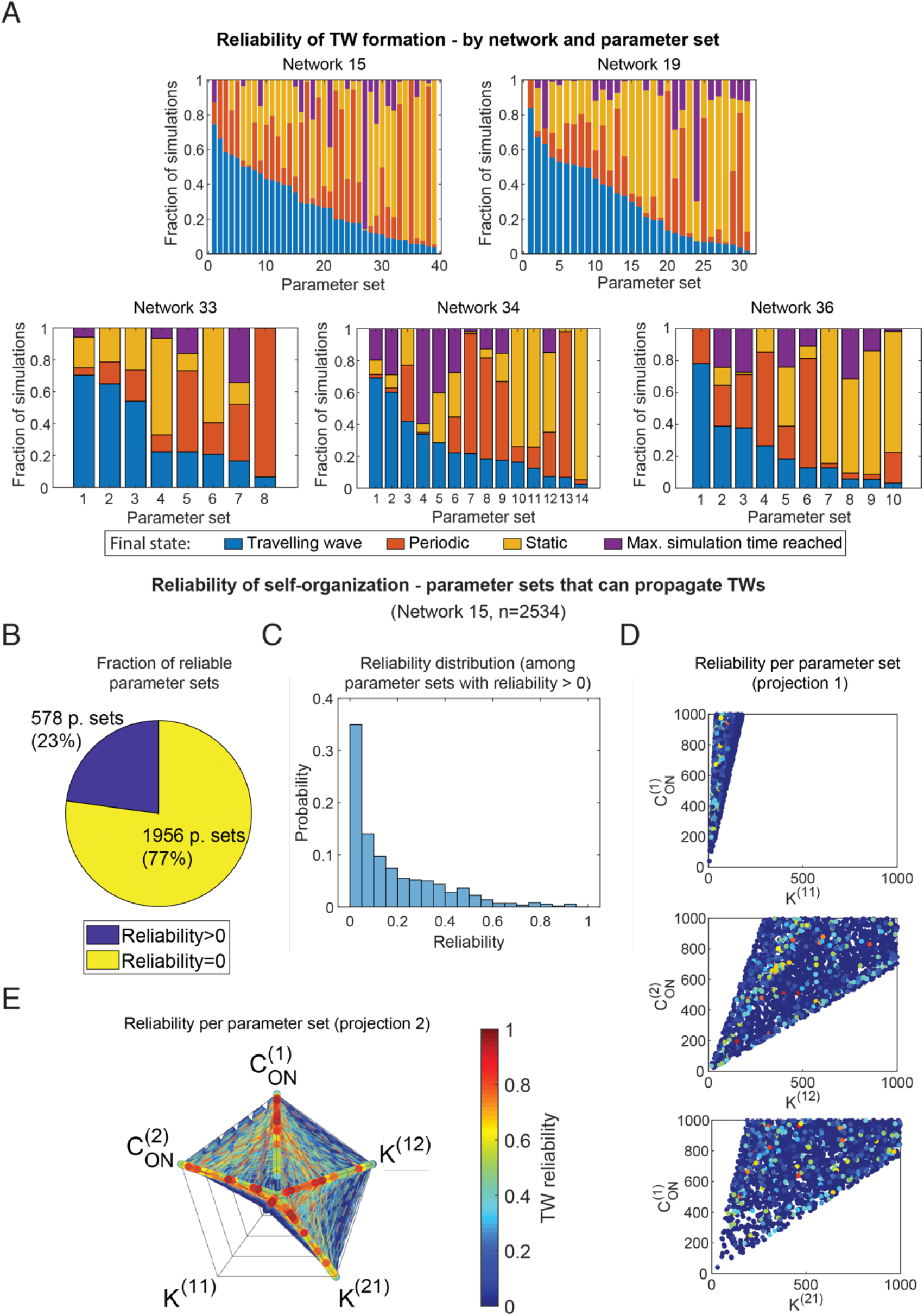
(Related to Fig. 4). Reliability of traveling wave (TW) formation, defined as the fraction of simulations with arbitrary initial conditions that self-organize into TWs at fixed parameters. **(A)** Reliability values for all parameter sets found to be capable of self-organizing TWs, for each of the five TW generating networks. The parameter sets are ordered from high to low with regard to the fraction of simulations that yielded a traveling wave (blue bars). The other simulations are classified into other periodic patterns (red), static patterns (yellow) and simulations that did not reach steady state at the maximum simulation time (purple). We performed 500 simulations with random initial conditions for each parameter set. **(B-E)** Reliability of TW *formation* for a large set of parameters (n=2534) capable of *propagating* TWs (i.e., for which a TW initial state continued to propagate indefinitely). For each parameter set, we performed 100 simulations to test whether random initial conditions led to self-organization of TWs. **(B)** Fraction of parameter sets that yielded at least one self-organized TW. **(C)** Distribution of reliability values among the set of parameters with reliability>0. **(D-E)** Reliability shows no clear dependence on the five parameters we varied. **(D)** Projection of the five-dimensional parameter sets on the space spanned by the two parameter specifying the strength of each interaction (see Fig. S3B for details). Each dot represents one parameter set and the color represents the reliability (color bar shared with Fig. S8E). **(E)** Spider chart projection. Each connected thread represents one parameter set, with the color of the thread representing the reliability for that parameter set.

## SUPPLEMENTARY MOVIES

**Video S1.** Self-organization of a traveling wave, corresponding to the trajectory depicted in Fig. 2A.

**Video S2.** The formation of a traveling wave follows a three-stage process, corresponding to the trajectory depicted in Fig. 5A.

**Video S3.** Dynamic spatial pattern with an oscillatory background, corresponding to the trajectory depicted in Fig. S3A.

**Video S4.** Macroscopic dynamics (fraction of ON-cells) of the trajectory shown in Fig. 5B.

**Video S5.** Explicit simulation corresponding to the trajectory shown in Fig. 5B and Video S4.

**Video S6.** Persistence of a traveling wave under the effect of stochasticity, corresponding to the example shown in Fig. 6C – top left.

**Video S7.** Formation of an oscillatory traveling wave with Hill coefficient 10, corresponding to the example shown in Fig. 6C – top right.

## Supplemental Information

### S1 Detailed description of the model

In this section, we give a detailed, mathematical description of the model under consideration. The aim here is to give a technically precise, but concise reference for the model details. As such, we leave out many steps in motivating the model assumptions - the reader can refer to the main text as well as our earlier work for a detailed motivation of the model assumptions ([1, 2]).

We consider a general model consisting of *N* cells that communicate through *l* distinct signaling molecules. The state of the system is specified by 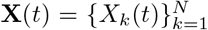, where 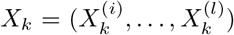 is the state of cell *k*. In the discussion below, we will distinguish between the *system state* **X** and the *cell state* of a single cell k, *X*_*k*_. Suppose that the cells secrete signaling molecules at a rate 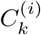, with bounds 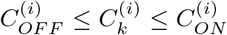. In principle the bounds can be different for each signaling molecule. However, in much of the analysis we will assume that 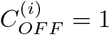 for all *i*. The secretion rate is related to the cell state through the relation

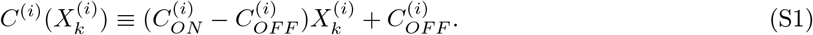

In the simplest scenario, the cells secrete signaling molecules at a rate which is either low or high. In this case the each of the 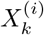 take binary values in {0,1}, such that 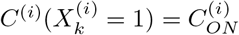 and 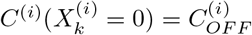. Alternatively, the secretion rate could take continuous values in 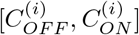. In this case the cell states are continuous and 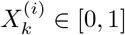. For convenience, we set 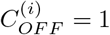 for all *i* and measure all concentrations in units of the OFF secretion rate (which we take to be equal for all genes, unless otherwise specified).

In steady state, the concentration of signaling molecule decays with distance to the cell that is secreting it according to [2]

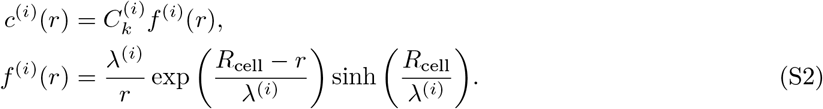

Here *R*_cell_ ≡ *r*_cell_*a*_0_ is the radius of the cell, *λ*^(*i*)^ is the signaling length and we introduced a function *f* (*r*) for the distance-dependent decay. Note that 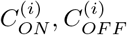 are effective secretion rates that lump together terms of the underlying reaction-diffusion equation ([2]), including the diffusion lengths *λ*^(*i*)^. However, we will not take into account this dependence and assume that the secretion rates scale accordingly with the *λ*^(*i*)^ such that 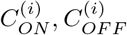 remain constant as *λ*^(*i*)^ is changed.

We can reduce the number of parameters by expressing all lengths in units of one length scale, for instance *a*_0_. We define 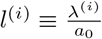 and write for the interaction function

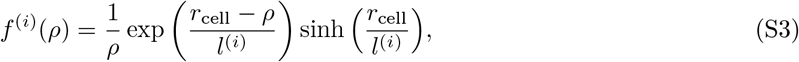

where we introduced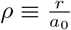

At any given time, the concentration that a cell senses is the sum of the concentrations due to each of the cells in the system. We express the sensed concentration of a cell *k* as

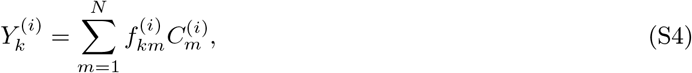

where 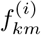 is a distance-dependent interaction strength between cells *k* and *m*. Explicitly,

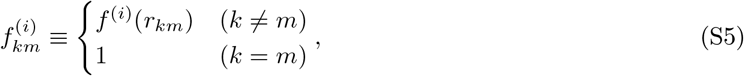

with *r*_*km*_ as the distance between cells *k* and *m* and *f*^(*i*)^(*r*) as defined in Eq. S2.

For later reference, introduce an interaction strength 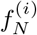 for each signaling molecule *i* as

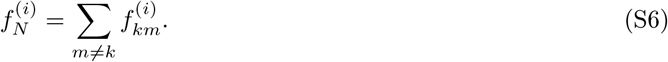

Note that if all cells secrete at the same rate *C*^(*i*)^, then they would all sense a concentration

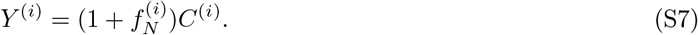

#### Regulatory interactions

Next, the effect of signaling molecule *j* on the production and secretion of signaling molecule *i* can be (1) positive, corresponding to activation of the gene responsible for producing *i*, (2) negative, corresponding to repression, (3) non-existent. To specify the interactions we introduce the interaction matrix *M*_*int*_, defined through

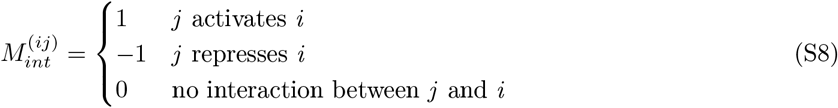

Note that the interaction network can be represented as a directed (multi)graph, where each node is a gene and each edge is a directed interaction between two genes.

The response of a cell can be a complicated function of the sensed concentrations of all signaling molecules, which depends on the biochemical details of the system. Here we consider a relatively simple case where the signaling molecules of both types need to simultaneously satisfy certain constraints to turn production on or off. In terms of logic functions, this corresponds to a regulatory construct where the two signals are put through an AND gate. For two activators, such a construct can easily be realized through employing cis-regulatory construct with weakly binding, adjacent sites for the two transcription factors, so that they can through some cooperative interaction, i.e. only bind when both factors are present ([3]). This scenario gives rise to a multiplicative update rule for our system. Namely, the effect of the different species multiply to determine a cell’s secretion rate at the next time step. Concretely, we have

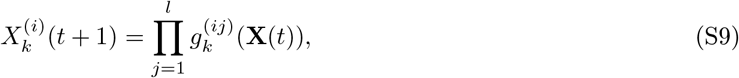

where

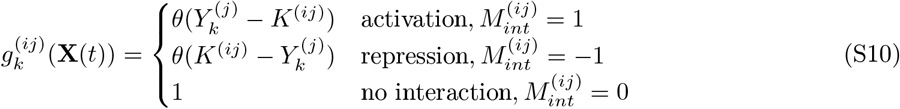

The result of molecule *j* regulating gene *i* is hence specified by *g*^(*ij*)^(**X**) as defined in Eq. S10. If *g*^(*ij*)^(**X**) = 1 then the interaction is ON, while *g*^(*ij*)^(**X**) = 0 means the interaction is OFF. Multiplicative interaction implies that whenever a single interaction is OFF, the gene will be turned OFF. However, if *g*^(*ij*)^(**X**) = 1 for some *j*, then we are not sure that gene *i* will be ON the next time step, unless *j* is the only regulating factor controlling gene *i*.

#### S1.1 Steady states of the system

In the discrete system with binary cell states, the number of states is finite (2^*N*^), so the system is bound to return reach a steady state after a finite number of time steps. In general, there are two possibilities:

1. Stationary steady state. The system reaches a stationary steady state, defined as a state for which **X**(*t* + 1) = **X**(*t*).
2. Periodic steady state. The system reaches a periodic steady state, with a period *τ*. There exists a time *t*\* after which we have **X**(*t* + *τ*) = **X**(*t*) for all *t* ≥ *t*\*.

The *equilibration time t*_*eq*_ is the time it takes to reach the steady state. For stationary steady states, this is simply the first time when the system reaches the steady state **X**(*t*_*eq*_). For all *t* > *t*_*eq*_, we have **X**(*t*) = **X**(*t*_*eq*_). For oscillatory steady states, we define the equilibration time as the onset time of the periodicity, e.g. *t*_*eq*_ = *t*\*, with *t*\* defined above.

#### S1.2 Counting topologies

For a two gene network, there are four possible interactions between the genes. Each interaction can be activating, repressing or absent. Hence there are 3^4^ = 81 possible topologies for systems with two genes. However, many of these topologies are equivalent through relabeling of genes 1 and 2. Under this operation, the interaction matrix maps to

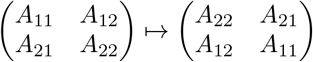

We see that the networks that are invariant under this operation satisfy *A*_11_ = *A*_22_ and *A*_12_ = *A*_21_. Since each of the two remaining independent variables can take three different values, there are 9 such networks. The remaining 72 networks can thus be reduced to a set of 36 unique networks. This gives a total of 45 distinct networks. After neglecting the trivial network with no interactions, we obtain the set of 44 networks shown in Fig. 3).

#### S1.3 Population-level description

To characterize the large-scale behavior of the system, we introduce macroscopic variables that characterize population-level features of the system. We characterize the macroscopic state of the system using two sets of parameters, with one for each gene. We first consider the average gene expression of the cells for a given gene *i*, which in the case of binary cells corresponds to the fraction of cells that have gene *i* turned ON:

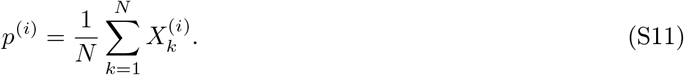

Next, we characterize how spatially correlated the gene expression levels using introduce an earlier defined “spatial index”, extended to a system with multiple genes [1, 2]:

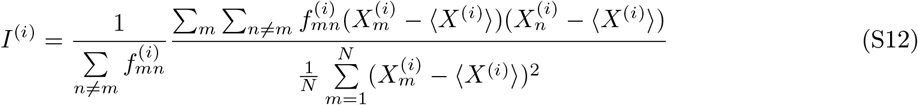

The quantity *I*^(*i*)^ is a measure for the spatial organization of the gene expression levels of different cells in the “channel” of gene *i*. It takes values between −1 and 1, with negative values indicating that neighboring cells tend to have different gene expression levels (such as in the case of checkerboard patterns or anti-ferromagnetism in spin models) and positive values corresponding to neighboring cells that tend to have similar levels of gene expression (forming islands with the same gene expression level, similar to the case of ferromagnetism). When *I*^(*i*)^ = 0, the expression levels of gene *i* are on average uncorrelated in space. In the case of spatially ordered patterns such as traveling waves, the values of *I*^(*i*)^ are positive and relatively high, with exact values depending on the parameters of the system and the shape of the wave.

Together, the set of macroscopic variables 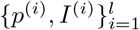 contain information about population-level features of each of the genes. However, note that this description does not contain information about correlations between different genes. For instance, we may specify for a two-gene system that *p*^(1)^ = *p*^(2)^ = 0.5, *I*^(1)^ = *I*^(2)^ = 0.5. This tells us that half of the genes of each type are on, and that the cells which have a certain gene on will tend to cluster with other cells that have the same gene on. However, it tells us nothing about whether a cell that has gene 1 on is likely to have gene 2 on, or whether its neighbors tend to have gene 2 on. The cells that have gene 1 on could be precisely those that also have gene 2 on, or they could be exactly its complement. There are different ways to consider metrics that also take into account cross-correlations. For instance, we can group together cells with each of the four cell states (i.e., (gene 1=ON, gene 2=ON), (ON, OFF), (OFF, ON), (OFF, OFF)) and study the evolution of these populations. However, the disadvantage of this approach is that it does not easily generalize to continuous descriptions of the gene expression states. Alternatively, we use established statistical metrics for correlations between two sets of values (the gene expression levels for the two different genes) such as the *Hamming distance*, the *Jaccard index* (JI) and the *Sørensen-Dice coefficient*. As we are mainly interested in knowing whether a configuration is spatially ordered or not (i.e. whether it may be an interesting pattern), we have not looked in detail at these cross-correlations.

#### Moving averages

The moving coefficient of variation shown in Figs. 5C and S4 is obtain by dividing the moving variance by the moving mean, both with a window size of 10. Specifically, we calculated the moving variance using the MATLAB function `movvar` and the moving mean using `movmean`. F

### S2 Simulation and analysis of the model

In this section, we provide a basic overview of parts of the simulations and analyses which may not be evident to the reader. This is by no means a comprehensive overview, for which we refer the reader to the source code of the simulation software.

#### Fixing initial conditions

We start the simulation by generating an random initial configuration subject to certain constraints. If no initial conditions were specified, we let each gene of each cell be ON with a probability 1/2. This tends to generate configurations where half of the cells of each type are ON, whereas configurations where all the genes are OFF or ON are rare generated (since the statistics follows a geometric distribution). Alternatively, we could fix the macroscopic variables of the system (Sec. S1.3). To fix *p*^(*i*)^, we randomly select this fraction of cells, for which we turn ON gene *i*. To fix *I*^(*i*)^, we used a Monte Carlo algorithm which we will outline below.

#### Algorithm for generating configurations with given spatial index

We constructed an algorithm that generated configurations with a given spatial index *I*^(*i*)^ at fixed values of *p*^(*i*)^. This is analogous to a similar problem for spin systems in physics, where we try to fix the energy of the Ising model (analogous to *I*^(*i*)^) without changing the average magnetization of the system (analogous to *p*). The algorithm is illustrated in Fig. S1 and can be summarized as follows:

1. Given an configuration with a given value of *p*, start by computing the value of the *I* for this configuration.
2. Check whether we should increase or decrease *I* by comparing it to the target value *I*_*target*_.
3. If *I < I*_*target*_

a. Select the ON-cell with the *minimum* number of neighbors which are also ON. Turn this cell OFF.
b. Select the OFF-cell with the *maximum* number of neighbors which are also OFF. Turn this cell ON.
4. Else if *I < I*_*target*_

a. Select the ON-cell with the *maximum* number of neighbors which are also ON. Turn this cell OFF.
b. Select the OFF-cell with the *minimum* number of neighbors which are also OFF. Turn this cell ON.
5. Compute the the spatial index of the new configuration, *I*_*new*_. Check whether it has increased or decreased as required.
6. If it has changed as required, accept the change. Go to step 8.
7. Else, reject the new configuration. Go to step 1.
8. If *I*_*new*_ ∈ [*I*_*target*_ − *ϵ*, *I*_*target*_ + *ϵ*], terminate the simulation.
9. Else, go to step 1 with the new configuration with *I* = *I*_*new*_.

Because we switch the state of both an ON-cell and an OFF-cell, the average number of ON cells is kept constant. To increase *I*, we choose cells that tend to have a different state from most of their neighbors and change their state. To decrease *I*, we change cells that tend to have a similar state to their neighbors. In practice, we typically set *ϵ* = 0.01. Note that this algorithm is not guaranteed to converge to *I*_*target*_ because at each iteration of the loop, we are not guaranteed to increase or decrease *I* as required. In particular, if the specified *I* is outside the range of possible *I* [2], the algorithm cannot reach the specified value of *I*. Therefore, we typically set a limit on the maximum number of iterations before we terminate the algorithm.

**Figure S1:**
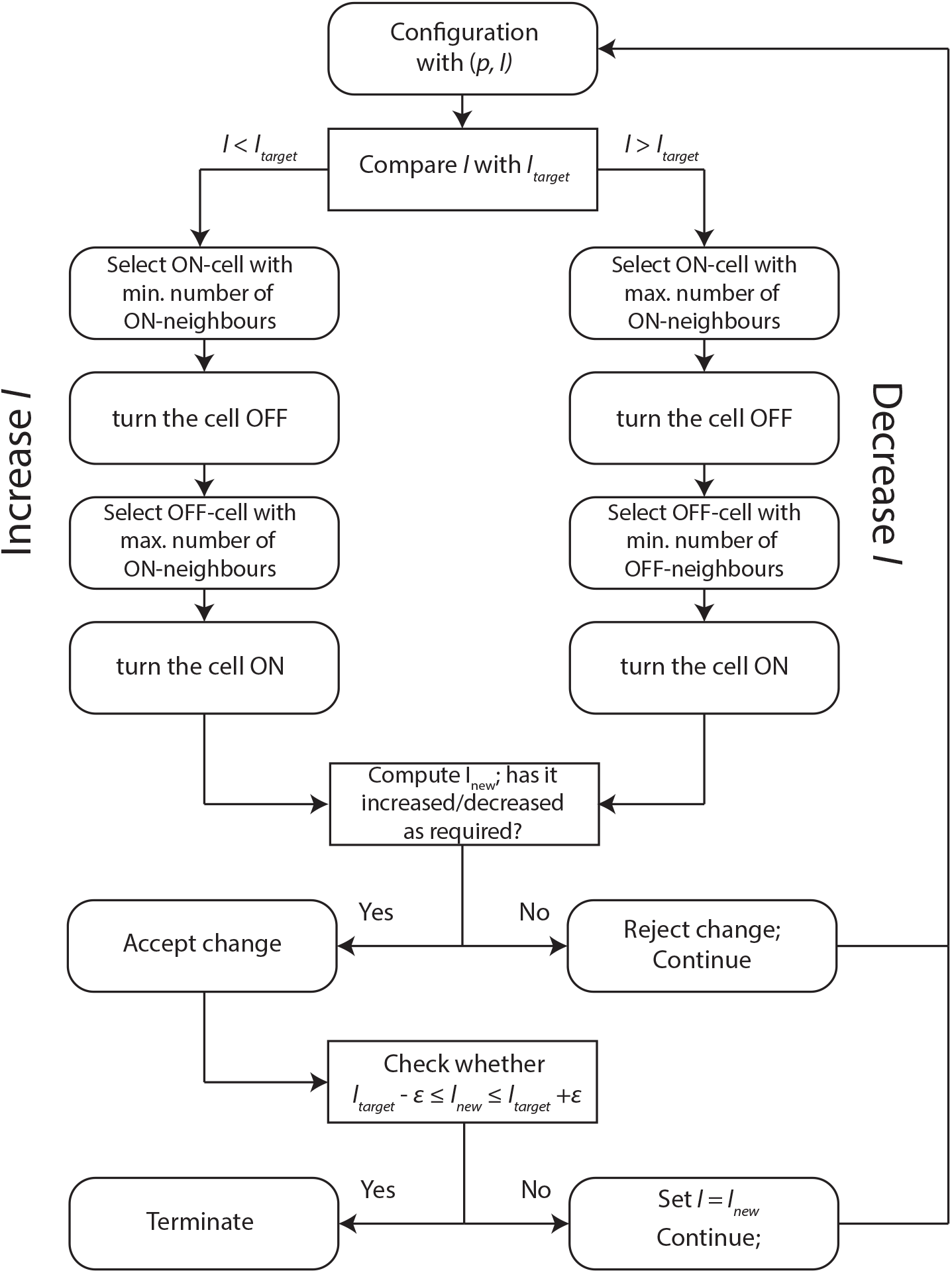
Work flow of the algorithm used to generate configurations with specific *I* at a fixed value of *p*.

#### Terminating the simulation

As noted earlier, we terminate the simulation whenever it has reached a steady state or when it reaches a maximum number of time steps *t*_*max*_ (in practice, we chose *t*_*max*_ = 10^5^ for systems of size *N* = 225). At each time step, we checked for stationary steady states by comparing the current system state with the system state at the previous time step. To check for periodicity, one method would be to brute-force check at every time step whether the current state has been visited earlier. This becomes computationally intensive when running large sets of simulations, so we devised a more efficient algorithm for checking periodicity. Instead of checking at every time step, we check every *t*_*check*_ time steps whether the last state has been visited earlier by brute force (in practice we set *t*_*check*_ = 10^3^). If we find periodicity, we run a second algorithm to find the earliest time at which any state has repeated itself, which gives us the onset time of the periodicity.

#### Identifying traveling waves

We devised an algorithm to automatically identify TWs in large sets of simulations. Since the waves keep their shape while translating across the lattice, their values of *p*^(*i*)^ and *I*^(*i*)^ do not change over time. Because of periodic boundary conditions, they reach the same state again after *n* time steps, where 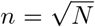 is the grid size (linear size of the system). Hence, we first filter the trajectories on ones that have a period which is a multiple of *n* (since we may also have more complicated waves that are slightly morphed as they travel across the boundaries). We then check whether *p*^(*i*)^ and *I*^(*i*)^ and (sufficiently) constant over one period of the wave. Using these two features, we could identify traveling waves in batch simulations without examining the simulations explicitly by eye. We then extended the algorithm such that it also gives the orientation of the wave of a TW forms (relevant for Section S6.5 and Fig. S6C). This is done in a two-step procedure: first we separate the background state from the wave states (assuming there are three states that makes up a wave), then we trace the cells of an arbitrary wave state layer to see whether they percolate the system from one horizontal (vertical) edge to the other. If so, then we assign a horizontal (vertical) orientation to the wave. Else, we assign a diagonal orientation to the wave.

#### Batch simulations

Many of the results presented in the figures and paper rely on batch simulations, where we performed a large set of simulations with similar but not identical conditions (parameters and initial states) to obtain overall statistics on various measures. In many cases, we fix all parameters of the system and only vary the initial conditions. By performing a large set of simulations with the same parameters but initial conditions, we can distinguish whether an observed feature is consistent for that set of parameters or is an artefact of the specific initial conditions. However, in a number of cases we have also considered the effect of varying parameter sets. In particular, we addressed the question of what a system with a number of fixed parameters (e.g. fixed cellular dialogue) is capable of in general, as one varies the parameters across some set range. Because the parameters vary continuously, we cannot simulate all possible parameter values and have to find a way to sample over this space. One particular method we used is *Latin hypercube sampling* [4], where we efficiently sample over a multi-dimensional parameter space by taking parameter sets that are non-overlapping in any of the dimensions. We used this method to initially generate a large set of simulations for each of the 44 networks to explore their range dynamic behavior - the results in Figs. 3, S2 and rely on this approach. Specifically, for this data set we defined a region where we varied the parameters *K*^(*ij*)^ and 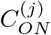 over a range from 1 to 10^3^ and kept all over other parameters constant. We sampled over this region by generating a Latin hypercube sample with 10,000 points using the MATLAB function lhsdesign. We did this for each of the 44 cellular dialogues.

### S3 Overview of self-organized patterns

In this section, we provide a detailed but mostly qualitative overview of the different types of self-organized patterns we observed in simulations of our model. The aim is give a general overview of the different possible morphologies and dynamic features of these patterns, and to understand basic features of these patterns in terms of the concepts we have introduced or will introduce. In Section S5, we will give analyze traveling waves - a subset of dynamic spatial patterns - more closely.

#### S3.1 Static patterns

While the focus of this work is mainly on dynamic patterns, we also observed static patterns with a high degree of spatial organization in all two-gene networks studied. They commonly arise after a relatively short transient phase (10s-100s of time steps) and can have different shapes and compositions of cell states (i.e. the cells can have different combinations of gene expression). In terms of shape, most patterns consist of one or more islands or stripes of cells with a different cell state from the surrounding cells. Since there is no Turing mechanism in our system, we did not identify a natural length scale for the patterns, although changing parameters did seem to affect the size of structures such as islands. Patterns were most commonly observed to have two sets of cell states, where one group of cells has one cell state and the other group has another cell state. Patterns with three cell states are rare, but not impossible to generate. We did not observe any patterns where all four cell states existed concurrently. The most common static patterns are ones that also arise in the model with one signaling molecule and consist of one group of cells with a given gene ON and another group with that gene OFF. In the case of two molecules, it is common to find islands with both genes ON (or OFF) with the rest of the system consisting of cells with both genes OFF (or ON). We also observed patterns where the two genes were mutually exclusive, i.e. if a cell has gene 1 ON, it has gene 2 OFF, and vice versa. Finally, we found that the boundary separating an island or stripe from the rest of the cells can sometimes take a different state from the rest of the system.

#### S3.2 Dynamic temporal patterns

Dynamic temporal patterns are oscillatory steady states are steady states where the system returns to an earlier state after a finite number of time steps *τ* > 1 (the period of the oscillation), but do not propagate information across space.

##### Single-cell oscillations

Oscillations can arise at the single-cell level in the case of a negative feedback loop. If certain parameter constraints are satisfied, the gene expression level of a single cell would oscillate between ON and OFF indefinitely. These oscillations are the result of our adiabatic description, where we assume that cells respond slowly compared to the time for signaling molecule concentrations to reach a steady state. Cells oscillate between the ON and OFF states due to its slow and binary response. An ON-cell turns OFF because the concentration it senses is high enough to suddenly switch to the other state. The OFF-cell then senses a low concentration and the cell switches to the other extreme immediately, without ever reaching the intermediate steady state.

With two genes, oscillations at the single cell level remain relatively simple and can only have periods up to four, since there are only four cell states with two genes. In practice, by examining all possible single-cell state diagrams (Sec. S4; not described in detail), we found that the vast majority of single-cell oscillations were of period 2 (see Fig. S2C). Single-cell period 2 oscillations arise in all networks that can generate dynamic patterns (temporal or spatial), while period 4 oscillations arise only in networks with an incoherent mutual feedback (i.e., for all networks generating dynamic spatial patterns in Fig. 3D as well as Network 14 in Fig. 3C).

We can interpret these results by looking at the three core network structure that give rise to most dynamic patterns (Table 1). For each of the motifs, the interpretation of the oscillations is straightforward. For a double repression, a cell is able to oscillate between (0, 0) and (1, 1), whereas (0, 1) and (1, 0) are stable states. When both genes are off, both are unrepressed and will turn on the next time step, after which they are both repressed and turn off again. However, if only one of the genes is on, it represses the other gene but is not repressed itself. For a double activation, the oscillation is between the states (0, 1) and (1, 0). Each gene can turn on the other, but turns off when the other gene is on. However, if both genes are ON, they sustained each other, whereas if neither is ON, they also cannot turn ON. Finally, for a positive-negative loop, the system undergoes a period 4 oscillation between the four states as one may see from going through all states one by one. These results obviously depend on the parameters chosen, but it is evident that they should be possible for some set of parameters. Again, these results rely on the separation of time scales between the relaxation of the signaling molecule concentrations and the response of the cells. They persist on a multicellular level, with cells synchronizing their oscillations depending on how strongly the cells interact. Oscillatory cells can co-exist with stationary cells that are in one of the stationary states (for negative-negative or positive-positive feedback).

**Table 1:**
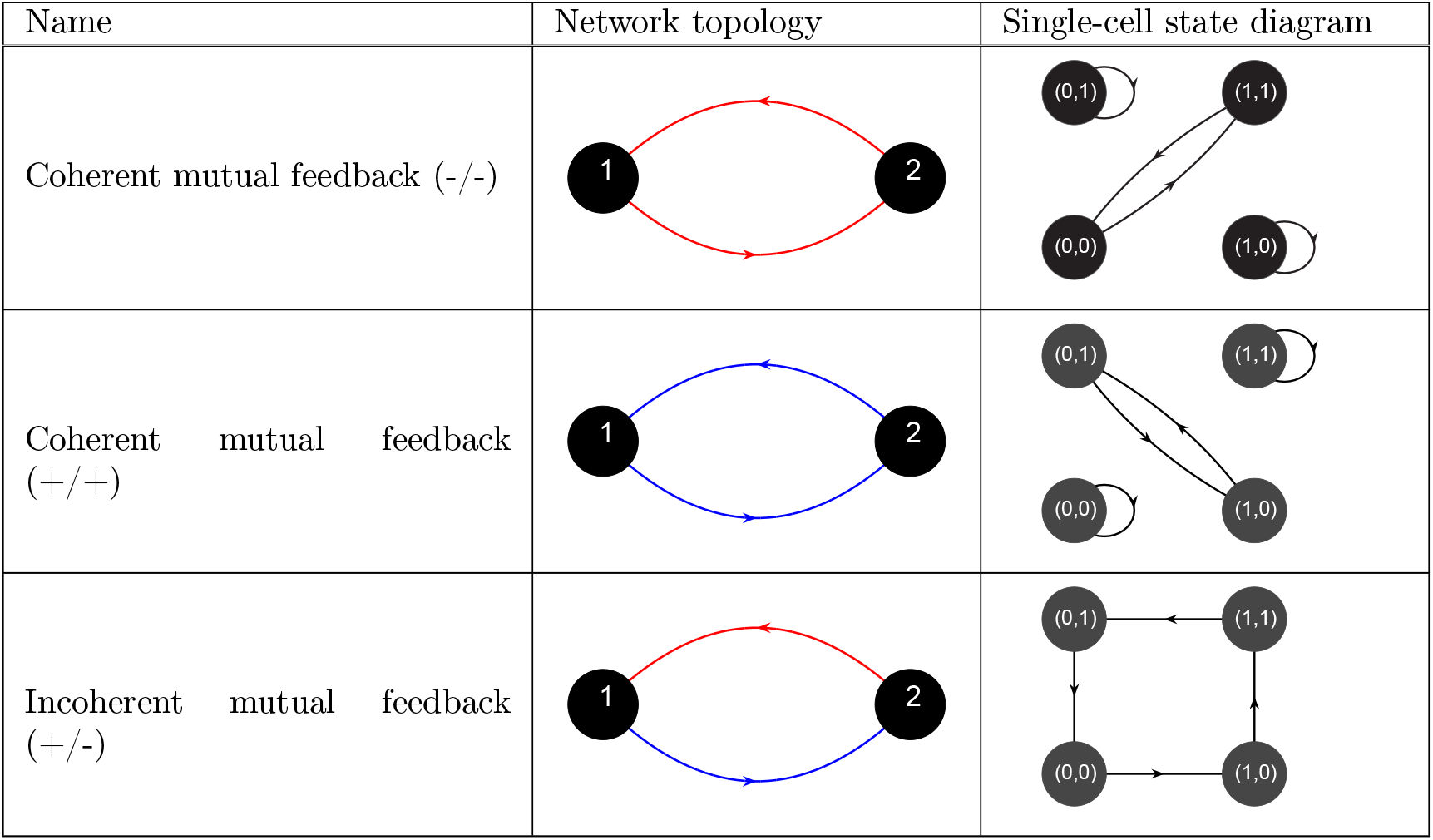
Two gene network motifs generating oscillations. The three core topologies for mutual interaction between the two genes are shown together with typical single-cell state diagrams showing oscillations. The state diagrams are for the case when the concentration of the regulator genes always surpass the threshold when the gene is ON and is below the threshold when it is OFF, i.e. 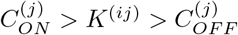 for all genes *i, j*.

##### Synchronization of single-cell oscillations

On the multicellular level, oscillations can persist in different ways, depending on the degree of synchronization and whether the oscillations are “simple”, i.e. superpositions of single-cell oscillations. The oscillations in the multicellular system can vary between completely autonomous (i.e., each cell independently oscillates) to completely synchronized (i.e., all cells oscillate synchronously; see Fig. 2I). Full autonomy is reached if and only if each of the interactions is in the autonomous (A01) phase (see Sec. S4). Full synchronization can be reached for a variety of other parameter conditions. In between, the system can partially synchronize and exhibit domains of cells oscillating together that do not extend over the entire lattice.

##### Complex dynamic temporal patterns

Oscillations of a more complicated form arise in the networks that are capable of generating dynamic spatial patterns. Each of these networks produces a wide range of complicated periods with *τ* ≥ 5 (Fig. S2D), most of which correspond to dynamic temporal patterns. Typically, they feature oscillating domains that coexist with a background of static cells (e.g., the oscillating island in Fig. 2H), but where different cells in the domain undergo different cycles of gene expression states over time. This can give rise to complicated temporal patterns because the oscillatory sequences of individual cells may not line up, especially when they are incommensurable.

These complex periods are indeed associated with dynamic patterns, as we can verify by measuring their degree of spatial order using the metric *I*^(*i*)^ (Eq. S12). Overall, steady states with a period *τ* ≥ 5 tend to have higher values of *I*^(*i*)^ for both genes (yellow and purple lines in Fig. S2E), indicating that they tend to be more spatially ordered than oscillations with simple periods (red lines in Fig. S2E) and static patterns (blue lines in Fig. S2E).

#### S3.3 Dynamic spatial patterns

Dynamic spatial patterns are characterized by gene expression profiles that translate across space, thereby allowing propagation of information across the multicellular system. These can be rigid profiles of gene expression that move across the system without changing shape, but we also count patterns that move and morph (i.e. change shape) at the same time as dynamic spatial patterns. Note that these patterns require periodic boundary conditions to be sustained indefinitely.

##### Traveling waves

Traveling waves are characterized by stripes of cells that translate across the lattice in a regular fashion (see Fig. 2A, 2C-E and 2G for examples). They typically consist of three types of cells (with different states) and travels on a background consisting of cells of the fourth type. When two traveling waves in opposite directions collide, they typically annihilate each other, leaving a void of cells with the background state. Characteristics of traveling waves and their propagation conditions will be discussed in Section S5.

##### Complex wavelets

In a number of cases, we observed complex wavelets that propagate indefinitely without repeating themselves, within the maximum simulation time (Fig. 2B). Since the total number of system states is limited to 2^*N*^, these waves will eventually settle down to a steady state. The transient wave patterns they generate look very similar to (less coordinated) traveling waves, and arise as transient states during the generation of all types of dynamic spatial patterns described here.

##### Spiral and concentric waves

Spiral and concentric waves are similar to traveling waves with the main difference that their orientation is outward from a source or center rather than linear in a fixed direction (Fig. 2F). Locally, they typically look like traveling waves, with the same set of cell states as in traveling waves. Due to annihilation of colliding waves, spiral and concentric waves are less stable, since only particular configurations where the outcome of the collision is an earlier spiral wave pattern will be observed as persistent spiral waves. It is more common to observe spirals and concentric waves as transient patterns that are created and annihilated repeatedly, until the system settles down to a more stable configuration such as a traveling wave.

##### Traveling pulses

We also observe small, localized patterns of a few cells that translate across the lattice in a regular way. They are similar to traveling waves, but the traveling pulses are rather small, localized patterns that do not span the entire size of the system.

##### Oscillatory traveling waves

In networks 16, 20 and 43 - characterized by the incoherent mutual feedback motif without positive self-regulations (Fig. 3D) - we found oscillatory traveling waves where both the wave states and the background state oscillate over time (Examples in Fig. S1). At any given time, these waves typically look similar to the non-oscillatory traveling wave, but due the oscillations the dynamics is different. Perfectly aligned waves where each wave state occupies a single band of cells are relatively rare. Most waves have bands that occupy the width of more than one cell (see for instance Fig. S1B). The waves undergo a successive sequence of static oscillations followed by an translation (Fig. S1D). Details of their dynamics are further discussed in Sec. S5.6.

### S4 Parameter-derived general constraints on the dynamics

In this section, we derive a number of a methods to derive general constraints on the dynamics of the system from the parameters of that system. First, we introduce the concept of dynamical phase for each regulatory interaction between two (possibly identical) signaling molecules. Next, we introduce the concept of state diagram - a graphical way to represent transitions between cell states - and discuss their usefulness in deducing constraints on the system’s dynamics. We then derive general constraints on the dynamics of a system with multiple signaling molecules, which arise as special combinations of these phases. Finally, we present a formal algorithm to calculate the dynamical constraints and represent them in a state diagram for any set of arbitrary parameters.

#### S4.1 Dynamical phases for each interaction

The idea behind dynamical phases is based on the observation that for extreme parameter values, the behavior of the system becomes predictable. For instance, if the interaction between cells is very strong and the threshold values characterizing their response very low, then we expect the cells to always exceed this threshold regardless of the precise states of the cells. These ideas were made precise in our previous work for systems with one signaling molecule, which represented these dynamical constraints as “phenotype functions” on a “phenotype diagram” [1]. In this work, we extend this formalism to a more general framework applicable to multiple interactions.

Consider an interaction between two genes where a regulating gene *j* controls the expression of the regulated gene *i* (possibly *i* = *j*). The interaction is specified by the threshold *K*^(*ij*)^ for turning the gene ON/OFF (depending on whether the interaction is activating or repressive), and the ON-secretion rate 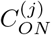. Suppose the cells have an effective distance *a*_0_ to their nearest neighbors, and we associate an interaction strength 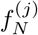 (Eq. S6) to the interaction. Let *Y* ^(*j*)^ be the sensed concentration of molecule *j*. Recall that the outcome of the interaction is specified by *g*^(*ij*)^(**X**). We then distinguish the following phases:

1. **P1**: sensed concentration permanently above threshold. The phase is defined by 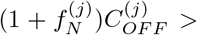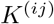. For an activating interaction, this implies that *g*^(*ij*)^(**X**) = 1 for any system state **X**. The interaction is always ON. For a repressive interaction, we have *g*^(*ij*)^(**X**) = 0 and the interaction is always OFF. For a single activating interaction, this corresponds to the always ON phase.
2. **P0**: sensed concentration permanently below threshold. The phase is defined by 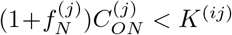. For an activating interaction this implies that *g*^(*ij*)^(**X**) = 0 and for a repressive interaction *g*^(*ij*)^(**X**) = 1. For a single activating interaction, this corresponds to the always OFF phase.
3. **A1**: autonomy whenever the regulating gene is ON. The phase is defined by 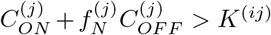. For an activating interaction, this implies that 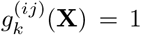 whenever 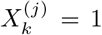. This means that the interaction is always ON in a cell *k* whenever gene *j* is ON regardless of the rest of the cells. However, when gene *j* is OFF, whether the interaction will be ON depends on the state of other cells. For repression, 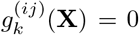 whenever 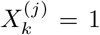. In this case, the interaction is always OFF in cell *k* whenever it has gene *j* ON. For a single activating interaction, this corresponds to the activation phase.
4. **A0**: autonomy whenever the regulating gene is OFF. The phase is defined by 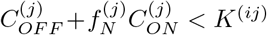. This is analogous to the previous case with some roles switches. For activation, we get 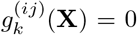 whenever 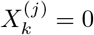. For repression, 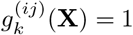 whenever 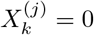. For a single activating interaction, this corresponds to the deactivation phase.
5. **A01**: autonomy regardless of whether the regulating gene is ON or OFF. This phase is defined by parameter values for which both inequalities of A1 and A0 hold. These conditions can only be met simultaneously if 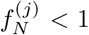. For activation, it implies that 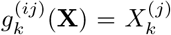. More explicitly, it means that 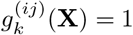 whenever 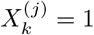 and 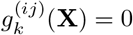 whenever 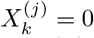. Hence 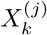 fully determines fate of the interaction. For repression, the roles are reversed and 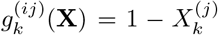. For a single activating interaction, this corresponds to the autonomy phase.
6. **U**: unconstrained. This phase is defined by parameter values for which the conditions of neither A0 nor A1 are true. Hence, we have 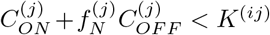 and 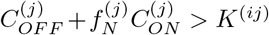. These conditions can only be met simultaneously if 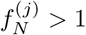. For a single activating interaction, this corresponds to the activation-deactivation phase. In this phase, we cannot deduce any general constraints on 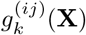 and have to look at the specific system state **X** to determine whether an interaction will be ON or OFF.

The interpretation of these phases are best understood for a system with only one signaling molecule [1]. For now, note that the phases **P0** and **P1** make the interaction trivial - the outcome is always known regardless of the state of the cell itself or its neighbors. The phases **A0**, **A1** and **A01** place constraints which are dependent on the current state of the cell, and the phase **U** does not place any constraints on the system’s dynamics. For a system with multiple signaling molecules, each interaction will be characterized by one phase. The obvious next question is how to put together the constraints from these different interactions to derive general constraints on the system’s dynamics.

#### S4.2 State diagrams

The basic idea of the state diagram is that it displays all the possible transitions between different cell states of a system. The concept of state diagram has been explored in earlier work on modeling genetic circuits with binary expression states [5], but has been limited to models of gene circuits at the single-cell level. In our case, the cell states are the binary states *X* = (*X*^(1)^, …, *X*^(*l*)^) ∈ {0, 1}^*l*^ specifying for a given cell whether each gene is ON or OFF. For one signaling molecule, there are only two cell states, 0 and 1. For two signaling molecules, we distinguish the four cell states (0, 0), (0, 1), (1, 0) and (1, 1). If a cell in a given cell state *X* can adopt the state *Y* after one time step - either under its own influence or through sensing molecules secreted by other cells - then we draw an arrow between the states *X* and *Y*.

For a single interaction, the procedure is straightforward and the diagrams are simple to interpret. For instance, the P1 phase with activation means that cells always turn ON after one time step, hence giving the diagram with all arrows going to the 1 state and no other arrows. For A01 with an activating interaction, the cells are autonomous, so 0 remains 0 and 1 remains 1. However, A01 with repressive interaction gives oscillations between 0 and 1. An ON-cell always turns OFF because it will always sense a concentration above the threshold, repressing its gene expression, while an OFF-cell will always turn ON because its sensed concentration will always be below the threshold, leaving the gene unrepressed. The collective set of possible transitions is then displayed as a diagram with the cell states and possible transitions.

For two genes, the interpretation is analogous. We draw a directed graph with the four cell states (0, 0), (0, 1), (1, 0) and (1, 1) as nodes and directed edges between these states to indicate (possible) transitions between the states. In Fig. 4E, in the left diagram for dialogue 14, the transitions are heavily constrained. Each of the four states leads only to one other possible state. This completely constrains the dynamics of the system, so that it becomes completely predictable - any cell’s dynamics in the system is completely determined by the transitions in one diagram. As such, this state diagram does not allow for multicellular pattern formation, as all individual cells will oscillate individually. In contrast, if we take a diagram such as the one depicted for dialogue 15 (Fig. 4E), the dynamics of the system is not entirely constrained. The state (0, 0) has two arrows leaving from it, indicating that either transition is possible, depending on the exact concentration a cell senses. As such, it allows for a pattern such as a traveling wave to propagate, because cells of the same state do not always evolve in the same way, but this depends on the other cells in the system. In practice, the state diagram of dialogue 14 (Fig. 4E) can be realized under a range of parameter conditions, whereas the diagrams for dialogues 15 and 19 (Fig. 4E) typically involve more possible transitions (which does impede TW propagation).

Two properties of the system are immediately evident from the graphical representation of the state diagram. To begin with, a state diagram tells us which cell states could potentially be stationary. Such states must have an arrow to themselves in the state diagram, which we call a *self-transition*. Should the system reach a non-oscillatory steady state, then that final state can only be composed of cell states which have a self-transition. If there are no self-transitions, then the system cannot generate stationary patterns - this happens for instance in dialogue 14 with certain parameters, which can produce the state diagram shown in Fig. 4E. If there is only self-transition, then the only possible stationary steady state is a uniform system where all cells have the state with the self-transition. Conversely, not all cell states with a self-transition need to appear in a stationary system state. In other words, having a self-transition does not imply that the state appears in any stationary pattern. As an extreme example, the system could have a fully connected state diagram, where each transition between two cell states is in principle possible. However, this system could still generate a uniform lattice of cells as final state if the parameters are chosen appropriately.

Secondly, the state diagrams show whether periodic steady states (e.g. oscillations) are possible. For any periodic steady state, all cells must revisit their earlier state after *τ* > 1 number of time steps, where *τ* is the period of the oscillatory state. This implies that the state diagram should permit cells to return to their initial states after a finite number of time steps, and after passing through other states (otherwise it would be a stationary pattern). This is only possible if the state diagram contains cycles, i.e. closed loops obtained by tracing the edges of the graph from some initial state. The presence of cycles is a necessary condition for oscillations. Conversely, the presence of a cycle is not a sufficient condition for generating dynamic temporal patterns. This is because it is not guaranteed that a cell can traverse the edges of any cycle one by one when there are possible “routes” on the graph. Each transition then corresponds to a specific condition which depends on the state of all cells of the system. We cannot directly deduce whether a sequence of such transitions is possible at the level of the entire system of *N* cells. However, in the special case that all transitions of the cycle are deterministic, i.e. when each node is connected to a unique other node on the graph, we obtain an oscillation. The length of the cycle then corresponds to the period of the oscillation.

In summary, the state diagram allows us to deduce two basic properties from first inspection: the set of stationary cell states and the possibility of finding oscillatory steady system states. These are not purely mathematical properties but have biological relevance. The former is an indicator of multistability and tells us whether a population of identical cells could potentially diversify, generating stable configurations with multiple gene expression profiles. This is known as *phenotypic heterogeneity* and has been observed in many experimental systems [6]. The latter tells us whether a multicellular system could potentially sustain oscillations (consult the main text for biological examples).

#### S4.3 Simplified dynamics

There are two limits in which the dynamics of the system simplifies dramatically.

1. All interactions are either extremely weak or extremely strong. To be precise, this is the case if each of the interactions is in the P0 or the P1 phase (all ON/all OFF phase). The system homogenizes after one time step, because each of the interactions is either ON or OFF for all cells in the system. For a spatially uniform system, the dynamics is simple and predictable.
2. All interactions are moderately strong, and the interaction between cells is relatively weak. In more precise terms, suppose all interactions are in the A01 phase. In this case the dynamics of each cell becomes equivalent to that of a single cell. The system is fully autonomous and each cell evolves under its own influence.

In these limits the state diagrams are identical to single-cell state diagrams with rescaled parameters. Therefore, these phases contain only deterministic state diagrams. Thus we know the exact dynamics of the system without running any simulations, for any initial conditions.

##### Formal derivation

Let us show these two limits more explicitly. Let *k* be arbitrary, and (*i, j*) be an arbitrary pair with 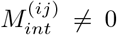. Suppose this interaction is in the P0 phase. Then as a result of 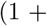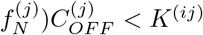, we have

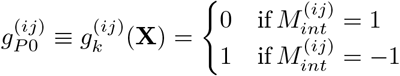

That is to say, 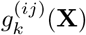 becomes independent of both *X* and *k*. Likewise, in the P1 phase we get 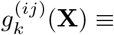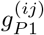, with 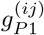 depending only on 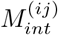. Now suppose all interactions are either in the P0 or P1 phase. Then, 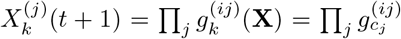 with *c*_*j*_ ∈ {*P*0, *P*1}. Hence, 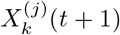 is also independent of both *X* and *k*. Therefore, all cells become identical after one time step. For an identical lattice with all cells in a state *X*, we note that 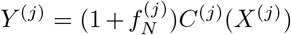. However, *K*^(*ij*)^ and other parameters are unchanged. Therefore, the evolution of a uniform lattice is equivalent to that of a single cell with a rescaled secretion rate 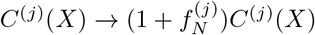.

Next, consider the case that all interactions are in the A01 phase. Again, let *k* and (*i, j*) be arbitrary with 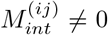. Let **X** be an arbitrary state of the system, and write **X** = (*X*_*k*_, *Z*_*k*_), with *Z*_*k*_ = {*X*_*l*_}_*l*≠*k*_. Then the A01 phase puts the following constraints on the system:

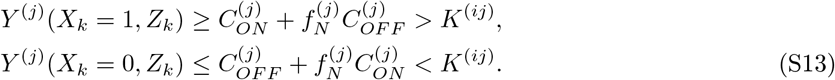

We see that any cell with gene *j* ON will always satisfy the first constraint, regardless of the rest of the system. Likewise, any cell with gene *j* OFF will always satisfy the second constraint. As a result, *g*^(*ij*)^(*X*_*k*_; *Z*_*k*_) = *g*^(*ij*)^(*X*_*k*_) becomes independent of *Z*_*k*_, the states of all cells other than *k* in the system. Therefore, 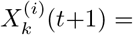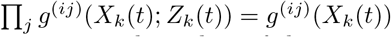 depends only on *X*_*k*_(*t*). In other words, the evolution of any cell in the system is independent of the state of the other cells.

#### S4.4 Algorithm for computing state diagrams

In this section, we present a general method for computing the state diagram for a system of one or two genes, given an arbitrary set of system parameters. The construction for two genes can be readily generalized to systems with more than two genes.

For a single gene, we state diagrams follow straightforwardly from the definition of the phases (Section S4.1). The end result can be represented as a directed graph with two nodes and up to four edges, which we can describe using its adjacency matrix

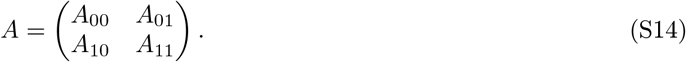

The adjacency matrix gives information on whether edges are present for each potential link between two nodes. The entries *A*_*ij*_ ∈ 0, 1 are for connections between node *i* and node *j*. If *A_ij_* = 1, there is an edge between the two nodes, if *A*_*ij*_ = 0 there is no edge.

As an example, consider cells with a single signaling molecule with negative feedback to itself. The graphs for positive feedback are deduced in a similar way. In the P1 phase, the system is permanently repressed, so all states go to the 0 state. Hence 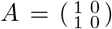. By contrast, in the P0 state, both ON and OFF cells always turn ON at the next time step, so 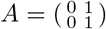. In the A1 state, ON cells always turn OFF, but we do not know anything about the OFF cells. Hence both transitions 0 → 0 and 0 → 1 are possible. Therefore, 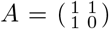. Conversely, in the A0 phase, only OFF cells are constrained to always turn on, so 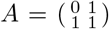. In the A01 phase, OFF cells turn ON and ON cells turn OFF, so 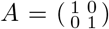. Finally, in the U phase, all transitions are unconstrained, so 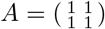.

For systems with two or more genes, the procedure is considerably more involved. We first outline the intuitive idea behind this derivation and then provide a formal, mathematical derivation of the construction. With two mutually interacting signaling molecules, the dynamics of a gene *i* depends in general on both regulation by itself and regulation by the other gene, which we label *j*. If we know the phases of both regulations *i* ← *i* and *i* ← *j*, then we can deduce the constraints they impose on the dynamics of *i*. To do this, we have to combine the constraints from both regulatory interactions *i* ← *i* and *i* ← *j*, for which we employ a three-valued logic operation. Intuitively, this three-valued logic system represents the fact that there are three possible outcomes of each interaction: the regulated gene is activated, the regulated gene is repressed or the outcome is unknown. Hence, we need to know what the final response of gene *i* is for each combination of the three outcomes that each of the two regulatory interactions can have. For instance, suppose that both *i* and *j* positively regulate *i* (i.e. 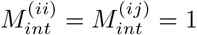), but the interaction *i* ← *i* is always activating (i.e., the sensed concentration of *i* always exceeds the threshold *K*^(*ii*)^) while the interaction *i* ← *j* is unknown. Then the final outcome for gene *i* is unknown, because both positive interactions must be activating for the gene to turn on. Next, recall that the phases in general place dynamical constraints that depend on the state of the system. Concretely, this means that the constraint placed by *i* on itself depends on whether it is ON or OFF, and the same holds for the constraint placed by gene *j*. Therefore, for each combination of states for genes *i* and *j* - there are four of these, corresponding to the four cell states (0, 0), (0, 1), (1, 0) and (1, 1) - we could have a different set of constraints on the dynamics for gene *i*. Thus, we have to separately consider each of the four cell states and see which constraints they impose on the dynamics of both gene *i* and gene *j*. In this way, for each cell state, we obtain all possible cell states to which it can transition to and draw the corresponding edges on the graph. This eventually gives us our state diagram.

##### Formal derivation of the algorithm for computing state diagrams

First, we note that the four-node graph with up to 16 edges is now represented by a 4 *×* 4 adjacency matrix, which we will denote

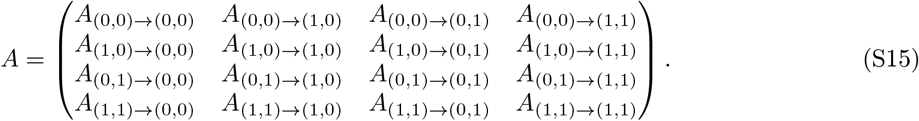

The interpretation is the same: the state (*i, j*) can transition into the state (*k, l*) if and only if *A*_(*i,j*)→(*k,l*)_ = 1, and we represent this graphically by drawing a directed edge between (*i, j*) and (*k, l*). Our goal is then to combine the constraints imposed by the different phases for each interaction to compute this adjacency matrix.

Recall that the time evolution for a cell determined by *X*^(*i*)^(*t*+1) = Π*g*^*ij*^(**X**(*t*)) (S9). This is a deterministic equation for *X*^*i*^(*t*) for when we know the precise input system state **X**(*t*). Now suppose we only know the cell’s own state *X* = (*X*^(1)^, *X*^(2)^) and the phase of each interaction *i* ← *j*. We want to calculate the set of *possible* output cell states for *X*(*t*+1). To do this, we introduce a set of three-valued logic states *S* = {0, 1, 2} and a logic AND function ∧ : *S* → *S* defined by the truth table

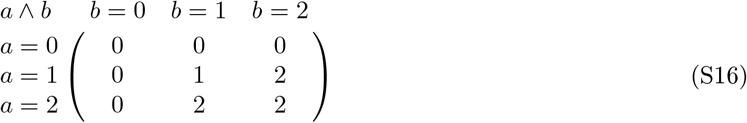

This 3-valued logic system is known as the *Kleene logic*. It has a third logic value UNKNOWN in addition to TRUE and FALSE. In our notation, 0=FALSE, 1=TRUE and 2=UNKNOWN. The UNKNOWN value can be interpreted a state that can be either TRUE or FALSE. When combined with a FALSE value, we know for sure that FALSE ∧ UNKNOWN = FALSE, since both FALSE ∧ FALSE = FALSE and FALSE ∧ TRUE = FALSE. However, TRUE ∧ FALSE = FALSE while TRUE ∧ TRUE = TRUE, and therefore TRUE ∧ UNKNOWN = UNKNOWN.

We employ the three-valued logic system as follows: whenever a cell state has uncertain transitions (i.e. can transition to multiple output states), we assign a value of 2 (UNKNOWN) to it. This also allows us to combine unknown outcomes from different interactions. Any remaining undetermined transitions imply that there are cell states for which multiple transitions are possible.

Concretely, define *g*^(*ij*)^(*X*) ∈ *S* as the outcome of the interaction *i* → *j* for a given input cell states *X*. Note that it takes value in *S*, indicating that the interaction is either on, off or the outcome is unknown. Let 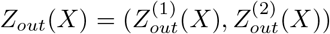 be the three-valued output state given input state *X*. We construct the output state as follows:

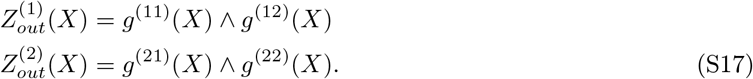

We have replaced the ordinary multiplication of Eq. S9 by the ∧ operation that takes into account unknown outcomes. This three-valued output state needs to be translated to the actual possible output (binary) cell states of the system. Intuitively, if there is an unknown outcome, i.e. 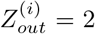 for some *i*, then we should take into account all possible outcomes of that state. Hence we should consider states with both 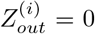 and 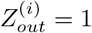.

Formally, let us denote the set of possible output cell states as Σ_*out*_(*X*), with elements in {0, 1}^2^. We construct the set Σ_*out*_(*X*) through the construction of two maps. Let 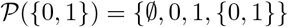 denote the power set of *{*0, 1*}*. First we define a map

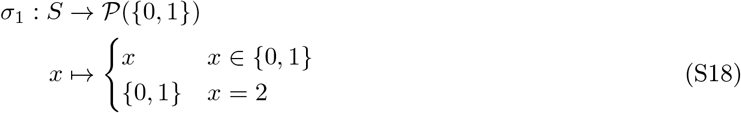

This map constructs the set of possible output gene states, by deconstructing the element 2 ∈ *S* into the set 0, 1 of possible outcomes. Extend the map to *S*^2^ by defining 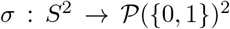 as *σ*(*Z*) = (*σ*_1_(*Z*^(1)^)*, σ*_1_(*Z*^(2)^)).

Next, we have to put together the deconstructions to arrive at a set of output cell states. Recall that 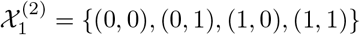 is the phase space of a single cell with two genes. We define a map

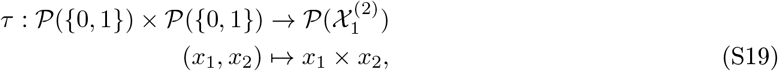

where the × denotes an ordinary Cartesian product between sets. For instance, if *x*_1_ = {0, 1} and *x*_2_ = 1, then *x*_1_ × *x*_2_ = {(0, 1), (1, 1)}. Hence, the second map constructs all the possible cell states from the possible states for each gene. Putting it together, we construct the set of output cell states as

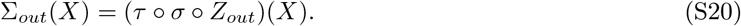

Finally, once we have the output cell states for our input state *X*, we set

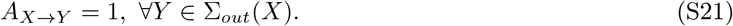

In other words, we draw edges from *X* to all cell states in the set of possible output states Σ_*out*_(*X*). Doing this for all input states *X* gives us the full adjacency matrix for the state diagram.

### S5 Analytic framework for traveling wave propagation

In this section, we provide a detailed analysis of traveling waves moving on a constant background of cells, which are found in Dialogues 15, 19, 33, 34 and 36. We first discuss features of traveling waves that characterize and distinguish different instances of traveling waves (Section S5.1). These features are used in an analytic estimate of the density of traveling wave states in the overall system in Section S5.2. The core of this section is composed of a derivation of a set of conditions for TW propagation (Section S5.3). We then discuss our method for evaluating the performance of the analytic theory in Section S5.5. Finally, we sketch how to extend our method to dynamic patterns on an oscillatory background in Section S5.6.

#### S5.1 Features of traveling waves

The traveling waves that we observe can be distinguished from each other through a number of features:

1. Orientation and direction of the wave. The waves can be oriented in different ways and for a typical topology we observe waves of all different orientations. We distinguish between horizontally, vertically and diagonally oriented waves (see e.g. Figs. 2C, 2D, 2E and 2G). Horizontal waves wrap around the horizontal axis once, without wrapping around the vertical axis, and travel in the vertical direction. Vertical wave wrap around the vertical axis once, without wrapping around the horizontal axis, and travel in the horizontal direction. Diagonal waves wrap around each of the two axes at least once and can travel in either direction. A more precise way of accounting for the geometry of the way is through winding numbers, which will be introduced in section S5.2.
2. Presence of bends in the wave. We distinguish between straight and bent waves according to whether all cells in a band of the wave are aligned in the same direction. For a triangular lattice, there are three directions along which the cells can align themselves. In one case, we get straight horizontal waves (e.g. Fig. 2C), whereas in the other cases we get diagonally oriented waves (e.g. Fig. 2G). However, we can also get waves with one or more bends (e.g. Fig. 2E), points at which the alignment of the cells changes direction. Note that the cells located at the bends have a different set of nearest neighbors from the aligned cells. Furthermore, we can distinguish between bends that are in the direction of propagation (outward bends) and bends that are opposite to the direction of propagation (inward bends).
3. Number of waves. In the simplest case, the system self-organizes into a single wave on a uniform background (e.g. Figs. 2C and 2E). However, we also observe multiple coexisting waves, separated by each other by regions of cells with the background state (see e.g. Figs. 2A and 2D). Such waves have the same orientation and direction of travel, but are not necessarily aligned parallel with each other.
4. Number of different cell states in the wave. For almost all the waves we observed, we found wave made up of three different cell states. The background was made up of the fourth cell states. The exact states which make up the wave and their order varies from topology to topology, and sometimes also between different parameter sets of the same topology. In rare cases, we also found waves consisting of two types of cell states on a background of a third cell state.
5. Number of bands in the wave. In most cases, we find waves consisting of single bands of cells of the same state. Waves with bands with two or more layers of cells and waves where different cell states have different band widths have also been observed (see e.g. Fig. 2C).
6. Defects. In rare cases we may see waves which contain single-cell defects such as an additional cell of the same cell state attached to an otherwise normal wave.

#### S5.2 Abundance of traveling waves

In this section, we provide an estimate of the relative abundance of traveling waves of the forms we observe in the system. Due to the variety of morphologies these waves can take, we could expect them to take up a considerable portion of the total phase space. In this scenario, finding system conditions under which most of the simulations go to traveling waves would not be entirely surprising. On the other hand, if the relative abundance of traveling waves in the system is low, we could interpret this as a sign of a self-organizing mechanism that drives the system towards traveling wave formation.

First, we identify the key aspects of traveling waves and divide them into a limited number of categories. For each category, we then calculate the number of distinct shapes the waves can take, as well as the total number of distinct “snapshots” each wave form is made up of. This then gives us an estimate of the total number of states in the system that can be considered traveling waves.

We have previously provided a list of features that distinguish traveling waves from each other. While we have observed waves that differ in all these categories, we note that the vast majority of waves have the same features for a number of categories. In particular, most waves are composed of three cell states (with a fourth background state), are composed of a single band and have no defects. We also rarely observe more than two waves propagating simultaneously in moderately large systems (e.g. *N* = 256). Hence we only need to consider the orientation and direction, the presence of bends and the number of waves to account for the vast majority of observed wave forms. In the following we consider a generic single-banded wave with *N* = *n*^2^ cells.

The orientation of a wave can be made more precise by considering the number of times the wave wraps around each axis. Since the system is periodic, effectively we are dealing with wave that winds around each axis of a torus different numbers of times. Let *W_x_*, *W_y_* be the winding numbers around the horizontal and vertical axis. For a plane horizontal plane wave such as shown in Fig. 2A, *W_x_* = 1*, W_y_* = 0. For a vertical wave such as in Fig. 2E, *W_x_* = 0*, W_y_* = 1. The diagonal wave in Fig. 2G has *W_x_* = 1*, W_y_* = 2, but we can also imagine diagonal waves that wrap around the system in different ways. The most common winding numbers are listed in Table 2. As is apparent from the table, we mostly observe simple waves that are either horizontal, vertical or diagonal, but wrap around the axes only a few times. Note that traveling waves are characterized by *W_x_* + *W_y_* ≥ 1, i.e. a traveling wave always wraps at least once around one of the axes. (Smaller structures that do not wrap around either axis but do translate in space are referred to as traveling pulses).

**Table 2:** Main properties of most common types of waves. *n* is the linear grid size, with *N* = *n*^2^. We assume that *n* is an even number, so that the system is a perfect hexagonal lattice on a torus. The data is based on empirical observations of self-generated traveling waves. *W*_*x*_, *W*_*y*_ are the winding numbers, *n*_*C*_ is the number of cells of the wave for a given cell state divided by the linear grid size *n*, *N*_*wf*_ is the number of wave forms and *T* is the period of the wave.

| $W_x$ | $W_y$ | $n_C(W_x, W_y)$ | $N_{wf}(W_x, W_y)$ | $T(W_x, W_y)$ |
| --- | --- | --- | --- | --- |
| 1 | 0 | 1 | 1 | $n$ |
| 0 | 1 | 1 | $\begin{pmatrix} n \\ \frac{1}{2}n \end{pmatrix}$ | $n$ |
| 1 | 1 | 3/2 | $\begin{pmatrix} \frac{3}{2}n \\ \frac{1}{2}n \end{pmatrix}$ | $2n$ |
| 1 | 2 | 2 | 1 | $2n$ |
| 2 | 1 | 5/2 | $\begin{pmatrix} \frac{5}{2}n \\ n \end{pmatrix}$ | $2n$ |

Once we fixed the winding numbers, the precise form of the wave is often still unspecified. For instance, a vertical wave can have different number of bends in both directions. Nevertheless, we can derive a general formula for the number of wave forms given (*W*_*x*_, *W*_*y*_). Let us consider a single wave that travels in a fixed direction. Suppose we pick a random cell of the wave. Empirically, we find that each cell of the wave that has the same state has precisely two neighbors with the same state. This is even the case when there are complicated bends in the wave. Now pick one of the neighbors of our selected cell that has the same cell state. The nearest-neighbor vector that connects the two cells lies along one of the six directions one can travel in on a hexagonal lattice. These can expressed in terms of the basis vectors of the lattice as 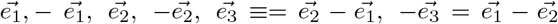 (Fig. S5B). The second cell has a unique neighbor of the same state that we have not selected yet. The vector between the second and third cell defines a new direction that we record. We can therefore continue this procedure and pick subsequent cells in our wave, until we get back to our original cell. This is because the wave wraps around an axis at least once as noted before. For each step we take, we keep track of the direction we need to move in to get to the next cell. At the end, we count the number of steps in each of the six directions. Let us denote these by {*n*_*i,α*_}, where 1 ≤ *i* ≤ 3 and *α* ∈ {−, +}. For example, *n*_2,−_ gives the number of steps we took in the 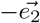 direction.

Wave forms differ by their set of nearest-neighbor vectors that we obtain with this procedure. Nevertheless, once we fix the winding numbers, this constraints the possible sets of direction vectors in a way we can make precise. First, we note that empirically we find that waves with fixed winding numbers always have the same number of cells of a given state, which we will denote *N*_*C*_ (*W*_*x*_, *W*_*y*_) = *n*_*C*_ (*W*_*x*_, *W*_*y*_) *n*. Empirical results for commonly found waves are listed in Table 2. For instance, for a horizontal wave, we find that it always has *N*_*c*_ (1, 0) = *n* cells of a given state, which make up exactly one row of the lattice. This constrains the total number of nearest-neighbor vectors to *N*_*C*_ (*W*_*x*_, *W*_*y*_), such that our first constraint is

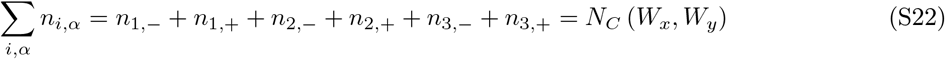

Next, the winding numbers constrain the number of occurrences of each nearest-neighbor vector. For instance, for a horizontal wave the nearest neighbor vectors when added up must be align in the horizontal direction, with a magnitude equal to the grid size. However, a priori this does not imply that all nearest neighbor vectors are in the 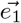 direction, since 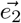 and 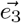 also have horizontal components. In general, the constraints are that the number of steps taken in the horizontal and vertical directions must be equal to the ±*W*_*x*_*n* and ±*W*_*y*_*n* in order to return to the original cell. The sign degeneracy comes from the fact that starting from the initial cell we pick, we can traverse the wave in two different directions, which yield winding numbers that differ by a minus sign. Working out these conditions, we derive the following constraints:

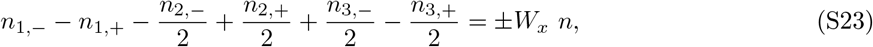

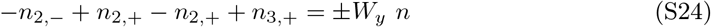

We can now try to solve these constraints together with the general constraints 0 ≤ *n*_*i,α*_ ≤ *n* for all *i, α* for given winding numbers *W*_*x*_, *W*_*y*_. For all the winding numbers listed in Table 2, we obtained solutions of the form 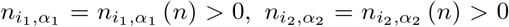 for some *i*_1_, *α*_1_ and *i*_2_, *α*_2_, *n*_*i,α*_ = 0 for all other *i, α*. The interpretation of this result is that in practice all waves are formed by traveling continuously in two directions, i.e. they never “bend back”. Secondly, we found that the number of steps in each direction is a linear function *n*, so we get an explicit scaling of our results with system size.

We can now readily obtain the number of wave forms that satisfy these constraints. This reduces to a simple combinatorics problem where the wave forms differ by the order in which the directions *i*_1_, *α*_1_ and *i*_2_, *α*_2_ appear. In total, there are 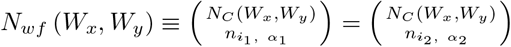 possible wave forms with the given winding numbers. Finally, the sign degeneracy of *W*_*x*_, *W*_*y*_ introduces an additional factor of 2 whenever both *W*_*x*_, *W*_*y*_ > 0. This is because for the four possibilities {(*W*_*x*_, *W*_*y*_), (*W*_*x*_, −*W*_*y*_), (−*W*_*x*_, *W*_*y*_), (−*W*_*x*_, −*W*_*y*_)} only (*W*_*x*_, *W*_*y*_) and (−*W*_*x*_, −*W*_*y*_) give equivalent waves, but (−*W*_*x*_, *W*_*y*_) ≡ (*W*_*x*_, −*W*_*y*_) yields a different wave.

The direction of a wave can in principle be in any of the six directions the hexagonal lattice allows. However, once the orientation of a wave is fixed, there are only two possible directions remaining. This is because a wave always moves perpendicular to the direction in which the cell states of the wave do not change. For instance, for a horizontal wave, the only directions are up and down.

The number of waves that can simultaneously propagate depends on the system size. For *N* = 256, we rarely observe more than two simultaneously propagating waves. Note that the waves need to have the same orientation and direction of motion, or else they would collide and annihilate or form new waves. Once the shapes of both waves are fixed, an additional variable is the spacing between the waves. Assume that both waves are horizontal, then the variable is the number of rows between the waves. For a double wave, the distance between the waves lies in the range 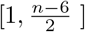, giving a degeneracy of roughly 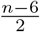. This is because both waves take up 3 rows, and from the distribution of remaining rows we take the shortest distance.

Putting everything together, we now obtain our general estimate for the density of traveling waves in phase space. We estimate this to be in the order of

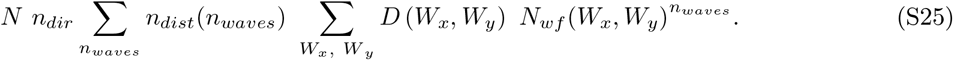

Here *n*_*dir*_ = 2 signifies the two directions of propagation, the first sum is over the total number of waves in the system *n*_*waves*_, the number of unique distances between the waves is denoted *n*_*dist*_, the degeneracy after accounting for negative winding numbers is denoted *D* (*W*_*x*_, *W*_*y*_), and the second sum is over the winding numbers *W*_*x*_, *W*_*y*_ ∈ ℕ_0_. From the previous part we estimate 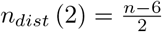. The final term signifies the fact that in case of multiple waves, they can in principle have independent wave forms (with the same direction and winding numbers). The term *N* in front accounts for the possible positions of the wave on the lattice, obtained simply by counting the number of ways to place a given cell of the wave on the lattice. This is an upper bound since in case waves with symmetry different placements of this selected cell could still give the same configuration.

#### S5.3 Traveling wave propagation conditions: nearest neighbor approximation

##### S5.3.1 General conditions for pattern propagation

Before working out the case of traveling waves in detail, let us discuss what pattern propagation means in general. Suppose we have a pattern that periodically repeats itself in time. All information about the pattern is encoded in the states of the pattern over one period. Let us denote these by {*X*(0), *X*(1), … *X*(*τ*)}, where 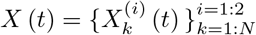 is the state of the system at time t as specified by the states of each of the genes of each cell. These can be considered a series of snapshots of the system that make up a movie of the dynamic pattern when played. In general, take an arbitrary state *ξ*(*t*) and suppose that the system is updated according to a rule

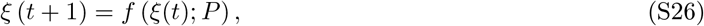

for some unspecified function *f* of the current state of the system that depends on the parameters of the system denoted by *P*. The explicit form of *f* for our model is specified in (Eq. S9–S10). The condition that the pattern can propagate under the set of parameters *P* is precisely that *f* updates each snapshot of the pattern to the next snapshot of the system. In other words, *X* (*t* + 1) = *f*(*X*(*t*); *P*) for all 0 ≤ *t* ≤ τ − 1.

In general, this would put constraints on each of the cells of the system, leading to a convoluted set of conditions. However, in cases where the pattern exhibits a symmetry, these conditions can be drastically simplified. In one extreme case, if the pattern is a homogeneous collection of identical cells, at any time we would only need to check one set of conditions for an arbitrary cell of the system. Conversely, suppose the system is completely anisotropic for the whole duration of the pattern trajectory. Each cell then sees a different environment at any time. We then need to check all *N* × *l* × *τ* conditions for each of the genes of each of the cells of the system, at each time step of the system.

The case of traveling waves allows us to exploit the symmetry of the system to drastically reduce the number of conditions for pattern propagation. First note that traveling waves are characterized by the fact that the state upon updating is related to the previous state by a simple translation in space. This means that rather than checking conditions for each time step, we only need to check that the wave propagates at one arbitrary time step. Furthermore, the spatial symmetry of the system allows us to check only a small number of cells of the system, as will be explained in the next section.

##### S5.3.2 Straight and bent waves

To derive conditions for the propagation of these waves, we look at straight (plane) waves (Fig. 4A, “straight wave”), waves with a single outward bend (Fig. 4A, “bent wave”) and waves with a single inward bend (Fig. 4A, “bent wave” with reverse direction of propagation). In this way, we obtain results applicable to the vast majority of waves observed in the system. More complicated waves such are typically composed of local motifs which are identical to the ones for these three basic types of waves. For instance, the configurations in Fig. 1F contains two waves with multiple bends. However, the nearest neighbors of any cell is identical to the neighbor structure of a cell in one of the three prototype waves. Namely, the cells at the tip of the wavefront have a nearest neighbors that is identical to the cell at the tip of the outward bent single wave. The cells that are bent towards the back of the wavefront have a neighbor that is identical to those at the bend of the inward bent wave.

Therefore, we argue that it suffices to study the conditions for propagation of each of these three simple types of waves. This gives a first approximation to the propagation conditions for more complicated waves. The types of waves which are not covered by this analysis are waves with multiple bands (because the cells of such waves have different nearest neighbors), and waves with defects (which are too rare to motivate analysis of each special case).

##### S5.3.3 Conditions for propagation of traveling waves

###### Wave structure

All waves consisting of three consecutive single bands of cells have a similar spatial structure. For any instance of such a wave, we can identify six types of cells that each have a unique set of nearest neighbors. Let us denote these six types of cells as follows (see Fig. 4B):

1. E_*F*_ – front exterior
2. F – front
3. M – middle
4. B – back
5. E_*B*_ – back exterior
6. E – exterior

Note that the types E, E_*F*_ and E_*B*_ all have the same cell state (the state of the white color in Fig. 2A). However, we divide the exterior cells up into three classes because they have different sets of nearest neighbors. A cell of type *E*_*F*_ in front of the wave neighbors *F* cells, whereas a cell *E*_*F*_ at the back of the wave neighbors *B* cells. Finally, the rest of the *E* cells that border only other *E* cells.

Hence, the six types of cells have four different cell states, which we denote as *X*(*F*), *X*(*M*), *X*(*B*) and *X*(*E*) = *X*(*E*_*F*_) = *X*(*E*_*B*_). For binary cells, the possible cell states form the set *S* = {(0, 0), (0, 1), (1, 0), (1, 1)}.

At any straight segment of a wave, the cells of the wave and those bordering the wave have exactly the same local structure (nearest neighbors). Concretely, this means that any cell of the straight segment borders the same number of cells of each state (Table 1). For instance, an *F* cell will always border two cells with state *X*(*F*), two cells with state *X*(*M*) and two cells with state *X*(*E*).

The neighbor of the cells of the wave differs from that of plane waves only when there is a bend in the wave. In particular, only the cells located precisely at the bend have a different nearest neighbors from the rest of the cells, which have nearest neighbors identical to plane wave cells (Fig. 4B). We must therefore take these into account separately (Table 3).

**Table 3:** Cell states of nearest neighbors of the six types of cells (*E*_*F*_, *F*, *M*, *B*, *E*_*B*_, *E*) for straight waves and for the cells at the tip of waves with bends (Fig. 4B). Results are for a hexagonal lattice with coordination number *z* = 6.

| Plane wave |  |  |  |  |
| --- | --- | --- | --- | --- |
| Wave state | Number of neighbor states |  |  |  |
| | $X(F)$ | $X(M)$ | $X(B)$ | $X(E)$ |
| $E_F$ | 2 | 0 | 0 | 4 |
| $F$ | 2 | 2 | 0 | 2 |
| $M$ | 2 | 2 | 2 | 0 |
| $B$ | 0 | 2 | 2 | 2 |
| $E_B$ | 0 | 0 | 2 | 4 |
| $E$ | 0 | 0 | 0 | 6 |

| Wave with outward bend |  |  |  |  |
| --- | --- | --- | --- | --- |
| Wave state | Number of neighbor states |  |  |  |
| | $X(F)$ | $X(M)$ | $X(B)$ | $X(E)$ |
| $E_F$ | 1 | 0 | 0 | 5 |
| $F$ | 2 | 1 | 0 | 3 |
| $M$ | 3 | 2 | 1 | 0 |
| $B$ | 0 | 3 | 2 | 1 |
| $E_B$ | 0 | 0 | 3 | 3 |
| $E$ | 0 | 0 | 0 | 6 |

| Wave with inward bend |  |  |  |  |
| --- | --- | --- | --- | --- |
| Wave state | Number of neighbor states |  |  |  |
| | $X(F)$ | $X(M)$ | $X(B)$ | $X(E)$ |
| $E_F$ | 3 | 0 | 0 | 3 |
| $F$ | 2 | 3 | 0 | 1 |
| $M$ | 1 | 2 | 3 | 0 |
| $B$ | 0 | 1 | 2 | 3 |
| $E_B$ | 0 | 0 | 1 | 5 |
| $E$ | 0 | 0 | 0 | 6 |

###### Propagation conditions

For the wave to propagate forward, we need a number of conditions to be satisfied. Since traveling waves have the property that the entire structure translates forward by one step, we can easily find these conditions. Basically, all types of cells shift up by one and the background type remains constant. For example, an *E*_*F*_ cell right in front of the wave should become an *F* cell at the next time step. Hence we require that the cell obtains the state *X*(*F*) upon updating. Let *α* → *X*(*α*′) denote the condition that a cell of type *α* acquires state *X*(*α*′) according to the update rule. Then we can succinctly write our set of conditions as:

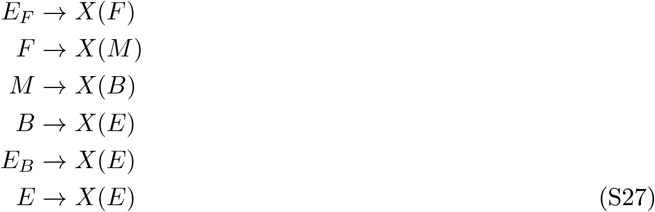

For a straight wave without bends, these conditions need to be checked only once, for cells that have the nearest neighbors as detailed in the first table in Table 3. For waves with at least one bend, both the condition at the location of the bend (either outward or inward) as well as the condition for plane waves (for the straight segments of the wave) need to be checked. For waves with a zig-zag pattern that have no straight segments (e.g. Fig. 2D), only the conditions for inward and outward bends need to be checked.

###### Nearest-neighbor approximation

Given an exact form of the wave and a specific interaction network of the two genes, we can work out the six conditions for traveling wave propagation, to obtain exact conditions in terms of system parameters. However, since the waves can have different features, we look for a more general approach that predicts propagation independent of the precise shape of the wave. To do this, we will apply a nearest-neighbor approximation (NNA). The idea is to only consider the immediate neighbors of a cell when calculating the concentration it senses, and take into account the rest of the cells through averaging and assuming they are randomly distributed.

Write 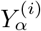 for the concentration of molecule i a cell of type *α* senses. Then we can split the sensed concentration into terms of the cell itself, its neighbors and an approximation of the rest of the lattice:

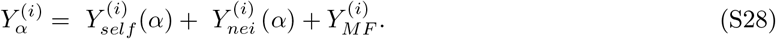

For any cell in the system with state *X*, we have a secretion rate

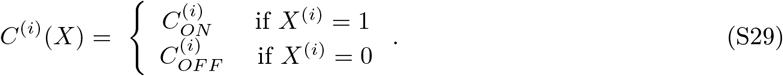

Denote 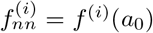 as nearest neighbor interaction strength, and *n*(*X*; *α*) as the number of cells of state *X* that neighbor a cell of type *α*. The sensed concentration due to neighbors can then be written as

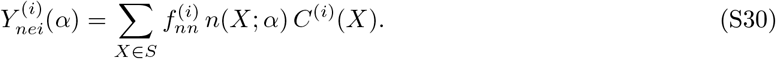

The contribution of rest of the lattice is estimated through a mean-field approximation. For a wave with *N*_*w*_ waves, each consisting of bands of width *W*, with winding numbers *W*_*x*_, *W*_*y*_ (Section S5.2), we can calculate the proportion of cells that have either of the genes on. This proportion depends on the cell states of the wave and background cells. Then the fraction of cells with a given gene on is

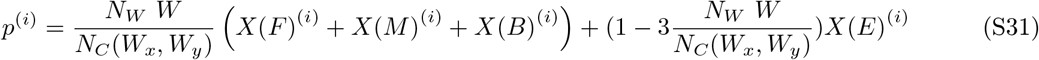

Here *X*(*S*)^(*i*)^ denotes the state of gene *i* of a cell state *X*(*S*). The mean-field contribution is then estimated to be the interaction strength of all cells excluding the nearest neighbors times the average secretion rate of the cells:

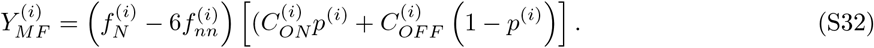

#### S5.4 Explicit example

In this section, we work out an explicit example of the NNA and show that it successfully predicts the propagation of traveling waves. We consider network 15 with interaction matrix 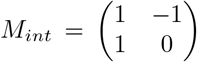, for which we observed traveling waves with the composition (see Section S5.3.3):

- *X*(*F*) = (1, 0),
- *X*(*M*) = (1, 1),
- *X*(*B*) = (0, 1),
- *X*(*E*) = (0, 0).

##### Conditions for propagation

Let us denote *E*_*F*_ = (0, 0)_*F*_ and *E*_*B*_ = (0, 0)_*B*_. Explicitly, the conditions for propagation can be expressed in terms of inequalities (Table 4).

**Table 4:** Explicit propagation conditions in terms of the sensed concentrations and parameters for the example under study. The gene network has interaction matrix 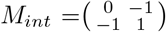. Here, ∧ denotes a logical AND and ∨ denotes a logical OR.

| Condition for propagation | Condition on gene 1 | Condition on gene 2 |
| --- | --- | --- |
| $(0, 0)_F \rightarrow (1, 0)$ | $Y_{(0,0)_F}^{(2)} < K^{(12)} \wedge Y_{(0,0)_F}^{(1)} > K^{(11)}$ | $Y_{(0,0)_F}^{(1)} < K^{(21)}$ |
| $(1, 0) \rightarrow (1, 1)$ | $Y_{(1,0)}^{(2)} < K^{(12)} \wedge Y_{(1,0)}^{(1)} > K^{(11)}$ | $Y_{(1,0)}^{(1)} > K^{(21)}$ |
| $(1, 1) \rightarrow (0, 1)$ | $Y_{(1,1)}^{(2)} > K^{(12)} \vee Y_{(1,1)}^{(1)} < K^{(11)}$ | $Y_{(1,1)}^{(1)} > K^{(21)}$ |
| $(0, 1) \rightarrow (0, 0)$ | $Y_{(0,1)}^{(2)} > K^{(12)} \vee Y_{(0,1)}^{(1)} < K^{(11)}$ | $Y_{(0,1)}^{(1)} < K^{(21)}$ |
| $(0, 0)_B \rightarrow (0, 0)$ | $Y_{(0,0)_B}^{(2)} > K^{(12)} \vee Y_{(0,0)_B}^{(1)} < K^{(11)}$ | $Y_{(0,0)_B}^{(1)} < K^{(21)}$ |
| $(0, 0)_R \rightarrow (0, 0)$ | $Y_{(0,0)_R}^{(2)} > K^{(12)} \vee Y_{(0,0)_R}^{(1)} < K^{(11)}$ | $Y_{(0,0)_R}^{(1)} < K^{(21)}$ |

From this, we see that the last condition for (0, 0)_*R*_ is redundant. Namely, we have 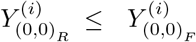 and 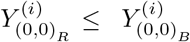. This implies that for gene 2, if the condition for 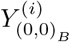 is fulfilled, then that for 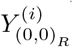 is automatically fulfilled. Likewise, for gene 1, the conditions 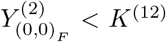 and 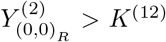 give a contradiction. Since the first condition has to be true, the second is necessarily false. This leaves 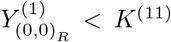. But then this is always true if 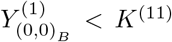 is fulfilled. A side consequence is that 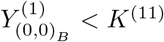 becomes the only condition for gene 1 for (0, 0)_*B*_. Note that this is specific to the network under consideration; leaving out the last constraint is not valid in general.

##### Plane waves

We work out the equations explicitly for plane waves. From Eq. S28-S32, we get the sensed concentrations

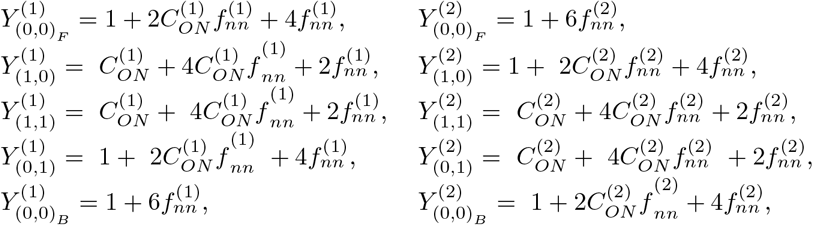

We can then write out the conditions from Table 4 explicitly. For gene 1 this gives:

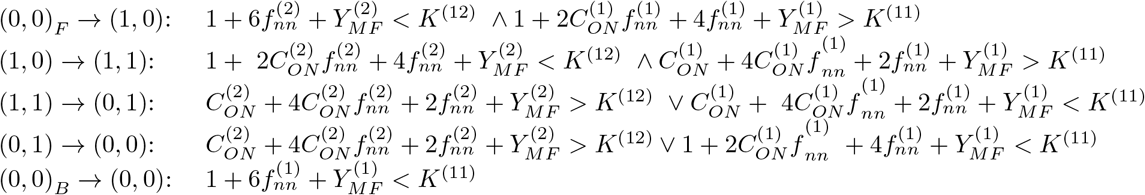

For gene 2, the conditions are

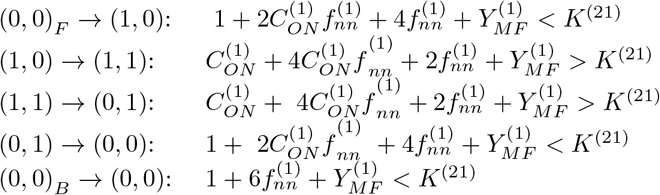

Since 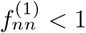 (i.e. the interaction of a cell with itself should be larger than that with its nearest neighbor), we can simplify the equations to account for redundancy. After some algebraic manipulations, this reduces the conditions to the following set of inequalities:

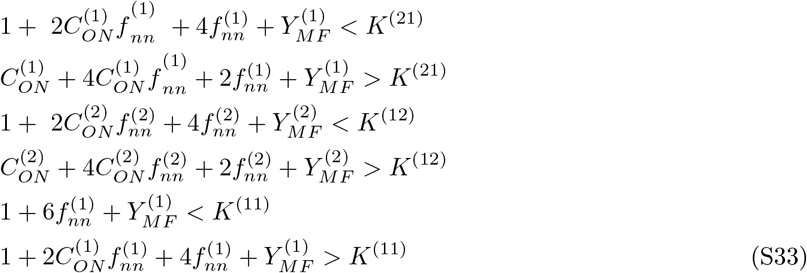

Note that for this particular example, the constraints reduce to a simple set of constraints for each of the three interactions in the system. Namely, the first two inequalities involve only parameters that affect the interaction 2 ← 1, e.g. 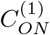 and *K*^(21)^, whereas the second and third pair involve only the interactions 1 ← 2 and 1 ← 1 respectively. This does not need to be the case in general, since genes which are regulated by both genes will produce coupled constraints in terms of both interactions. Hence, we can recast the conditions into a concise set of equations:

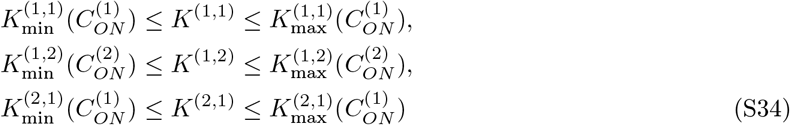

where the 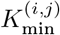 and 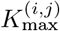 are functions of 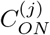 defining the minimal and maximal possible values of *K*^(*i,j*)^.

Explicitly, we have

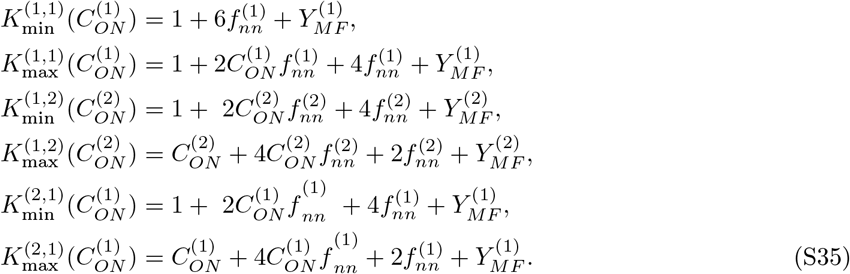

Note also that 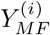 is a linear function of 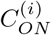 (Eq. S32), so that these constraints reduce to linear relations between 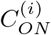 and *K*^(*ij*)^ for each interaction *i* ← *j*. The values 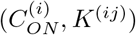 together determine the relative strength of this interaction, and we find that its strength is constrained by two inequalities that determine a reduced but unbounded region of phase space. The boundaries of these regions together with the predicted TW conditions (both true positives and false positives) are plotted in Fig. S3B. This gives an alternative view of the parameter sets that can support TWs from the spider charts, which only show that most parameters can span several orders of magnitude but do not directly reveal the structure of the set of TW parameters. However, this representation reveals that each of the circuit parameters 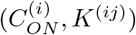 is unbounded and confined to a region which indicates that each interaction can be neither too strong 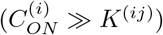 nor too weak 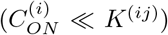, except in the case of the self-activation loop where we tend to have 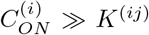. Similar results to Fig. S3B are obtained for the other networks that can support TWs. We also obtain robustness measures from this set of inequalities, as will be discussed in Section S7.

#### S5.5 Performance of the analytic framework

To assess the validity of the analytic framework derived in the previous sections, we directly compared the predictions from the theory to actual simulations of the waves. We quantified the degree to which these results match and considered the accuracy of the main approximation in the analytic framework, the nearest-neighbor approximation (Eqs. S28 and S32).

##### Computational search for traveling waves

We verified with our analytic approach that the above wave forms are indeed the only possible wave forms for two-gene networks. To this end, we screened a large number of parameter sets for all distinct two-gene networks. Specifically, we checked the six conditions Eq. S27 for wave propagation for a total of 10^6^ parameter sets for each network. The parameter sets were generated by Latin hypercube sampling over all non-zero 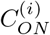 and *K*^(*ij*)^ parameters. We considered a network to be capable of generating a wave if for at least one of the 10^6^ parameter sets all the conditions for traveling wave propagation were fulfilled. The results were consistent among the three types of waves (plane, with inward bends, with outward bends) that we examined: in all three cases exactly the same results were found.

##### Statistical measures for performance

The performance of the analytic method we derived is determined by how well it predicts the conditions under which traveling waves can propagate. We can view NNA approach as a binary classifier that predicts for a given gene network and given set of parameters whether TWs can propagate. The theory takes as input a set of parameters and gives as output a binary prediction about whether the TW can propagate or not. As such, we quantify its performance using well-established concepts for evaluating classifiers from machine learning. In particular, we look at the *accuracy*, *precision* and *recall* of the predictor for all the six cellular dialogues and corresponding waveforms we found. These are defined as

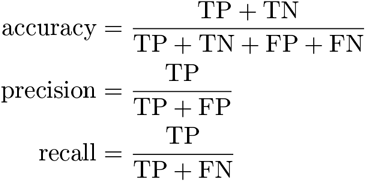

Here TP = true positives, FP = false positives, FN = false negatives. True positives are parameter sets for which the TW propagates according to both theory and simulation. False positives are predicted to be capable of sustaining TWs by the theory, but turn out not to do so in an actual simulation. False negatives are parameter sets that are capable of propagating TWs, but are missed by the theory.

##### Interpretation of the performance metrics (Fig. S3)

For all of the networks that yielded waves, we find that the theory correctly predicts plane TWs to an extremely high degree of accuracy, close to 100% (Fig. S3A). This means that the theory correctly predicts whether a wave can or cannot propagate in almost all cases. In contrast, the precision and recall take slightly lower scores, with a precision is between roughly 0.6-0.8 and a recall in the range of roughly 0.5-0.7. The interpretation is that roughly 60-80% of conditions predicted to allow TW propagation are indeed ones that can propagate a TW in an exact simulation, and that 50-70% of the all the conditions for TW propagation are correctly identified by the classifier. These lower values are caused by the low number of actual positives (conditions under which a TW can propagate) is low compared to the total number of parameters we examined, as we discussed before in the context of robustness of TW propagation. This means that the few incorrect predictions that arise from the approximation have a relatively large impact on these performance metrics. Nevertheless, we find that the theory is still extremely accurate in most cases and qualitative predictions about which networks can yield TWs and orders of magnitude for the corresponding parameters should be highly accurate. The successes and failures of the analytic theory also become apparent when we plot the interaction parameters of the predicted and actual waves together with the theoretical bounds (Fig. S3B). This shows that the false predictions (false positives and false negatives) are mainly due to slight misestimations of the boundaries of the regions allowing for TW propagation. In particular, both the upper bound and lower bound for *K*^(*ij*)^ are slightly underestimated, meaning that the estimations for the mean-field contribution 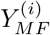 are underestimated. This is evident from the fact that most false positives are near the lower bound for *K*^(*ij*)^ and most false negatives are near the upper bound for *K*^(*ij*)^.

##### Validity of the nearest neighbor approximation

The accuracy of the nearest neighbor approximation depends on how much of the total interaction the nearest neighbors capture. The more the nearest neighbors contribute to the total interaction strength, the more accurate the approximation is. This is because all deviations from the exact model come from the mean-field approximation, which has only a marginal contribution if the sensed concentration is mostly due to the cell itself and its nearest neighbors. We can quantify this by comparing 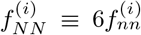 (since there are six nearest neighbors) to the total interaction strength 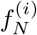 for signaling molecule *i*. If 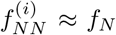, then the cells beyond the direct neighbors have only marginal influence on the concentration a cell senses. However, if 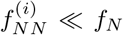, then the nearest neighbor approximation will be comparatively inaccurate, because we take into account the rest of the cells in an averaged manner only and neglect their spatial positions.

Note that 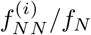 depends on the parameters *N*, *a*_0_ and *λ*^(*i*)^. By examining how this quantity depends on these parameters, we get a picture of when the NNA is most accurate. For weak interaction (high *a*_0_), the nearest neighbor approximation matches closely with the actual system. For stronger interaction, the nearest neighbor approximation becomes worse (Fig. S3C). In this case, one possible solution would be to extend the analysis to next-to-nearest neighbors. When their influence is taken into account, the 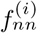 becomes considerably closer to *f*_*N*_. In contrast, the ratio is hardly dependent on system size *N* and diffusion length *λ*^(*i*)^ (Fig. S3C). Finally, longer diffusion length implies comparatively more influence from cells further away, leading to less accuracy for the nearest-neighbor approximation. However, this effect is also weak, accounting for less than 30% variation in interaction strength.

We also considered how taking next-to-nearest neighbors into account improved the accuracy of the NNA. As such, we defined an interaction parameter 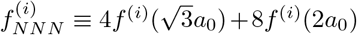, which takes into account the interaction with the t*√*welve cells in the second layer surrounding a cell on a hexagonal lattice. There are four cells at a distance of 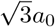 and eight cells at a distance of 2*a*_0_ in this layer. We find that total contribution to the interaction strength from the two layers of cells closest to a given cell, 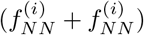 indeed captures a significantly larger portion of the interaction strength. Thus, by extending the analytic theory to take into account next-to-nearest neighbors, we could improve on the approximation. However, based on the performance metrics of the NNA we found, we would argue that such an extension is unnecessary since a NNA theory already gives accurate predictions.

#### S5.6 Oscillatory traveling waves

So far, we focused on traveling waves that are characterized by a propagating pattern on a fixed background. However, we also observed a variety of dynamic spatial patterns with oscillatory background cells in Networks 16, 20 and 43(Fig. S1). In particular, oscillatory traveling waves (Sec. S3.3) form a subset that we can analyze using our framework. Here, we outline how to adapt our framework to the analysis of these dynamic patterns. From simulations, we observe that the oscillations have period 3 and always follows a fixed pattern. Using the definition of the Exterior (E), Front (F), Middle (M) and Back (B) states of a wave (S5.3.3 and Fig. 4), we can trace out how these states transition on a state diagram (Section S4). We found that the cells of oscillatory waves follow a fixed pattern of cell state transitions (Fig. S1D). Networks 16 and 20 each have one distinct state diagram, and network 43 can generate waves that follow either of the two state diagrams. Each of the states undergoes a separate period 3 oscillation, but together they follow an regular pattern on the state diagram. At the transition with the dotted lines, the wave moves one step further (i.e., the entire pattern not only oscillates but also moves one cell layer ahead). This always occur for transitions where both genes are switches: either between (0, 0) and (1, 1) or between (0, 1) and (1, 0). Waves in network 43 can follow either the transitions of network 16 or those of network 20, or show more complicated patterns which we will not go into here. Hence, by imposing each of the transitions on either the same group of cells (when the wave oscillates but not moves) or a neighboring group of cells (when the wave translates), we could in principle derive a more complicated set of constraints for the propagation of such waves.

### S6 Extensions of the model

In this section, we discuss how we extended our model to include more complex elements of communicating multicellular systems. The effect of the five extensions on the formation and propagation of dynamic spatial patterns is discussed in the main text and Figs. 6 and S6.

#### S6.1 Stochastic sensing and response

There are various sources of stochasticity and variability in the system that could play a role in multicellular patterning. These include cell-to-cell variability, stochastic gene expression and fundamental limits in sensing accuracy. We did not try to derive exact expressions for each source of noise, but rather work with phenomenological terms that lump together various stochastic effects. In particular, we extend the noise description of [2] to two genes by allowing for fluctuations in the threshold of each interaction. We consider a stochastically fluctuating threshold concentration obtained through the addition of a noise term:

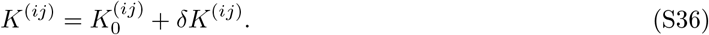

Here 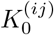 is the threshold concentration for the interaction *i* ← *j* in the absence of noise and 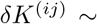 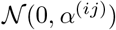 is a normally distributed random variable. At each time step, we apply Eq. S36 to update the threshold independently for each interaction *i* ← *j* and independently for each cell. In order to define a global noise strength without introducing many variables, we take 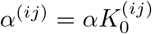. In other words, 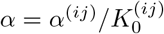 is fixed for all interactions, meaning the variation in the threshold is scaled by the same factor *α* for each interaction.

#### S6.2 Continuous cell response function

In the model we assumed that cells are binary and secrete signaling molecule *i* at either a low rate 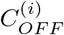 or a low rate 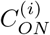. This is a valid assumption whenever the response function is sufficiently ultrasensitive, as discussed in the main text. However, to take into account more gradual response functions, we replace the step-function response S10 by a Hill function. The steepness of the Hill function is characterized by the Hill coefficient. For simplicity, assume that all molecules have the same Hill coefficient *n*. The update rule for the cells’ states is still given by Eq. S9, but now with

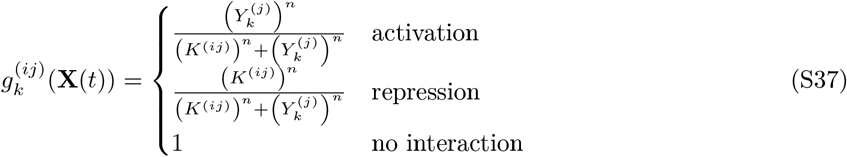

Note that the Hill coefficient in our model does not have a direct physical interpretation, but is a phenomenological parameter describing the steepness of the response. This is because in real systems, the sensed molecule induces an effect on the regulated gene typically through a complex signal transduction pathway rather than a single regulatory step. As such, the Hill coefficient does not relate to the cooperative binding of any specific biochemical process. Therefore, whereas in cooperative binding models values of *n* < 1 or *n* > 2 are rare, in our model the Hill coefficient can in principle be arbitrarily high or low.

#### S6.3 Disordered cell positions

We have so far considered cells that are positioned on a perfect hexagonal lattice. This is reasonably accurate for certain multicellular systems (see Table S1 in [2] for a list of examples), but in general communicating cells do not need to be arranged on a regular lattice. For instance, cell culture experiments are often done by streaking a liquid colony of for instance yeast cells onto a plate. The cells that appear in these colonies are usually not regularly arranged on a grid. Collective behavior in unicellular quorum sensing organisms tend to have different spatial configurations. To extend our model to take into account alternative spatial arrangements, we adapted our model to allow for randomization of the cell positions through an algorithm adapted from Markov Chain Monte Carlo (MCMC) simulations of hard spheres [7]. The algorithm allows us to tune the degree of randomness of the cell positions, varying from a perfect lattice to a fully disordered placement of cells. However, we still assume that the cells are immobile or move at a much slower time scale than their gene expression dynamics.

##### Randomization algorithm

We model the cells as 2D hard spheres with a radius of *R*_cell_ (identical for all cells). The cells should be placed in such a way that no two cells overlap. Initially, the cells are placed on a regular hexagonal lattice, with distance *a*_0_ between the cells. We select a random cell *j* with position 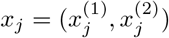. We then perform a Monte Carlo step, where we attempt to move the cell by a displacement *x*_*j*_ → *x*_*j*_ + *δx*. Here *δx* = (*δx*^(1)^, *δx*^(2)^), with random numbers *δx*^(1)^, *δx*^(2)^ independently drawn from a *uniform* distribution on [−*ε, ε*]. If the cell does not overlap with any other cell at the new position, the move is accepted. Otherwise, the move is rejected and a new move is a proposed. To avoid repeated rejections, the cell radius and *ε* are chosen to be sufficiently small. In all our simulations we took *R*_cell_ = 0.2*a*_0_ and *ε* = (*a*_0_ − 2*R*_cell_)/4 = 0.15*a*_0_.

The number of Monte Carlo steps we perform using this algorithm is a measure for the degree of randomness in our cells’ positions. As a rough indication, for a system of *N* = 12 × 12 cells, after 100 MC steps the arrangement still looks very similar to a lattice with slight perturbations. After 10^4^ Monte Carlo steps the cells are clearly not on a lattice anymore, but distinct rows and columns of cells are still recognizable. After 10^5^ Monte Carlo steps the arrangement looks identical to that obtained by randomly seeding the cells in the 2D space. We can make these statements more precise by looking at the spatial distribution of cells surrounding each cell. Quantitatively, we now have a different interaction strength (Eq. S6) for each cell in the system. As the cells become more randomly arranged, the distribution of the interaction strengths becomes broader and the mean also increases. From these calculations, it can be shown that after roughly 10^5^ Monte Carlo steps our system of *N* = 144 cells obtains spatial structure that is indistinguishable from that of a system of randomly placed cells (detailed results not shown).

#### S6.4 Cell motility

We also extend our model to account for (undirected) movement of the cells. Cell motility has been proposed to be a stochastic process that can be modeled by a Langevin equation. This approach has been used for a variety of systems, including human chick heart fibroblasts [8], endothelial cells [9] and human granulocytes [10]. More precisely, these papers propose that the underlying process is that of an Ornstein-Uhlenbeck process, whereby the cells drift randomly but experience a restorative force (corresponding to friction in a Brownian motion process), that tends to bring cells back to their original position. The discrete time process corresponding to this is obtained through a non-trivial derivation and is shown to be of the following kind [8]:

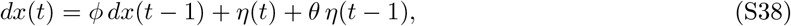

where *dx*(*t*) = *x*(*t*) − *x*(*t* − 1) is the displacement of a cell at time *t*, *η*(*t*) is a discrete random noise term with mean zero and *ϕ, θ* are real numbers that depend on the strength of the restorative force.

For our system, we take a simpler approach to modeling cell motility that neglects the time correlations arising from the frictional term. Instead, we assume the cells drift around in an uncorrelated manner, corresponding to a classic random walk or Wiener process. To model this process, we use the same Monte Carlo algorithm we used for randomizing lattice cell positions (Section S6.3), but now move the cells at each time step instead of for the initial state. We define the *cell motility σ*_*D*_ to be the width of the Gaussian term, in units of *a*_0_, describing the Brownian motion process through which we update the cell positions. Explicitly, at each time step we update the cells one by one through

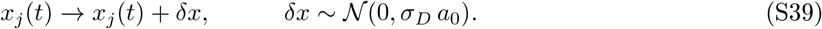

Here *σ*_*D*_ is a parameter characterizing the extent of the motion in units of the lattice constant *a*_0_ (the distance between two neighboring cells when placed on a lattice).

#### S6.5 Parameter gradient

It has been shown that gradients in production rates, model parameters and anisotropy can influence the orientation of stripe patterns in Turing systems [11]. Analogously, we wondered whether parameter gradients could influence traveling waves in our system, in particular whether they could exert an influence on the orientation of the waves formed. To this end, we experimented with applying parameter gradients in various directions and for various parameters. Starting from a parameter set which is able to generate waves, we modify one of the parameters *P* of a cell *k* to be space-dependent,

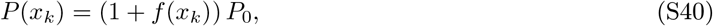

where 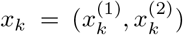 is the position of cell *k* and *f*(*x*_*k*_) is a modulation term and *P*_0_ is a constant. The simplest type of gradient we can take is a step function defined in the horizontal or vertical direction (Fig. S8C). For instance, we could take a vertical gradient by choosing 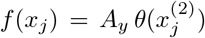, with 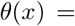 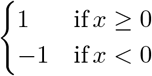 (assume that half of the cells are at *x*^(2)^ > 0). The sharpness of the gradient is then quantified through a gradient strength parameter *A*_*y*_, which represents the fractional change in the parameter value at either side of the step. Note that with this gradient, the average value of the parameter remains constant over the lattice, i.e. Σ*f*(*x*_*j*_) = 0.

### S7 Robustness and reliability of traveling waves

#### S7.1 Robustness (Fig. S7A-C)

Biological robustness is typically referred to as the ability of a biological system to adapt to environmental perturbations by maintaining its function [12]. Control mechanisms such as feedback loops may play a role in maintaining robustness. In our system, different time scales allow us to study the concept of robustness at different levels. The dynamics of the parameters of the system occurs at an evolutionary time scale (unless the experimentalist intervenes), while the dynamics of the gene expression happens on a much shorter time scale (minutes to hours) and the dynamics of the signaling factors occurs on an even faster time scale. As such, we may consider perturbations at each of these levels of description to see how they affect the system’s ability to perform a certain function - which in this case means it’s ability to generate patterns. In this paper, we considered the system’s response to changes in parameter values. Since we assume the parameters to stay constant during the entire simulation, we will use robustness as a static quantity obtained by comparing simulations at different sets of fixed parameters.

Specifically, we considered the robustness of traveling waves for two different situations. We considered the robustness of TW formation - how changing parameters impacts the system’s ability to self-organize into a TW, as well as the robustness of TW propagation - how parameters influence the ability of an already formed TW to continue propagating. We quantified the robustness in both cases by the fraction of parameter sets, or Q-value, that can generate or propagate a TW [13, 14]. In the absence of further information about the parameters, this tells us how likely it is to find parameters which are compatible with a certain property or behavior of the system - formation or propagation of TWs in our case.

##### Normalized Q-value

The Q-values we obtain as a fraction of parameter set compatible with TWs depend on the number of parameters *m* we sample over for each network. These values will tend to be higher for networks with fewer parameters than for networks with higher number of parameters. One method used to correct for this is to take the m-th root of the Q-value [13]. We will call this the “normalized Q-value”. This value represents the chance for each of the *m* parameters to be compatible with TW formation over a specified range of values.

##### Calculation of Q-values

In the absence of any predictive theory, the phase space volume compatible with TWs can only be estimated through drawing random samples from the parameter space and determining whether TWs can form or propagate for each of these samples. Note that in principle our parameters are unbound, i.e. *K*^(*ij*)^ ≥ 1 and 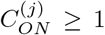 with no upper bound. To calculate the robustness, we therefore specified a finite region defined by 1 ≤ *K*^(*ij*)^ ≤ *L* and 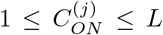 for each signaling molecule *j* and each interaction *i* ← *j* (neglect the parameters for non-existent interactions). In practice, we took *L* = 1000 everywhere. We then used Latin hypercube sampling to generate a large number of parameter sets and tested whether TWs could form or propagate for each of the parameter sets.

For TW formation, we tested how likely it is to find self-organization of TWs in the five networks for which we found self-organized TWs (Fig. 3D). We used the same 10,000 sampled parameter sets as were used to generate the initial network classification (Figs. 3 and S2). For each parameter set, we considered it to be capable of self-organizing TWs if at least one simulation (out of 10 runs per parameter set) led to a self-organized TW. The Q-value we obtained for TW formation in this way is of the order of 10^−3^ (Fig. S10A). Alternatively, after correcting for the number of parameters, the normalized Q-value corresponds to a randomly generated parameter having around 30% chance of taking a value compatible with TW formation across a 1,000-fold range for each parameter (Fig. S10A). The Q-values obtained in this way are in fact lower estimates as we perform only a finite number of simulations and would be higher if we could screen over all possible initial states (in which case the Q-values for TW formation and TW propagation would coincide, since we would also include the final pattern as initial state). Nevertheless, this approach mirrors the situation in wet lab experiments, where can only test a finite number of replicates before concluding that a particular result is highly unlikely to be reached.

For TW propagation, we tested the two types of TWs we found (Fig. 4D) for each of the networks in which we found them. We used the same data as obtained from Latin hypercube sampling which we used to quantify the performance of our analytic predictor (Fig. S3), as this contains precise information on whether each parameter set should be able to propagate TWs according to both the theoretical prediction as well as explicit simulations. This gave higher Q-values, in the order of 10^−2^ for TW propagation (Fig. S7B), corresponding to normalized Q-values of about 40−50% for each parameter to be compatible (Fig. S7A). In comparison, the robustness of the *Drosophila* segment polarity gene network was quantified for a network with a far larger set of parameters, for which the authors found Q-values corresponding to normalized values of about 80 − 90% for each parameter [13].

#### S7.2 Reliability (Figs. 5F and S8)

While robustness deals with the probability of finding parameter sets compatible with TWs, we can also ask what the chance of finding a TW is once the parameter set has been fixed. We define the *reliability* of TW formation as the percentage of simulations with varying initial conditions that generate TWs given a set fixed parameters. For each of the sets of parameters that yielded self-organized TWs, we determined the reliability by running a large set of simulations and counting in how many of those TWs spontaneously formed. Overall, we found an average reliability of 0.2-0.4 across all networks, indicating that we expect TWs to form in roughly 20-40% of the time for these parameters sets (Fig. 5F). However, upon closer inspection we find that this average results from a considerable variability between different parameter sets, indicating that the precise choice of parameters has a large influence on the reliability of TW formation (Fig. S8A). While for many parameter sets the reliability is exceedingly low (5-10%), there is a continuum of reliability values all the way up to about 80% (Fig. S8A).

This finding raises the question of whether we could identify any source of this variability in reliability values between different parameter sets. To address this question, we took a large set of parameter sets (*n* = 2534) capable of sustaining a TW once it has formed, as tested explicitly in simulations starting with a TW. For each of these parameter set, we then ran a large number of simulations (100) to see whether it could also self-organize into traveling waves if we set up the initial configuration to be random. Surprisingly, we found that a large set of these parameter sets did not yield self-organized waves at all (Fig. S8B). This indicates the system may be able to propagate a pattern, but have only few ways of generating such a pattern in the first place. Between the parameter sets that were found to self-organized TWs, the reliability varies dramatically along a continuum between 0 - virtually no simulations become TWs - to close to 1 - almost all simulations become TWs. When we plot the distribution of the 578 parameter sets found to generate TWs, we find that the probability to find a given reliability decays nearly monotonically (Fig. S8C), indicating that parameter sets with higher reliability increasingly rare. However, when we examined the reliability of these different parameter sets as a function of the parameters, we observed no clear trend or correlation in two different projections of the parameter sets (Fig. S8D-E). The parameters sets with extremely high reliability values are scattered around the entire region in which TWs are possible (Fig. S8D). Furthermore, for any of the parameters, there are parameter sets with high reliability for both very high and very low values of that parameters, and the same applies to low reliability (Fig. S8E).

#### S7.3 Influence of initial conditions (Fig. S7D)

In our deterministic model, the initial state of the system fully determines whether a TW forms or not, even though the link between initial and final state is typically unclear unless one runs an actual simulation. It is clear that not all initial states lead to TWs even when suitable parameters are chosen. As a counterexample, consider initiating the system as a uniform lattice of cells that each have the same state. Since all cells sense the same concentration, each cell will evolve to the same state. More generally, in a deterministic cellular automaton there is no symmetry breaking mechanism that can produce a pattern with a set of symmetries from an initial state with a different set of symmetries.

We therefore studied whether certain features of the initial states had a significant impact on whether TW formed or not. In particular, we looked at the contribution of a few statistical variables characterizing the initial states - the initial mean expression level of the genes (*p*^(1)^, *p*^(2)^) (Eq. S11) and the initial spatial order of both genes (*I*^(1)^, *I*^(2)^) (Eq. S12). This is because even for moderately large systems, the number of states exceeds the computational limits of ordinary computers (e.g. for a lattice with *N* = 100 cells, there are 4^*N*^ ≈ 10^60^ states), making it impossible to exhaustively simulate all initial states. We found that the fractions of cells with either of the genes ON had a significant impact on whether TWs formed or not, while the initial spatial order had a notable but much smaller impact.

##### Initial fractions of active genes (Fig. S7D - left plot)

We studied the influence of the initial fractions of ON-cells for both genes by generating states with fixed values of (*p*^(1)^, *p*^(2)^) - this can be easily done by randomly drawing a fixed number of cells to be ON for both genes. Our first observation is that no TWs form for extreme values of *p*^(1)^, *p*^(2)^. For instance, if we start with very low initial *p* values we cannot produce a TW. Both fractions of ON-genes will go to zero and the system will go to the homogeneous state. In contrast, the highest probabilities to produce a TW was found for intermediate values of *p*^(1)^, *p*^(2)^, and can reach a maximum of more than 80%. This is remarkable, since it implies that for the given circuit is capable of generating TWs almost certainly if one initiates the system with the given parameters and fractions of genes that are ON, regardless of how the cells that have these genes ON are placed in space. In particular, it suggests that one may improve the maximal reliability of the system (previous part and Fig. S8) by placing constraints on the initial states.

These findings were confirmed in simulations using other parameters sets capable of generating TWs (not shown). While the exact numbers differ between parameter sets, we consistently observed that more moderate levels of *p*^(1)^, *p*^(2)^ close to (0.5, 0.5) had higher probabilities to generate TWs.

##### Initial spatial order (Fig. S7D - right plot)

In a similar vein, we studied the effect of the initial amount of spatial clustering on TW formation by running simulations where the initial *I*^(1)^ and *I*^(2)^ were varied (using the approach described in Section S2). We observed highly similar results across the range of initial *I* values, but found that fraction of TWs formed tended to be lower at lower values of *I*.

In order to draw more statistically rigorous conclusion about the results displayed in Fig. S7D (right plot), we fitted a logistic regression model to the simulation data using the statistical programming language R. A logistic regression model uses one or multiple predictor variables to calculate a probability that a sample belongs to one out of two possible classes. In this case we can use the initial values of *I*^(1)^ and *I*^(2)^ to predict if a TW forms (classes: TW, no TW).

First, we determined that the logistic regression model using the information of both *I* values (the proposed model) was significantly better than the null model, which takes uses only information about the values of (*p*^(1)^, *p*^(2)^). This indicates that there is information in the *I* values, and knowing the initial *I* has a non-negligible influence on whether TWs form or not. However, the *residual deviance* (indication of goodness of fit of the model based on the log-likelihood of the data given the model) is hardly reduced when going from the simple null model to the more complicated proposed model. This means that including the initial *I* only marginally improves the quality of the predictor.

Next, the proposed model is used as a classifier. For each of the simulations the logistic model uses the initial *I*-values to predict whether the simulation will result in a TW or not. More specifically, the logistic model gives a probability that initial *I* values (*I*^(1)^ and *I*^(2)^) will result in a TW. If this probability exceeds 0.5 a TW is predicted.

The classifier always predicted a wave, except for the relatively low *I* values that correspond to the lower wave fractions observed in simulations. The overall performance of the classifier was further assessed and the results are in Table 5, using the same metrics as used in S5.5. The performance metrics indicate that the logistic classifier performed slightly better than random (accuracy > 0.5). Most TW were correctly predicted (high recall), but this is only because the classifier mostly predicts waves. Many simulations that did not yield a TW were incorrectly labeled as TW (low specificity). It should be noted that the classifier was used to predict the data it was trained on, leading in general to overestimation of the performance metrics. Altogether, this implies that the influence of initial spatial order on the formation of TWs is only marginal.

**Table 5:** Performance metrics of the logistic model fitted to the data shown in the right plot of Figure S7D. The logistic model is used as classifier on the same data as it was fitted on.

| Metric | Accuracy | Precision | Recall |
| --- | --- | --- | --- |
| Value | 0.562 | 0.565 | 0.835 |

